# Associative memory formation reshapes the learning trajectory of a novel cognitive map

**DOI:** 10.64898/2026.09.07.749796

**Authors:** Nathalie Immerzeel, Cecilia Gallego-Carracedo, Lorenzo Mauro, Catalin Mitelut, Flavio Donato

## Abstract

Learning never occurs on a blank slate, as each experience an animal encounters in its daily life is encoded by circuits already shaped by previous ones. During such experiences, the hippocampus builds place-cell-based cognitive maps of the environments explored, and encodes ensemble-based memories for associations among events that occur within them. These spatial and mnemonic representations draw on neurons embedded within the same network. However, whether forming an associative memory of an experience unfolding in one context changes how the same hippocampal network maps a different one is unknown. Here, we longitudinally imaged CA3 neurons as mice familiarized with one environment, encoded either an associative fear memory or a context-only memory in a different environment, and then explored a novel one. Across separate experiences, neuronal recruitment into cognitive maps and memory ensembles followed structured, partially orthogonal allocation rules that were consistent across fear and neutral memories, with ensemble membership and place-cell identity showing no reciprocal enrichment. Fear memory encoding left the properties of the familiar map unchanged, while altering those of the novel one. From first exposure, the novel map already recruited a familiar-like fraction of place cells and supported spatial decoding with near-familiar precision, and its population activity occupied a latent subspace that aligned with the familiar map while remaining context-specific. Yet this novel map failed to undergo the experience-dependent refinement of decoding precision and across-day latent alignment that normally accompanies familiarization, even though place cells continued to gain spatial information and to remap. Associative memory encoding therefore shifts CA3 into a distinct representational regime, decoupling the immediate precision of a new cognitive map from the experience-dependent refinement of its context-specific population embedding.

## Introduction

During everyday life, animals learn continuously from experiences that accumulate one after another, and every new experience is processed by circuits that earlier ones have already modified. Learning does not occurs on a blank slate: an animal might create a memory of an event that happened in one context, navigate an unfamiliar environment moments later, and return to a well-known one soon after. During these experiences, the hippocampus builds cognitive maps of the explored space through the activity of spatially tuned neurons, the place cells^1,2^, and encodes memories for associations among events that occur within these contexts through the activity of distributed neuronal ensembles that are activated during memory encoding, and reactivated during its retrieval (memory ensembles)^3–19^. These two forms of hippocampal representations are increasingly understood as complementary expressions of a common coding system that organizes experience relationally across places, events, and contexts^15,16,20–23^. If cognitive maps and memory ensembles are generated within the same network, then a representation formed during one experience may condition how the next one is built. Yet, whether and how the encoding of an associative memory for an experience unfolding in a context shapes how the hippocampus represents an experience that follows in a different one remains poorly understood, in part because answering these questions requires tracking the same hippocampal population across a prolonged succession of learning episodes across multiple contexts.

When an animal enters a new environment, individual neurons rapidly acquire spatial tuning, allowing a map of space to emerge early during exploration^24–26^. This initial representation is then progressively refined over repeated exposures: population decoding of position becomes more accurate^27^, the proportion of place-cells and their spatial information increases^24,25,28,29^, and population activity stabilizes, with a slower degree of drift between visits to the context^30–34^. The formation of a cognitive map therefore comprises an initial organization of a spatial representation and its subsequent refinement into a more precise and stable population code. Whether the encoding of an associative memory might influence these two steps, and whether it influences them to the same extent, remains unknown. In principle, such influences might arise through two complementary routes. The first is through shared neurons that participate in both representations. Memory ensembles are not allocated randomly: neuronal excitability, prior activity, local competition, and inhibitory constraints influence which neurons are recruited^19,35–38^, and several of these same factors also shape which neurons acquire spatial tuning^39–41^. Consistent with this shared logic, memory ensembles and place-cell populations can overlap, although their relationship varies with experience: in novel environments ensemble membership and stable spatial tuning across episodes can dissociate^15^, whereas in familiar environments they align more closely^21^. The second route is through the surrounding network. Memory encoding alters excitability and plasticity beyond the neurons recruited to memory ensemble, extending to inhibitory neurons that undergo experience-dependent plasticity after learning^4,42–49^. At the same time, interneuron networks regulate place-cell recruitment and tuning^40,50–53^. Thus, a mnemonic representation may affect future spatial coding not only by sharing neurons with cognitive maps, but by reshaping the population-level state in which a subsequent map is initialized and refined.

Within the hippocampal formation, CA3 is a circuit in which such influences are especially plausible. Its recurrent collateral architecture supports auto-associative dynamics and the storage of multiple representations within a shared network^9,20,54–58^. Recurrent networks of this kind are moreover intrinsically sensitive to recent activity in their dynamics, inhibition, and plasticity^54,55,57,59–64^. Whether one representation constrains the next should therefore be visible not only in which neurons are recruited, but in the low-dimensional dynamics of the population as a whole^65–74^.

Here, we longitudinally imaged CA3 population activity as animals first formed a stable map of a familiar environment, then acquired either an associative fear memory or a neutral context experience in a distinct training context, and finally encoded and repeatedly re-experienced a novel environment. This interleaved design allowed us to compare spatial and mnemonic representations within the same tracked neuronal population, and to examine their interaction at two complementary levels: the recruitment, activity, and tuning of individual neurons, and the organization of spatial information within population dynamics. We find that spatial and mnemonic representations recruit overlapping CA3 neurons but remain governed by structured, largely segregated recruitment rules. Fear memory formation does not overwrite this architecture or disrupt the familiar map. Instead, it changes the trajectory of new map learning: a novel environment is encoded with familiar-like precision from its first exposure, but fails to undergo the experience-dependent refinement of its population-level embedding that normally accompanies familiarization. These findings reveal that associative memory encoding reshapes future spatial coding by decoupling immediate map precision from the experience-dependent refinement of a context-specific population representation.

### Cognitive maps and memory ensembles converge on overlapping but structurally segregated subnetworks

To investigate how mnemonic and spatial representations interact with each other in the hippocampal area CA3, we designed a longitudinal paradigm in which activity from the same neurons could be recorded across interleaved episodes of spontaneous foraging and associative memory acquisition and recall (Figure 1a-c). Using head-mounted miniature microscopes, we recorded calcium activity from large populations of CA3 neurons in freely behaving mice over weeks (Figure 1a, b), which allowed us to follow individual neurons as animals acquired either a context-specific aversive memory or experienced the same context without aversive reinforcement, while simultaneously forming cognitive maps of multiple environments.

**Figure 1:**
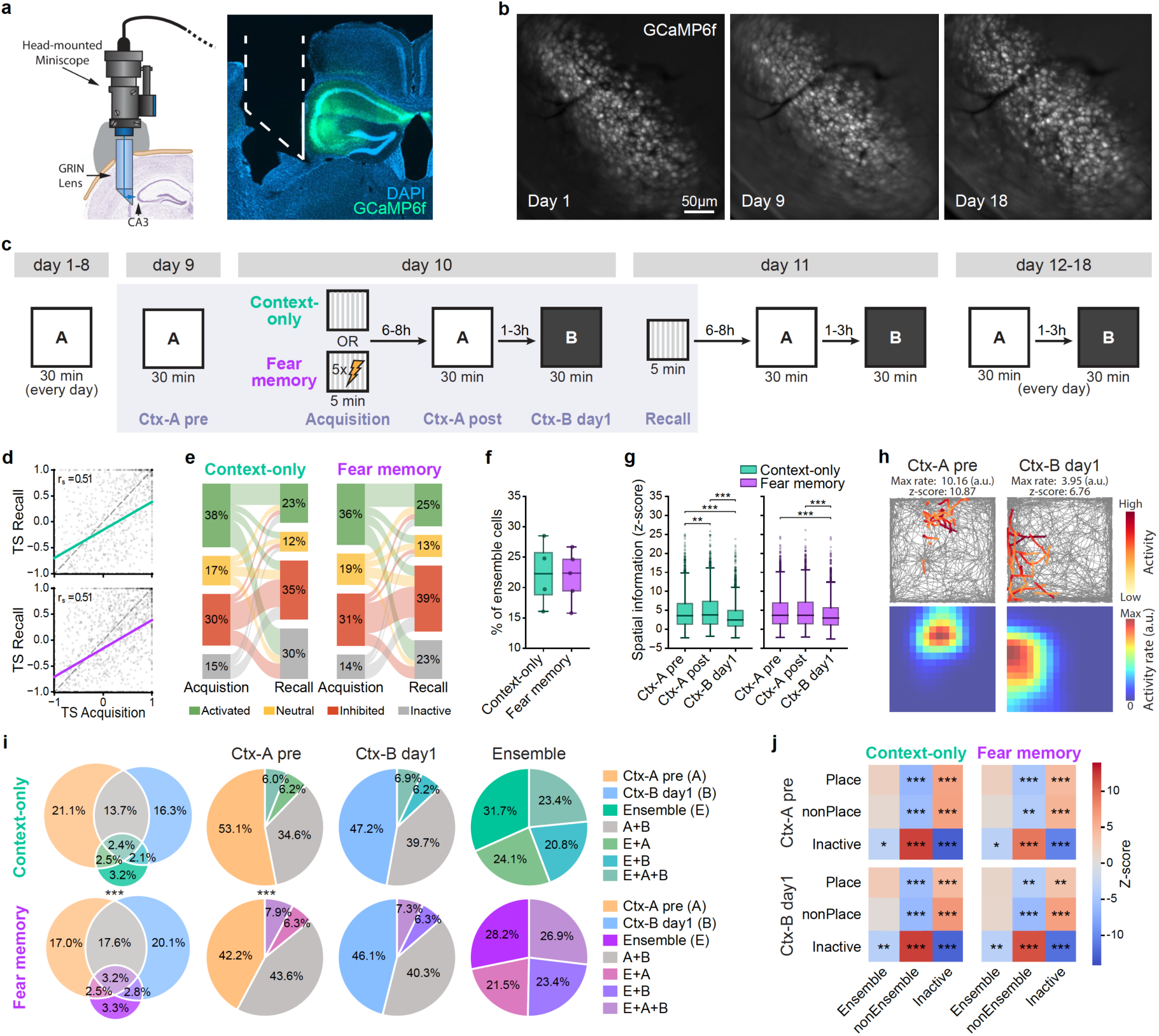
Memory ensembles and cognitive maps converge on overlapping neuronal populations, but coding dimensions are functionally orthogonalized in the network. (**a**) Implant location for miniscope recordings. Left, schematic representation of the implant location for miniscope recordings; right, histology revealing the position of the microendoscope (white dashed outline) and the prism imaging face (solid white outline) in relation to the hippocampal formation. (**b**) Example of a stable field of view. Maximum intensity projections of the same recording field of view across different days of the experiment. (**c**) Schematic of the experiment timeline. Mice were imaged while freely exploring two contexts (Ctx-A, Ctx-B) over the course of 18 days, and underwent contextual fear conditioning acquisition or a neutral exploration experience in a Training Context (TC) on day 10. The TC, Ctx-A and Ctx-B were located in different behavior rooms. The lightly shaded box highlights the five key behavioral sessions used for the longitudinal analyses: Ctx-A pre (day 9 in Ctx-A), Acquisition in the TC, Ctx-A post (day 10 in Ctx-A), Ctx-B day1 (day 1 in Ctx-B), and Recall in the TC. (**d**) Tuning scores (TS) were positively correlated between acquisition and recall in both groups (Spearman correlation on detected cells in both sessions. Context-only: r_s = 0.51, n = 1,345 cells from 4 animals, *p* < 0.001. Fear memory: r_s = 0.51, n = 2,100 cells from 5 animals, *p* < 0.001). The same relationship held within each animal (mean within-animal r_s = 0.51 and 0.49 for the two groups); with only 4 (context-only) and 5 (fear memory) animals the per-animal permutation test is at or near its minimum attainable *p* (sign-flip permutation test, 10,000 two-tailed resamples; context-only *p* = 0.12, fear memory *p* = 0.06), so inference here is based on the cell-level correlation. Each dot is one cell; the dashed line is the identity and the colored line the linear regression fit per group. Analysis included only cells detected in both the acquisition and recall sessions. (**e**) Transition of neurons between functional classes from acquisition to recall. Each node is a functional class: activated (green), neutral (yellow), inhibited (red), or inactive (grey). Each flow represents the number of cells transitioning from their acquisition class (left) to their recall class (right); flow width is proportional to cell count. Cells detected in at least one of the two sessions are included; cells detected in only one session are assigned to the inactive class for the session in which they were absent. Ensemble neurons are defined as cells classified as activated in both acquisition and recall (the activated ◊ activated flow), requiring above-threshold positive tuning during both encoding and retrieval. (**f**) The size of the memory ensemble did not differ between groups (independent permutation test, 10,000 two-tailed resamples. Diff is fear memory minus context-only. Diff = +0.49, *p* = 0.86). Each dot is one animal (context-only n = 4, fear memory n = 5); boxes show the group median and interquartile range, whiskers the full range. Values are the percentage of ensemble neurons per animal, among neurons detected in both acquisition and recall. (**g**) Spatial information changed across sessions within groups but did not differ between groups. Within groups, context-only animals showed a small increase from Ctx-A pre to Ctx-A post and a large decrease in Ctx-B day1 relative to both, whereas fear memory animals showed no change from Ctx-A pre to Ctx-A post but the same decrease in Ctx-B day1 (cell-level linear mixed models with animal as random intercept, REML; BH-FDR-corrected across all contrasts within each analysis. Coefficients in units of z. Context-only: Ctx-A pre → Ctx-A post, coef = +0.29, *p* = 0.004; Ctx-B day1 relative to both Ctx-A sessions, coef = −1.19, both *p* < 0.001. Fear memory: Ctx-A pre → Ctx-A post, *p* = 0.83; Ctx-B day1 relative to both Ctx-A sessions, coef = −0.89, both *p* < 0.001). Between groups, no session differed after correction (all *p* ≥ 0.38). Boxes show the cell-level distribution (median, interquartile range, whiskers to 1.5 × IQR, outliers as dots). Statistical inference was performed with animal as a random intercept, whereas the boxes show the pooled cell-level distribution. (**h**) Example place cells in sessions Ctx-A pre and Ctx-B day1. Top, animal trajectory (grey) in the open field during Ctx-A pre and Ctx-B day1, with example cells’ activity overlaid and color-coded by amplitude (low to high); bottom, corresponding spatial activity rate maps for the same cells. The values above each trajectory indicate the peak activity rate (a.u.) and the spatial information z-score of the corresponding place cell. (**i**) Ctx-A pre place cells, Ctx-B day1 place cells, and ensemble neurons converge on partially overlapping neuronal populations (cell-level chi-square tests of independence. Overall distribution: χ²(6) = 43.4, *p* < 0.001. Ctx-A pre place-cell distribution: χ²(3) = 33.8, *p* < 0.001. Ctx-B day1 distribution: χ²(3) = 0.36, *p* = 0.95. Ensemble distribution: χ²(3) = 2.73, *p* = 0.43. The same direction was present at the animal level: fear memory 52.6 ± 11.8%, context-only 40.6 ± 2.9% of Ctx-A pre place cells retained as Ctx-B day1 place cells, independent permutation test, diff = +11.9 percentage points, *p* = 0.10). Left, Venn diagram of the three populations; right, pie charts showing, within each population, the fraction that also belongs to the other two. Rows are the two groups (context-only, top; fear memory, bottom). The denominator is all cells detected in at least one of the sessions underlying these populations (Ctx-A pre and Ctx-B day1 for the place cell populations, acquisition and recall for ensemble membership). Asterisks between the Venn diagrams indicate a significant difference in the overall overlap distribution between groups, whereas asterisks between the pie charts indicate significant differences for the corresponding population. (**j**) Both groups showed the same pattern of enrichment and depletion of spatial map classes within ensemble categories (within-animal Monte Carlo permutation. For each combination of ensemble category (Ensemble, nonEnsemble, Inactive) and exploration session (Ctx-A pre, Ctx-B day1), we tested whether the fraction of cells also classified as Place, nonPlace, or Inactive differed from chance given the marginal proportions. Spatial map labels were shuffled within each animal, 10,000 permutations, preserving per-animal cell counts and spatial class proportions, and aggregated across animals to form a population-level null. Z-scores give the observed value’s deviation from this null in s.d. units, positive = enrichment, negative = depletion; two-tailed permutation *p*-values are the fraction of shuffles at least as extreme as observed, Holm-corrected across the 18 tests per group, 2 sessions × 3 spatial classes × 3 ensemble categories). Per comparison, the denominator is all cells detected in at least one relevant session (acquisition, recall, or the exploration session, excluding globally silent cells).

Animals were first familiarized with an Open-Field arena (“Ctx-A”) through nine days of repeated daily exploration (Figure 1c). On day 10, animals were split into two groups: one underwent contextual fear conditioning acquisition in a distinct training context (TC; “fear memory” group), while the other explored the same training context without shock (“context-only” group). Following TC exposure (“memory acquisition”), all animals re-explored the now familiar Ctx-A and were introduced to a novel open-field arena (“Ctx-B”). On day 11, re-exposure to the TC (“memory recall”) elicited robust freezing selectively in animals from the fear memory group (Figure S1a), confirming successful formation of a context-specific aversive memory. Animals were reintroduced to Ctx-A and Ctx-B on the same day and additionally performed daily exploration of both contexts for the next seven days, allowing us to track how cognitive maps evolved with repeated experience following memory acquisition. Exploratory behavior in Ctx-A and Ctx-B was indistinguishable between groups throughout the protocol, indicating that fear did not generalize beyond the training context and did not lead to the formation of retrospective or prospective linking (Figure S1b-g). This interleaved design allowed us first to ask whether mnemonic and spatial representations are allocated to distinct CA3 populations (a memory ensemble and a cognitive map, respectively) or instead emerge from a shared network component. The specific sequence of experiences further allowed us to determine whether associative memory encoding affects the cognitive map of an already familiar environment and the formation of a new one.

To establish a common analytical framework for comparing memory ensembles and cognitive maps within the same longitudinally tracked population, we focused on five key behavioral episodes: memory acquisition and recall in the TC, exploration of the familiar environment before and after memory acquisition (Ctx-A pre and Ctx-A post), and first exploration of a novel environment following memory acquisition (Ctx-B day1) (Figure 1c). Field-of-view alignment confirmed stable imaging quality and high-quality longitudinal tracking of individual neurons across these episodes (Figure S2). Because of the high-quality tracking and the fact that every tracked neuron was detected in at least one episode, the absence of detection in a given session was treated as session-specific functional inactivity in an otherwise stably registered cell. Among the longitudinally tracked population, the fraction of active neurons were similar across groups up to memory acquisition, after which fear memory animals consistently exhibited a larger activated population across all experiences (memory recall and open field explorations), consistent with broader recruitment of the CA3 network after an aversive learning experience (Figure S2a). This increase in the active population reflected both neurons retained from previous sessions and neurons newly activated after conditioning, without a selective expansion of either class (Figure S2b). The distribution of activity rates across the recorded population, however, did not differ between groups (Figure S2f), suggesting that conditioning increased the fraction of the CA3 network activated during subsequent experience without detectably altering the activity rates of participating neurons.

Tracking the same neurons across sessions let us define memory ensembles as those activated during both memory acquisition and retrieval relative to a home-cage baseline, overcoming single-snapshot classifications based on immediate early genes^3,6,8,11–13,75^. To identify memory ensemble neurons, we focused on acquisition and recall sessions in the TC and calculated a tuning score for each neuron reflecting its change in activity in the TC relative to baseline activity recorded in the home cage immediately before TC exposure^8,76^. Tuning scores were comparable across groups within conditions, and consistently lower during recall than acquisition (Figure 1d and S3a). Neurons were classified as activated, neutral, inhibited, or inactive based on their tuning score; neurons activated during both acquisition and recall were defined as memory ensemble neurons (Figure 1e and S3b). The proportion of memory ensemble neurons did not differ between the fear memory and context-only groups, indicating that the size of the recruited ensemble was comparable across the two memories despite differences in salience and valence of the two learning episodes (Figure 1f).

To identify neurons participating in spatial representations, we quantified spatial information of each active neuron during exploration of Ctx-A and Ctx-B. Spatial information was continuously distributed across the recorded CA3 population in both groups (Figure 1g). Within-group differences reflected known discrepancies between familiar and novel environments; across-group differences were non-significant. Neurons whose spatial information exceeded chance levels were defined as place cells and formed the core of each cognitive map (Figure 1h). The fraction of place cells remained stable between Ctx-A pre and Ctx-A post in both groups (Figure S3c). During the first exploration of the novel Ctx-B, it decreased in the context-only group but remained comparable to Ctx-A pre and Ctx-A post in the fear memory group (Figure S3c), suggesting that the formation of an associative memory may attenuate the novelty-associated reduction in place-cell recruitment. Place-cell fractions did not differ significantly between groups in any individual context (Figure S3d).

Having independently characterized memory ensembles and place cells within the same longitudinally tracked population, we next asked whether these two forms of hippocampal representations converge on overlapping neurons or recruit distinct CA3 subpopulations. Individual CA3 neurons could simultaneously contribute to the familiar context map, the novel context map, and the memory ensemble (Figure 1i). Similar recruitment patterns were observed when comparing the two sessions in the familiar Ctx-A before and after memory acquisition (Figure S3e). In the fear memory group, larger fractions of both the familiar cognitive map and the memory ensemble also contributed to the novel cognitive map. Since the corresponding fraction of the novel cognitive map contributing to either representation was comparable between the two groups, this asymmetry reflected the increased proportion of CA3 neurons active across sessions occurring after memory acquisition. These data thus establish that mnemonic and spatial representations converge on partially overlapping neuronal populations rather than being implemented by distinct and exclusive subnetworks.

This overlap, however, was non-random. Monte Carlo simulations revealed that memory-ensemble neurons were less likely than expected by chance to be silent during exploration of both familiar and novel environments (Figure 1j, S3f). This re-recruitment did not reflect a privileged subpopulation with an intrinsically greater capacity for spatial coding, since ensemble neurons were equally likely to be classified as place or non-place cells in both Ctx-A and Ctx-B, and place cells were no more likely to be ensemble neurons than expected by chance (Figure S3g). Moreover, ensemble and non-ensemble neurons exhibited indistinguishable levels of spatial information in both contexts (Figure S3h). Ensemble membership therefore predicted greater availability for recruitment across experiences, but not stronger or preferential spatial coding. This broader recruitment was also not associated with elevated baseline event rates, which instead were significantly lower for ensemble neurons, indicating that it was not simply explained by persistently higher baseline activity (Figure S3i, S3j). Complementarily, non-ensemble neurons were more likely to remain inactive during random foraging, whereas neurons that were inactive during both memory acquisition and recall were preferentially recruited during exploration of Ctx-A and Ctx-B, again without any bias toward place-cell identity (Figure 1j). Together, these reciprocal biases indicate that mnemonic recruitment and spatial tuning follow structured but partially independent allocation rules, revealing a functional orthogonalization of these coding dimensions despite their convergence on a shared neuronal substrate. Importantly, these allocation biases were comparable across the fear memory and context-only groups, indicating that associative memory formation did not alter the underlying relationship between mnemonic and spatial representations.

The reciprocal biases we observed between memory-session identity and recruitment into cognitive maps at the pairwise session level suggested that individual CA3 neurons might exhibit recurring recruitment propensities across episodes, rather than being recruited independently to each representation. To determine whether these pairwise biases combined into a coherent population-level structure, we examined joint functional profiles in two complementary three-way comparisons: memory-session identity together with place-cell identity across familiar and novel cognitive maps, and memory-session identity together with place-cell identity across two exposures to the familiar cognitive map before and after acquisition (Figure 2a). Under a null model in which functional identities were assigned independently while preserving their frequency within each animal and condition, a neuron’s role in any one representation would carry no information about its role in another, and the expected frequency of each three-way profile would be given by the product of the marginal frequencies of its constituent categories. Departures from this expectation would indicate that particular combinations of functional identities recur across experiences more often than can be explained by their prevalence within the individual sessions alone.

**Figure 2:**
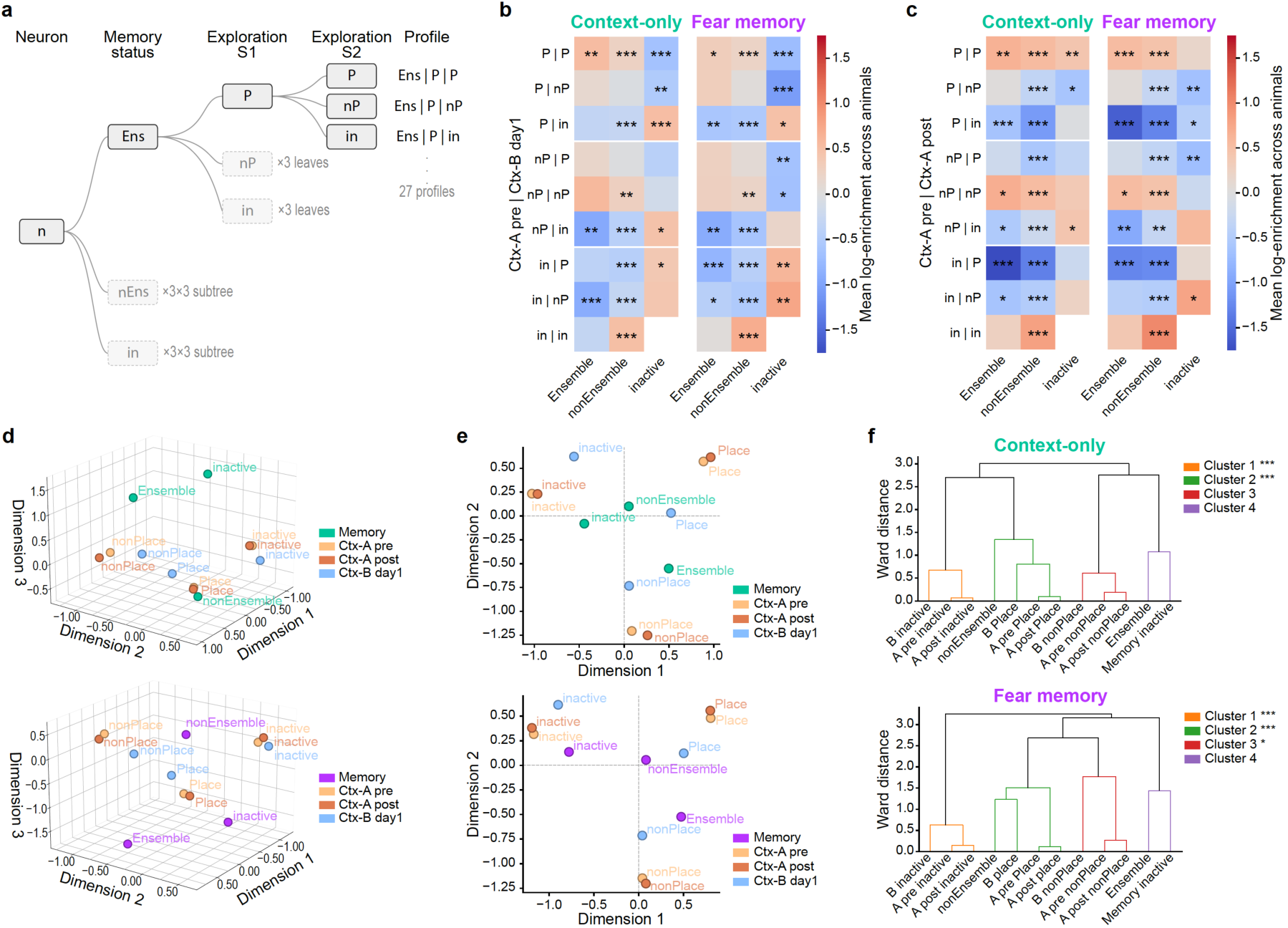
Joint profiling of memory and spatial identity reveals organized population structure. (**a**) Profiling tree for neuron category assignment. Each neuron is assigned a categorical profile combining its memory status (Ensemble/Ens, nonEnsemble/nEns, inactive/in) with its spatial map classification (Place/P, nonPlace/nP, inactive/in) in each of two exploration sessions (S1, S2). The tree enumerates these combinations (memory status × S1 class × S2 class), giving 3 × 3 × 3 = 27 profiles, of which the all-inactive profile (in | in | in) cannot occur since the denominator requires detection in at least one session; 26 profiles are therefore observable. One branch is drawn in full (Ens → P → …); collapsed branches (dashed) follow the same structure and are annotated with the number of leaves (×3) or sub-branches (×3 × 3) they represent. Each leaf corresponds to one profile in the co-occurrence heatmaps (**b**, **c**). The denominator includes all neurons detected in at least one of the four relevant sessions (acquisition, recall, and the two exploration sessions). (**b**) Both groups showed a comparable overall pattern of co-occurrence enrichment and depletion across the profiles defined in **a**, for the Ctx-A pre (S1) × Ctx-B day1 (S2) pairing (permutation test, 10,000 iterations: condition labels shuffled independently within each animal, preserving marginal distributions, observed and expected counts recomputed, and the observed mean log-enrichment tested against the resulting null; exact two-tailed p-values BH-FDR-corrected across the 26 profiles. Profiles of interest are considered in the main text). Color is the mean log-enrichment across animals, log((observed + 0.5) / (expected + 0.5)), computed separately for context-only and fear memory animals, with expected counts computed per animal under independence of memory status and spatial class and +0.5 Laplace smoothing for zero-count profiles; red indicates enrichment above chance, blue depletion, near-white chance. Rows are the two sessions’ spatial classes (labelled S1 | S2), columns the memory status; white borders group rows by first-session class. (**c**) As in **b**, both groups showed a comparable overall pattern of enrichment and depletion, for the Ctx-A pre (S1) × Ctx-A post (S2) pairing; color scale shared with **b**. (**d**) Multiple correspondence analysis (MCA) of neuron profiles across memory and spatial conditions, shown separately for context-only and fear memory animals, on the first three MCA dimensions. Unlike the heatmaps in **b** and **c**, which pair two exploration sessions, the MCA profiles each cell across all four conditions: memory status, and spatial map classification in Ctx-A pre, Ctx-A post, and Ctx-B day1. Each condition contributes three labels, giving 12 label-condition points, colored by condition. A point’s position reflects how strongly its label co-occurs with the others, so points lying close together tend to appear in the same neurons. The axes are MCA dimensions rather than directly interpretable variables, but relative positions reveal the structure of label co-occurrence across the population. Variance (inertia) explained by each dimension is shown in Supplementary figure 4d. (**e**) The same MCA projected onto the first two dimensions, showing the 12-label condition points separately for context-only and fear memory animals, colored by condition. This projection shows the dominant structure captured by the leading dimensions. (**f**) Hierarchical clustering of the 12 label-condition points in MCA space resolved an equivalent four-cluster structure in both groups, with the Spatial-inactive and Place clusters significantly compact in each (Ward hierarchical clustering, minimizing total within-cluster variance at each merge, in the dimensions exceeding the mean inertia of 12.5%: the first three for context-only and the first four for fear memory, so compactness was assessed within each group’s own MCA space; cluster compactness was quantified as the mean pairwise Euclidean distance between within-cluster points (smaller = tighter) and tested against a within-animal shuffled null representing marginal label frequencies alone, 10,000 permutations, one-tailed, BH-FDR-corrected across the four clusters. Both groups: the three Spatial-inactive labels formed one cluster and the three Place labels together with nonEnsemble another, both significantly compact, all *p* < 0.001; the three nonPlace labels formed a cluster that was compact in fear memory only, *p* = 0.020, Context-only *p* = 0.57; and Ensemble with Memory-inactive formed a cluster compact in neither group, both p ≥ 0.9999). In the dendrogram, k = 4 was selected based on the largest increase in Ward linkage distance (elbow criterion), which occurred when merging four into three clusters in both groups, consistent with visual inspection of the dendrogram. Branch colors indicate cluster membership, and corresponding clusters are shown in the same order and color across both groups to facilitate direct comparison. Branches above the color threshold are black. The null was generated by shuffling condition labels independently within each animal, preserving per-animal frequencies and overall category abundances but destroying cross-condition association, rerunning the MCA on each shuffled dataset, and recomputing within-cluster distances under the original cluster assignments. In the panel, A pre, A post, and B denote Ctx-A pre, Ctx-A post, and Ctx-B day1.

We therefore quantified the frequency of all 27 possible profiles within each comparison. Their distributions departed markedly from the independent-recruitment model, with strong enrichment of specific profiles (Figure S4a, b). Among the most prominently enriched were profiles reflecting place-cell identity in both open-field sessions, whether these comprised repeated exploration of the familiar Ctx-A or exploration of the familiar and novel contexts (Figure 2b, c). These profiles were enriched among both ensemble and non-ensemble neurons, indicating that a stable propensity to participate in cognitive maps is associated with broad recruitment across experiences, but not with preferential allocation to a memory ensemble of that experience. A distinct enriched profile comprised neurons that contributed to the familiar map in Ctx-A but were inactive in the novel Ctx-B and during memory acquisition and recall, revealing a population selectively available to the familiar spatial representation. Conversely, profiles characterized by inactivity in both open-field sessions but consistent activation during memory acquisition and recall were strongly enriched among non-ensemble neurons. Finally, active neurons classified as non-place cells across both open-field sessions and not-ensemble during TC sessions formed another prominently enriched profile. Rather than constituting an inactive reserve, these neurons define a broadly engaged yet consistently untuned population, demonstrating that persistent activation across experiences does not necessarily entail incorporation into any of the spatial or mnemonic representations measured here. Together, these joint profiles demonstrate that the pairwise allocation biases observed across sessions combine into recurring functional configurations within the CA3 network. Their similar organization across the fear memory and context-only groups further indicates that fear memory formation does not detectably reorganize these underlying allocation biases.

Because each analysis considered only a subset of the animals’ experiences, we next asked whether the same relationships were evident when the five focal behavioral episodes were considered jointly. We therefore considered each neuron’s functional identity across all four conditions simultaneously (its classification as a place cell, non-place cell, or inactive neuron in Ctx-A pre, Ctx-A post, and Ctx-B day 1, together with its classification as an ensemble, non-ensemble, or inactive neuron during memory acquisition and recall) and applied Multiple Correspondence Analysis (MCA) to the resulting categorical matrix. Whereas the preceding enrichment analyses identified specific combinations of identities that recurred across selected session sets (Figure S4c), MCA summarized how these categorical identities co-varied across the analyzed episodes. The resulting high-dimensional point cloud could be summarized by a smaller number of dominant axes identified by their elevated inertia (akin to variance in Principal Component Analysis, Figure S4d), revealing a simple geometrical organization for neuronal recruitment to spatial and mnemonic representations aligned along three main dimensions (Figure 2d). The first MCA dimension primarily separated inactive from active identities across conditions, defining an axis of general neuronal availability for recruitment into hippocampal representations (Figure 2d, e). The second dimension separated place-cell from non-place-cell identities among active neurons, organizing the correspondence space into distinct spatially tuned and untuned subspaces (Figure 2e). Within each subspace, the Ctx-A pre and Ctx-A post labels occupied neighboring positions, whereas the corresponding Ctx-B day1 labels were displaced, indicating that functional identities were organized both by spatial tuning and by the environment in which that identity was expressed. Ensemble and non-ensemble neurons were fully segregated along the third MCA dimension (Figure S4e, S4f).

Within the first two MCA dimensions, memory-session identities were embedded within this geometry rather than segregating from the spatial categories. Ensemble identity occupied the region of the correspondence space associated with non-place categories, whereas non-ensemble identity was positioned closer to the place-cell region (Figure 2e, S4e, S4f). Thus, across the complete four-condition configuration, neurons classified as ensemble and neurons classified as non-place participated in similar multivariate patterns of recruitment, while non-ensemble and place-cell identities showed a complementary alignment. This geometrical association does not imply that ensemble neurons were less likely to become place cells within any individual session: ensemble and non-ensemble neurons displayed equivalent session-specific probabilities of place-cell classification (Figure 1j). Rather, proximity in the MCA space indicates that these identities occurred within similar combinations of labels across the complete experiment, revealing differences in their longitudinal organization that were not apparent from marginal probabilities alone.

To test whether the organization revealed by MCA contained reproducible structure beyond visual proximity in the correspondence space, we applied Ward hierarchical clustering to the label-condition coordinates in the retained MCA dimensions. This analysis identified four major clusters in each cohort, corresponding to biologically interpretable groupings of functional labels (Figure 2f). In both context-only and fear memory animals, labels associated with inactivity across open-field sessions clustered together, whereas place-cell labels across Ctx-A pre, Ctx-A post, and Ctx-B formed a separate cluster, consistent with a shared spatial-tuning axis across environments. Non-place labels also grouped together, indicating that active but spatially untuned identities occupied a distinct region of the MCA space. Memory-session labels were embedded within this organization rather than forming an isolated domain: non-ensemble identity clustered with spatially active labels, whereas ensemble or memory-session inactivity labels occupied neighboring but distinct positions depending on group. To determine whether these clusters reflected structure beyond the marginal abundance of each label, we compared their compactness with a Monte Carlo null in which condition labels were shuffled within each animal while preserving category frequencies. Several individual clusters were significantly more compact than expected after FDR correction, including the inactive and place-cell clusters in the context-only group and the inactive, place-cell/non-ensemble, and non-place clusters in the fear memory group (Figure 2f). Thus, specific combinations of memory– and spatial-coding identities formed reproducible, compact regions of the MCA low-dimensional space. Importantly, we interpret these clusters as structured co-occurrence patterns among functional identities, rather than as evidence for immutable cellular classes, because individual neurons could still move between representational roles across experiences (Figure 1i).

Because the labels entering the MCA were derived from activity-based measurements, activity levels themselves could contribute to the observed organization. We therefore compared event rates between ensemble and non-ensemble neurons across sessions to identify if differences in their clustering in MCA space might be explained by them. Ensemble neurons showed lower event rates than non-ensemble neurons during baseline home-cage recordings (Figure S3i), and became selectively more active during acquisition and recall in the TC, consistent with their operational definition as neurons recruited by the TC experience (Figure S4g). In stark contrast, during exploration of Ctx-A and Ctx-B ensemble and non-ensemble neurons exhibited comparable event rates (Figure S4g, S4h). This equivalence was preserved when the analysis was restricted to non-place cells, excluding the possibility that differences in place-field activity masked a more general activity difference. It also persisted when only the initial minutes of open-field exploration were considered, indicating that this difference did not reflect a transient novelty or context-entry response. Ensemble neurons were therefore selectively more active during memory-related TC episodes while firing at rates comparable to non-ensemble neurons during open-field exploration, indicating that the organization captured by the MCA could not be trivially attributed to a general difference in activity level.

Together, these findings indicate that CA3 memory ensembles and cognitive maps for consecutive experiences are neither assigned to separate neuronal substrates, nor are they recruited indiscriminately from a common pool. Instead, the same CA3 population supports representations through structured, partially orthogonal allocation rules. Individual neurons remain functionally flexible, with many cells capable of participating in both across experiences. However, their recruitment is biased by stable population-level relationships among availability to recruitment, spatial tuning propensity, context specificity, and memory-session engagement. Memory-ensemble identity is expressed selectively during memory-related episodes, whereas spatial coding emerges along distinct axes of neuronal recruitability, tuning, and environmental specificity. This organization creates a shared but structured representational substrate: mnemonic and spatial codes overlap at the level of individual neurons, but separate at the level of population architecture. Importantly, this architecture was preserved across groups, indicating that the formation of an associative memory, despite expanding subsequent CA3 recruitment, did not overwrite the network’s pre-existing allocation logic.

### Forming an associative fear memory increases the precision of a new cognitive map

We next asked whether associative memory encoding influenced the quality of the spatial maps built by the same CA3 network. To address this question, we used a Bayesian position decoder as a population-level readout of coding quality. We compared decoder performance during exploration of the familiar environment before (Ctx-A pre) and after (Ctx-A post) memory acquisition, and during first exploration of a novel environment before (Ctx-A day1) and after memory acquisition (Ctx-B day1). Because fear memory animals recruited a larger active neuronal population after conditioning, we kept the number of neurons used to train the decoder constant between cohorts (N=100).

Decoding error in a familiar map remained comparable between Ctx-A pre and Ctx-A post both within the fear memory and context-only groups, indicating that fear memory encoding did not interfere with the precision of an established spatial representation (Figure 3a, S5a). The result was different for the first exposure to a novel context. In the context-only group, decoding error in Ctx-B was significantly higher than in the familiar context and similar to a novel exposure to Ctx-A, consistent with the lower precision expected for a newly formed spatial representation (Figure 3b, S5a). In contrast, the fear memory group exhibited a significantly more precise Ctx-B day1 map, with decoding precision statistically indistinguishable from the level observed in Ctx-A after nine days of prior familiarization (S5a). Thus, following associative fear memory formation, CA3 generated, upon first exposure to a novel environment, a spatial representation with familiar-like precision.

**Figure 3:**
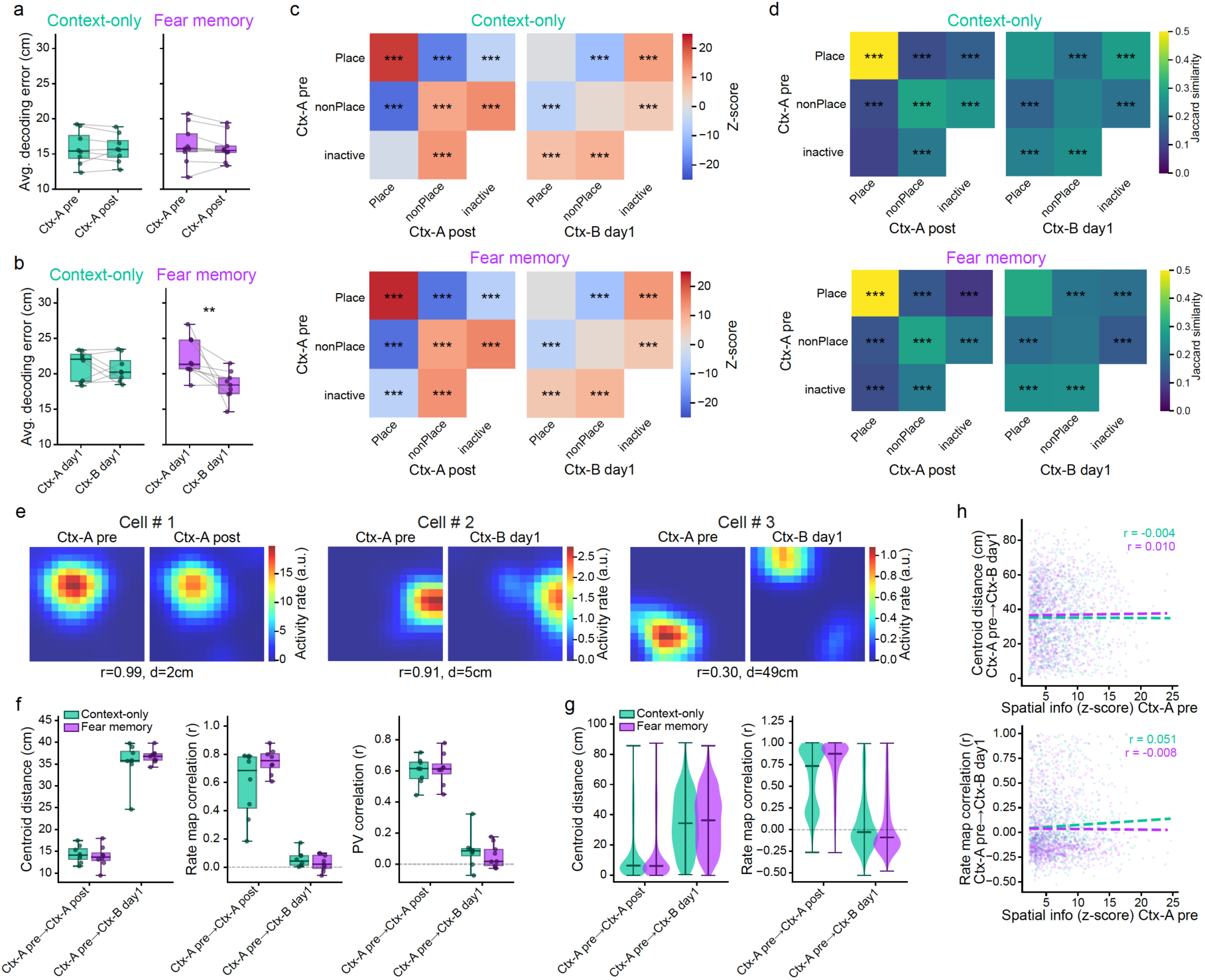
Increased precision of a cognitive map formed after fear conditioning, yet intact context discrimination. (**a**) Bayesian population vector decoding error did not differ between the Ctx-A pre and Ctx-A post sessions in either group (paired permutation test, 10,000 two-tailed resamples. Context-only: Δ = −0.087, *p* = 0.82. Fear memory: Δ = −0.418, *p* = 0.24). Each dot is the median decoding error of one animal, boxes show the group median and interquartile range, whiskers the full range, and lines connected paired observations. Decoding was performed on a random subsample of 100 neurons to control for differences in the number of detected neurons across groups and sessions. (**b**) Bayesian population vector decoding error was lower during Ctx-B day1 than during Ctx-A day1 in the fear memory group, whereas it did not differ between sessions in the context-only group (paired permutation test, 10,000 two-tailed resamples. Context-only: Δ = −0.485, *p* = 0.53. Fear memory: Δ = −3.830, *p* = 0.008). Plotted and subsampled as in **a**. (**c**) Neurons classified as place cells in Ctx-A pre were more likely than chance to be classified as place cells again in Ctx-A post, whereas place cell identity did not persist above chance between Ctx-A pre and Ctx-B day1. Heatmaps show Monte Carlo enrichment z-scores for all 3 × 3 combinations of spatial class in session 1 (rows: Place, nonPlace, Inactive) and session 2 (columns), for Ctx-A pre → Ctx-A post (left) and Ctx-A pre → Ctx-B day1 (right), shown separately for the context-only and fear memory groups. Z-scores quantify enrichment or depletion relative to chance, estimated by independently shuffling session 2 spatial class labels within each animal while preserving spatial class proportions (10,000 permutations). All cells detected in at least one session were included and the inactive × inactive combination was masked. Asterisks indicate exact two-tailed permutation *p* values, Holm-corrected across all non-masked comparisons. (**d**) The same pattern held for the overlap of the place cell populations themselves: overlap exceeded chance between Ctx-A pre and Ctx-A post but not between Ctx-A pre and Ctx-B day1. Heatmaps show the Jaccard similarity index for the same 3 × 3 combinations, session pairings and groups as in **c**. Significance was assessed by independently shuffling spatial class labels in both sessions within each animal while preserving spatial class proportions (10,000 permutations), with cells included and masking as in **c**. Asterisks as in **c**. (**e**) Rate maps for three representative place cells, each shown in the Ctx-A pre session (left) and a second session (right), illustrating the range of within– and across-context map similarity. Cell # 1 is compared between two exposures to the same context (Ctx-A pre → Ctx-A post), and cells # 2 and 3 across contexts (Ctx-A pre → Ctx-B day1). Color indicates activity rate (a.u.) in each spatial bin. For each cell, r is the Pearson correlation between the two activity maps computed over spatially co-visited bins, and d is the Euclidean distance (cm) between the place field centroids. (**f**) Place cells remapped more strongly across contexts (Ctx-A pre → Ctx-B day1) than across repeated sessions in the same context (Ctx-A pre → Ctx-A post) in both groups, shifting their place field centroids further and showing lower rate map and population vector (PV) correlations (paired permutation test, 10,000 two-tailed resamples, BH-FDR-corrected across the two groups for each metric. Centroid distance: context-only, Δ = +21.1 cm, *p* = 0.009; fear memory, Δ = +22.5 cm, *p* = 0.009. Rate map correlation: context-only, Δ = −0.53, *p* = 0.008; fear memory, Δ = −0.72, *p* = 0.008. PV correlation: context-only, Δ = −0.51, *p* = 0.010; fear memory, Δ = −0.55, *p* = 0.010). The extent of remapping did not differ between groups for either session pair (independent permutation test, 10,000 two-tailed resamples, BH-FDR-corrected across the two session pairs for each metric, all *p* > 0.19). PV correlation is the correlation of population activity-rate vectors across spatial bins. Analysis included only cells classified as place cells in both sessions. Δ is the across-context minus the within-context value. Boxes show the group median and interquartile range of the per-animal means, whiskers the full range, and each dot is one animal. (**g**) The same pattern was apparent across individual place cells. Violin plots show the cell-level distributions of centroid distance and rate map correlation for within-context stability (Ctx-A pre → Ctx-A post) and across-context remapping (Ctx-A pre → Ctx-B day1). Inclusion criteria and metrics are as in **f**. Statistical comparisons were performed on the per-animal summary values shown in **f**. (**h**) Spatial tuning strength in Ctx-A pre did not predict the extent of across-context remapping in either group. Neither the mean within-animal correlation between Ctx-A pre spatial information and remapping magnitude differed from zero (sign-flip permutation test, 10,000 two-tailed resamples, n = 17 animals. Centroid distance: mean r = 0.004, *p* = 0.84. Rate map correlation: mean r = 0.022, *p* = 0.42), nor did this correlation differ between groups (independent permutation test, 10,000 two-tailed resamples. Centroid distance: *p* = 0.76. Rate map correlation: *p* = 0.47). Top, Ctx-A pre spatial information z-score versus Ctx-A pre → Ctx-B day1 centroid distance. Bottom, the same versus rate map correlation. Each point is one place cell, and analysis included only cells classified as place cells in both sessions (n = 1,998 cells total; context-only, n = 778; fear memory, n = 1,220). Regression lines are shown for each group. Pearson r values displayed in the panels are based on pooled cell-level data, whereas statistical inference was performed on the per-animal correlations.

We next directly compared decoding precision between cohorts (Figure S5c-f). With decoder input fixed at 100 neurons per session, decoding error in Ctx-B day1 was significantly lower in the fear memory group than in the context-only group, whereas no statistical difference was observed for either the Ctx-A pre or Ctx-A post familiar maps. When decoders were instead trained on the entire recorded population, the difference in Ctx-B became even more robust (Figure S5e). Thus, including the additional neurons recruited after fear conditioning further increased the precision advantage of this experimental group, indicating that the expansion of network recruitment carried additional spatial information. Removing all conventionally defined place cells increased decoding error in both cohorts (Figure S5f), but performance remained substantially better than that obtained after shuffling neuronal activity, demonstrating that neurons outside the conventionally defined place-cell population also carried spatial information. Under this condition, the fear memory group retained a numerical precision advantage of similar direction, although the between-group difference no longer reached statistical significance. Together, these results establish that fear memory encoding selectively enhanced the population-level precision of a newly formed map and suggest that this precision was not carried exclusively by conventionally defined place cells, but was distributed across the broader CA3 network.

Since a substantial population of neurons participated in both spatial representations of Ctx A and B, (Figure 1i, S3e), and neurons retained as place cells across sessions were more spatially informative than newly recruited place cells (both when they were retained across repeated explorations of Ctx-A and when they were retained across Ctx-A and Ctx-B, Figure S6a), we next tested whether the increased precision of the Ctx-B map in the fear memory group could reflect a failure to generate a genuinely novel representation, with animals instead reusing the extensively familiarized map of Ctx-A. We examined this possibility at two distinct levels: the reuse of the same neurons as place cells across maps, and the persistence of their spatial tuning across environments.

Neurons were more likely to be persistently classified as place cells across repeated sessions in Ctx-A than expected by chance (Figure 3c, d, S6b), indicating preferential preservation of place-cell identity within the familiar environment across repeated exposures. By contrast, place-cell overlap between Ctx-A and Ctx-B did not exceed that expected from independent recruitment, and many neurons classified as place cells in Ctx-A were inactive in Ctx-B, consistent with orthogonalization of the familiar and novel spatial representations (Figure 3c, d; S6c). Direct comparisons between cohorts revealed that the percentage of retained place cells from Ctx-A to Ctx-B was higher in the fear memory animals than in the context-only ones (Figure S6d). This was driven by the larger fraction of active cells detected in Ctx-B in this group, as the increased retention of previous place cells was accompanied by increased recruitment of novel place cells, while the relative distribution of retained and newly recruited place cells within the Ctx-B cognitive map was comparable between cohorts (Figure S6d, S6e). In line with this, the spatial information distribution showed only a small rightward shift in the fear memory group, detectable primarily among cells with lower spatial-information values (Figure S6f, g, S7a-c). This pattern supports the view that the increased Ctx-B precision was not concentrated in a small subset of highly informative retained neurons, but reflected a broader population-level change.

Finally, we tested whether associative memory formation caused retained place cells to preserve their spatial tuning across contexts, effectively carrying over the familiar Ctx-A map into Ctx-B. In both groups, place cells retuned more strongly across contexts than across repeated sessions in the same context, as revealed by reduced spatial information z-score correlations (Figure S7d, e), reduced activity rate correlations (Figure S7f, g), and stronger shifts in place-field centroids and reduced single-cell rate-map and population-vector correlations (Figure 3e-g). Moreover, spatial tuning strength did not predict the extent of cross-context remapping (Figure 3h), suggesting that neurons with a stable capacity for spatial coding did not preserve the specific spatial information encoded in the familiar environment.

Together, these findings reveal that associative memory formation changes the population-level organization of the spatial code, allowing a novel cognitive map to emerge with familiar-like precision while leaving the precision of established familiar maps intact and preserving contextual specificity across maps.

### Associative memory formation prevents experience-dependent refinement of a new cognitive map despite preserved components of cellular functional plasticity

The familiar-like precision of Ctx-B during the first exposure to this context after contextual fear conditioning raises the question of whether the associative memory formation shifted the entire trajectory of familiarization toward higher coding precision, or whether it had bypassed the experience-dependent refinement that normally follows the formation of a new cognitive map. To distinguish between these possibilities, we examined how place cell and population-level coding properties evolved across longitudinal Ctx-B explorations and compared these trajectories with the initial familiarization of Ctx-A, when animals were still naïve.

During naïve familiarization with Ctx-A, its cognitive map underwent a refinement process. Decoding error decreased between the first exposure and later days of familiarization, indicating that spatial coding precision improved with repeated experience (Figure 4a, Figure S8a-c). This improvement was accompanied by coordinated changes across the CA3 population. The proportion of place cells increased (Figure 4b), whereas neurons with weak spatial tuning became less active (Figure S8d, e). Place cells carried more spatial information (Figure 4c and S8f), became more selective for a preferred location (Figure S8g), and fired less frequently outside their place fields (Figure 4d). They also showed fewer place fields (Figure S8h), and at the population level, the number of fields representing each spatial bin decreased (Figure S8i). Together, these changes indicate that familiarization progressively sharpens and optimizes the spatial code, with a progressive expansion of highly informative place cells.

**Figure 4:**
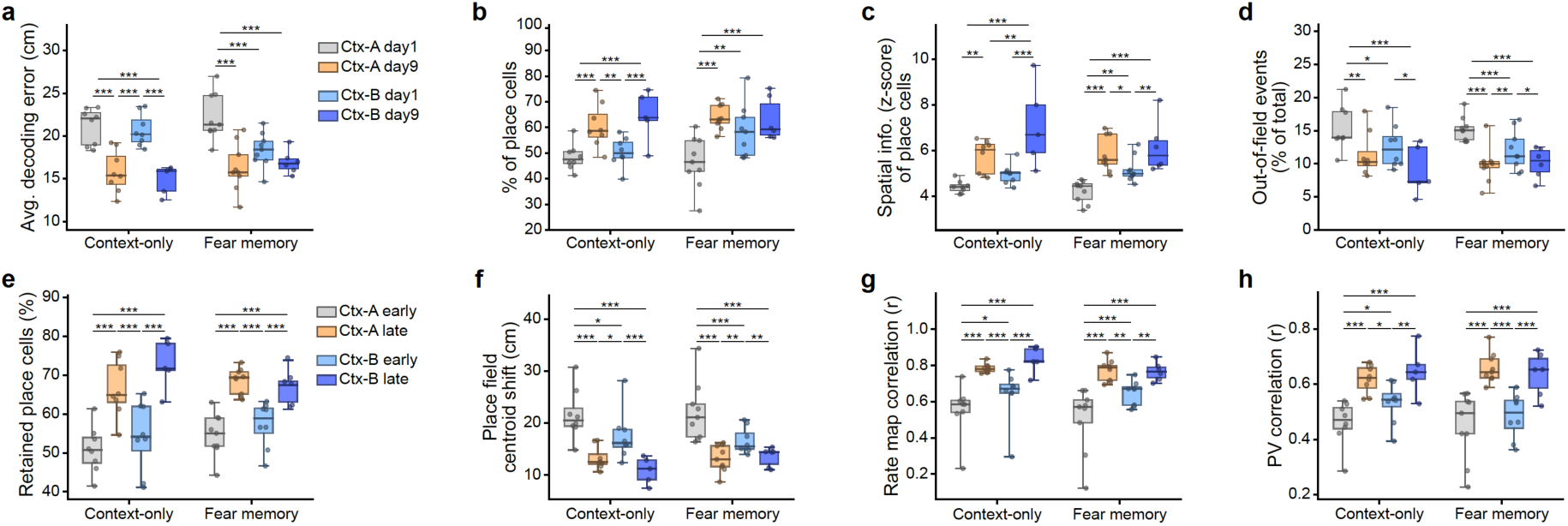
Associative fear memory formation selectively prevents experience-dependent refinement of population-level spatial coding. (**a**) Bayesian population vector decoding error decreased from the first to the last day of familiarization in both contexts in context-only animals, but in fear memory animals this improvement was absent in Ctx-B (linear mixed-effects model with session as a categorical fixed effect and animal as random intercept, REML; pairwise contrasts BH-FDR-corrected across all comparisons. Δ is in cm, positive values indicating lower error on the later session. Context-only: Ctx-A day1 → day9, Δ = +5.33, *p* < 0.001; Ctx-B day1 → day9, Δ = +4.84, *p* < 0.001; Ctx-A day9 → Ctx-B day1, Δ = −4.84, *p* < 0.001; Ctx-A day1 → Ctx-B day9, Δ = +5.16, *p* < 0.001. Fear memory: Ctx-A day1 → day9, Δ = +5.88, *p* < 0.001; Ctx-A day1 → Ctx-B day1, Δ = +3.83, *p* < 0.001; Ctx-A day1 → Ctx-B day9, Δ = +5.92, *p* < 0.001; Ctx-B day1 → day9, Δ = +0.84, *p* = 0.21, not significant). The absence of improvement across Ctx-B in fear memory animals contrasts with the corresponding improvement in Ctx-A of the same animals (Δ = +5.88) and with Ctx-B in context-only animals (Δ = +4.84). All remaining contrasts had *p* ≥ 0.0504. The model was chosen to handle the repeated-measures structure and missing data at later timepoints (attrition due to logistical constraints; missing-at-random assumed); sensitivity analyses (paired permutation, bootstrap, paired t-tests, Wilcoxon signed-rank) on complete-case pairs were consistent for the well-powered comparisons. Decoding used a random split of train and test frames during mobility. Each dot is one animal; boxes show the group median and interquartile range, whiskers the full range. (**b**) The percentage of place cells increased from the first to the last day of familiarization in both contexts in context-only animals, but in fear memory animals this increase was absent in Ctx-B, mirroring the decoding result in **a** (statistical approach as in **a**. Δ is in percentage points, negative values indicating more place cells on the later session. Context-only: Ctx-A day1 → day9, Δ = −12.20, *p* = 0.0002; Ctx-B day1 → day9, Δ = −14.47, *p* = 0.0002; Ctx-A day9 → Ctx-B day1, Δ = +10.21, *p* = 0.0017; Ctx-A day1 → Ctx-B day9, Δ = −16.43, *p* < 0.001. Fear memory: Ctx-A day1 → day9, Δ = −17.42, *p* = 0.0001; Ctx-A day1 → Ctx-B day1, Δ = −12.06, *p* = 0.0065; Ctx-A day1 → Ctx-B day9, Δ = −20.97, *p* = 0.0009; Ctx-B day1 → day9, Δ = −2.97, *p* = 0.38, not significant). The absence of an increase across Ctx-B in fear memory animals contrasts with the corresponding improvement in Ctx-A of the same animals (Δ = −17.42) and with Ctx-B in context-only animals (Δ = −14.47). All remaining contrasts had *p ≥* 0.29. Each dot is one animal; boxes show the group median and interquartile range, whiskers the full range. (**c**) The median spatial information z-score of place cells increased from the first to the last day of familiarization across Ctx-A and Ctx-B in both groups (statistical approach as in **a**. Δ is in z-score units, negative values indicating higher spatial information on the later session. Context-only: Ctx-A day1 → day9, Δ = −1.33, *p* = 0.003; Ctx-B day1 → day9, Δ = −2.20, *p* < 0.001; Ctx-A day9 → Ctx-B day9, Δ = −1.40, *p* = 0.008; Ctx-A day1 → Ctx-B day9, Δ = −2.61, *p* < 0.001. Fear memory: Ctx-A day1 → day9, Δ = −1.64, *p* < 0.001; Ctx-A day1 → Ctx-B day1, Δ = −0.97, *p* = 0.004; Ctx-A day9 → Ctx-B day1, Δ = +0.68, *p* = 0.038; Ctx-A day1 → Ctx-B day9, Δ = −2.12, *p* < 0.001; Ctx-B day1 → day9, Δ = −0.88, *p* = 0.007). The cross-context drop in spatial information from Ctx-A day9 to Ctx-B day1 reached significance in fear memory animals (Δ = +0.68) but not in context-only animals (Δ = +0.78, *p* = 0.081). Unlike decoding error (**a**) and place cell fraction (**b**), the increase in spatial information across Ctx-B was present in both groups. All remaining contrasts had *p ≥* 0.19. Each dot is one animal; boxes show the group median and interquartile range, whiskers the full range. (**d**) The median percentage of out-of-field events of place cells decreased from the first to the last day of familiarization across Ctx-A and Ctx-B in both groups, indicating more spatially precise firing with experience (statistical approach as in **a**. Δ is in percentage points, positive values indicating fewer out-of-field events on the alter sessions. Context-only: Ctx-A day1 → day9, Δ = +4.02, *p* = 0.003; Ctx-B day1 → day9, Δ = +3.00, *p* = 0.026; Ctx-A day1 → Ctx-B day1, Δ = +2.77, *p* = 0.034; Ctx-A day1 → Ctx-B day9, Δ = +7.22, *p* < 0.001. Fear memory: Ctx-A day1 → day9, Δ = +5.44, *p* < 0.001; Ctx-A day1 → Ctx-B day1, Δ = +3.18, *p* < 0.001; Ctx-A day9 → Ctx-B day1, Δ = −2.25, *p* = 0.005; Ctx-A day1 → Ctx-B day9, Δ = +5.07, *p* < 0.001; Ctx-B day1 → day9, Δ = +1.74, *p* = 0.031). The cross-context rise in out-of-field events from Ctx-A day9 to Ctx-B day1 reached significance in fear memory animals (Δ = −2.25) but not context-only animals (Δ = −1.25, *p* = 0.30). Like spatial information (**c**), the reduction in out-of-field events across Ctx-B was present in both groups. All remaining contrasts had *p ≥* 0.13. Each dot is one animal; boxes show the group median and interquartile range, whiskers the full range. (**e**) Place cell retention across days increased from early to late exposure within both contexts in both groups (statistical approach as in **a**, but comparing exposure periods rather than single sessions: early = days1–3, late = days 7–9, with retention rate computed for each consecutive day pair within a period and averaged per animal. Δ is in percentage points, negative values indicating higher retention in the later period. Context-only: Ctx-A early → late, Δ = −15.68, *p* < 0.001; Ctx-B early → late, Δ = −17.42, *p* < 0.001; Ctx-A late → Ctx-B early, Δ = +12.25, *p* < 0.001; Ctx-A early → Ctx-B late, Δ = −22.20, *p* < 0.001. Fear memory: Ctx-A early → late, Δ = −13.93, *p* < 0.001; Ctx-A late → Ctx-B early, Δ = +10.91, *p* < 0.001; Ctx-A early → Ctx-B late, Δ = −13.20, *p* < 0.001; Ctx-B early → late, Δ = −6.92, *p* = 0.0007). The cross-context drop in retention from late Ctx-A to early Ctx-B was significant in both groups (Context-only Δ = +12.25; Fear memory Δ = +10.91). The increase in retention across Ctx-B was present in both groups. All remaining contrasts had *p ≥* 0.070. Each dot is one animal; boxes show the group median and interquartile range, whiskers the full range. (**f**) The change in place field centroid location across days decreased from early to late exposure within both contexts in both groups, indicating that place fields stabilized with experience (statistical approach and period definitions as in **e**. Δ is in cm, positive values indicating smaller centroid shifts in the later period. For each stable place cell, the centroid shift is the Euclidean distance between the primary place field’s center of mass across sessions, averaged per animal; larger values indicate more remapping. Context-only: Ctx-A early → late, Δ = +8.40, *p* < 0.001; Ctx-B early → late, Δ = +6.40, *p* < 0.001; Ctx-A late → Ctx-B early, Δ = −4.31, *p* = 0.014; Ctx-A early → Ctx-B early, Δ = +4.09, *p* = 0.017; Ctx-A early → Ctx-B late, Δ = +11.62, *p* < 0.001. Fear memory: Ctx-A early → late, Δ = +8.93, *p* < 0.001; Ctx-A early → Ctx-B early, Δ = +6.18, *p* < 0.001; Ctx-A late → Ctx-B early, Δ = −2.90, *p* = 0.009; Ctx-A early → Ctx-B late, Δ = +9.91, *p* < 0.001; Ctx-B early → late, Δ = +3.79, *p* = 0.009). The cross-context increase in centroid shift from late Ctx-A to early Ctx-B was significant in both groups (Context-only Δ = −4.31; Fear memory Δ = −2.90). The stabilization of place fields across Ctx-B was present in both groups. All remaining contrasts had *p ≥* 0.20. Each dot is one animal; boxes show the group median and interquartile range, whiskers the full range. (**g**) Rate map correlation across days increased from early to late exposure within both contexts in both groups, indicating that firing maps stabilized with experience (statistical approach and period definitions as in **e**. Δ is in correlation units, negative values indicating higher correlation in the later period. For each stable place cell, rate map correlation is the Pearson r between two sessions’ activity-rate maps over commonly visited bins, averaged per animal; higher values indicate more stable maps. Context-only: Ctx-A early → late, Δ = −0.23, *p* < 0.001; Ctx-B early → late, Δ = −0.20, *p* < 0.001; Ctx-A late → Ctx-B early, Δ = +0.14, *p* < 0.001; Ctx-A early → Ctx-B early, Δ = −0.09, *p* = 0.039; Ctx-A early → Ctx-B late, Δ = −0.28, *p* < 0.001. Fear memory: Ctx-A early → late, Δ = −0.27, *p* < 0.001; Ctx-A early → Ctx-B early, Δ = −0.16, *p* < 0.001; Ctx-A late → Ctx-B early, Δ = +0.12, *p* = 0.0025; Ctx-A early → Ctx-B late, Δ = −0.30, *p* < 0.001; Ctx-B early → late, Δ = −0.13, *p* = 0.0035). The cross-context drop in rate map correlation from late Ctx-A to early Ctx-B was significant in both groups (Context-only Δ = +0.14; Fear memory Δ = +0.12). The stabilization of activity rate maps across Ctx-B was present in both groups. All remaining contrasts has *p ≥* 0.32. Each dot is one animal; boxes show the group median and interquartile range, whiskers the full range. (**h**) Population vector (PV) correlation across days increased from early to late exposure within both contexts in both groups, indicating that population-level spatial coding stabilized with experience (statistical approach and period definitions as in **e**. Δ is in correlation units, negative values indicating higher correlation in the later period. For each animal, PV correlation is the Pearson correlation of the population activity-rate vector between sessions, computed per spatial bin and averaged across visited bins; higher values indicate more stable population coding. Context-only: Ctx-A early → late, Δ = −0.16, *p* < 0.001; Ctx-B early → late, Δ = −0.12, *p* = 0.007; Ctx-A late → Ctx-B early, Δ = +0.09, *p* = 0.013; Ctx-A early → Ctx-B early, Δ = −0.07, *p* = 0.030; Ctx-A early → Ctx-B late, Δ = −0.18, *p* < 0.001. Fear memory: Ctx-A early → late, Δ = −0.21, *p* < 0.001; Ctx-A late → Ctx-B early, Δ = +0.16, *p* < 0.001; Ctx-A early → Ctx-B late, Δ = −0.20, *p* < 0.001; Ctx-B early → late, Δ = −0.12, *p* < 0.001). The cross-context drop in PV correlation from late Ctx-A to early Ctx-B was significant in both groups (Context-only Δ = +0.09; Fear memory Δ = +0.16). The stabilization of population coding across Ctx-B was present in both groups. All remaining contrasts had *p ≥* 0.32. Each dot is one animal; boxes show the group median and interquartile range, whiskers the full range.

The same trajectory was observed during Ctx-B familiarization in the context-only cohort, with some measures showing an even steeper change. However, the fear memory cohort exhibited a markedly different trajectory. Decoder precision did not improve across repeated exploration (Figure 4a). The proportion of place cells did not increase, and selectivity and place-field number showed no experience-dependent change (Figure 4b, S8g, S8h). Thus, several of the coordinated transformations that normally accompany increasing map precision with familiarization were absent after associative memory formation. This did not, however, reflect a generalized loss of cell-based functional plasticity. Place cells showed a significant albeit modest increase in spatial information (Figure 4c), their out-of-field activity decreased across familiarization (Figure 4d), and neurons with weak spatial tuning became less active (Figure S4e). The CA3 network therefore remained responsive to repeated experience at the cellular level, but these ongoing cellular adjustments no longer translated into a progressive improvement in population-level coding precision. This altered population-level trajectory was specific to associative memory formation rather than shock exposure or physiological arousal. Animals subjected to an immediate-shock procedure, in which the amount and intensity of aversive stimulation through footshocks was comparable to the contextual fear conditioning protocol, but hippocampus-dependent associative learning was prevented (Figure S8j, k), exhibited the normal decrease of decoding precision for the first exposure to Ctx-B and experience-dependent improvement (Figure S8l).

We next asked whether the absence of experience-dependent refinement in the fear memory group reflected a failure to stabilize the newly formed cognitive map. During naïve familiarization with Ctx-A, the prevalence of stable place cells increased from early days to late days of exposure to the context (Figure 4e, S9a). Stable place cells were more active and spatially tuned than newly recruited place cells, and their tuning increased from early to late days in the context (Figure S9b, c). Stability initially emerged within a core of highly informative place cells before expanding to neurons spanning a broader range of spatial-information levels (Figure S9d). This population also became more tuned to the same location across days, as indicated by reduced remapping across late days in the context (Figure S9e-g). Familiarization thus normally combines representational turnover with the progressive stabilization of an increasingly broad and precise place cell population. These stabilization mechanisms were largely preserved during Ctx-B familiarization in both groups (Figure 4e-h; S9h-o). Several features were indistinguishable between the fear memory and context-only groups: the probability that a place cell retained its functional identity on the following day (Figure 4e), the higher activity and tuning of stable place cells than newly recruited ones (Figure S9l-m), preferential early stabilization of highly informative neurons (Figure S9n), and the remapping of retained place cells (Figure 4f-h). Associative memory formation therefore did not alter the persistence of individual place cell representations across exposures.

The two groups differed in how quickly the Ctx-B cognitive map reached its full complement of place cells. In the context-only group, new place cells continued to be recruited during early familiarization, so the place cell fraction rose gradually across days. Because of this, only a limited fraction of the place cells on a given day had already been classified as place cells the day before (“prior place cells”) during early days (Figure S9i). In the fear memory group, the place cell fraction resembled that of a familiar map from the first day for Ctx-B, and this fraction did not increase further across days (Figure 4b). As a result, a larger proportion of the place cells contributing to the map on later days had already been recruited during early exploration, and the percentage of prior place cells was high from the earliest sessions (Figure S9i, k). Critically, this occurred even though the probability that any individual place cell survived across days was unchanged (Figure 4e), indicating that the higher retention reflects the map’s initial composition rather than enhanced stability of individual cells. The same pattern was evident in the higher early Jaccard overlap between Ctx-B maps (Figure S9j, k).

Together, these findings indicate that associative memory formation selectively prevents the normal experience-dependent refinement of population-level spatial coding for a novel cognitive map, despite preserved changes in several single-cell properties such as retention, remapping, and preferential stabilization of highly informative neurons. Cellular adaptation and representational drift were therefore no longer coupled to the progressive gain in population precision that normally accompanies familiarization.

### Associative memory formation decouples cognitive-map precision from latent embedding refinement

Having established that the effect of fear memory formation on the novel cognitive map could not be accounted for by modulation of properties of individual place cells, we reasoned that this effect likely arises from the collective organization of spatial information across the population. We therefore examined the low-dimensional structure of CA3 activity during Ctx-A and Ctx-B exploration. For each recording session, we constructed spatial population vectors by averaging the activity of a fixed subset of neurons across visits to each spatial bin, and applied principal component analysis (PCA) to these vectors (Figure 5a). CA3 spatial activity was organized in a low-dimensional subspace: the leading components captured most of the variance (Figure 5b, and S10a, b). This structure was comparable between the context-only and fear memory groups. Thus, fear memory did not expand, compress, or otherwise alter the dimensionality of CA3 population activity. Similar results were obtained when PCA was applied to temporally organized neural activity rather than spatial population vectors, indicating that the low-dimensional organization was not an artifact of spatial binning (Figure S10c-e).

**Figure 5:**
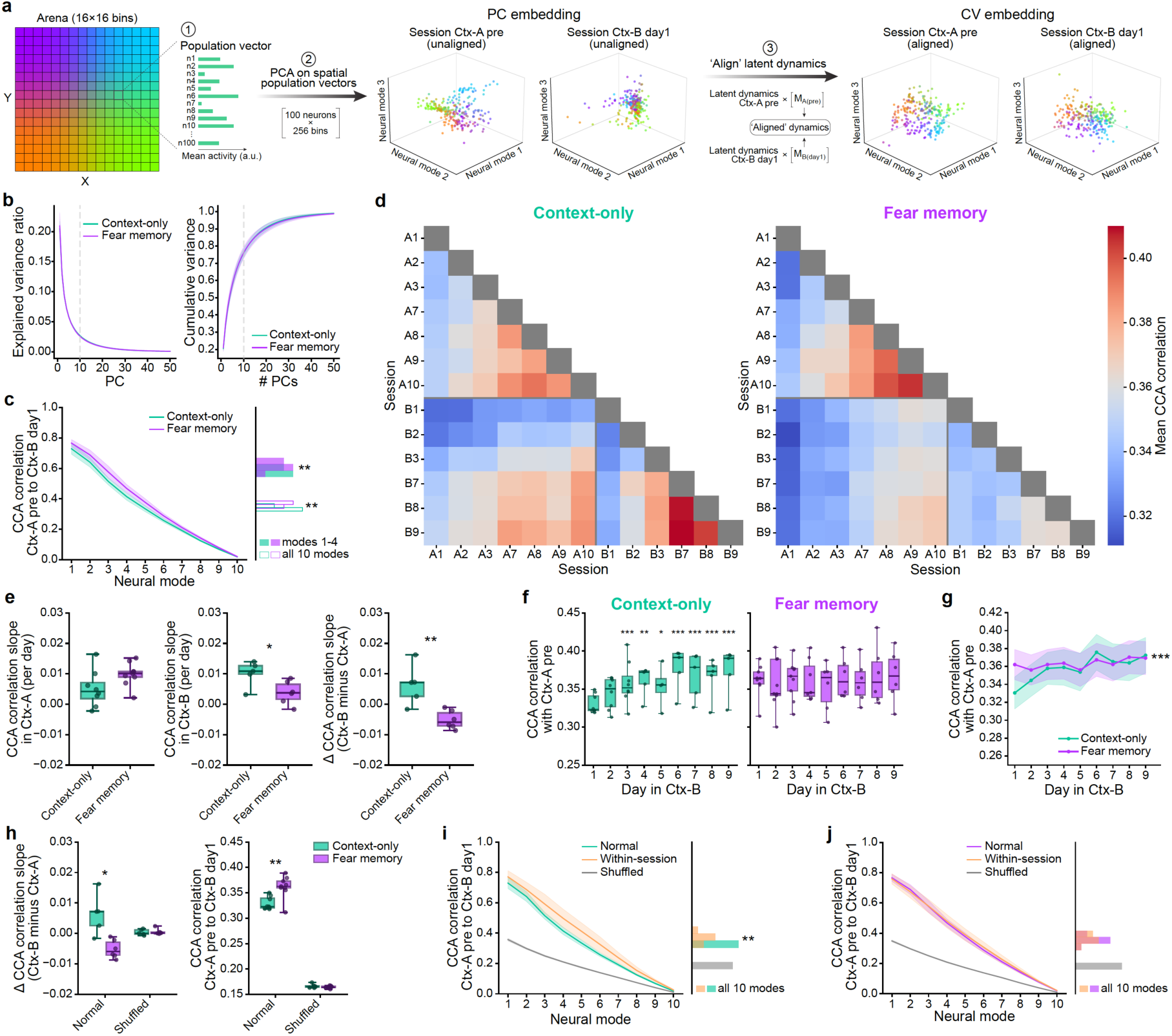
Fear memory formation causes a novel context to be represented in the latent regime of the familiar context from first exposure. (**a**) Schematic of the population-geometry analysis. (Step 1) For each session, a random subset of 100 neurons was selected, and spatial population vectors were constructed by averaging each neuron’s calcium signal across visits to the same spatial bin, summarizing activity in a common spatial reference frame while reducing bias from variable sampling density. (Step 2) Principal components analysis (PCA) was applied to these spatial population vectors, yielding session-specific neural modes corresponding to the leading principal components (PCs). The leading 10 PCs captured most of the variance and provided a low-dimensional representation of each session. (Step 3) Canonical correlation analysis (CCA) was applied to each session pair, aligning the corresponding 10-dimensional PC subspaces by maximizing their mutual correlation through linear transformations, yielding paired canonical variates (CVs) that define a shared latent space for cross-session comparison. The three-dimensional plots show the low-dimensional embedding of CA3 population activity for two example sessions from the same animal before (PC embedding) and after (CV embedding) CCA alignment. (**b**) The dimensionality of spatial population activity did not differ between context-only and fear memory animals. Neither the explained variance of individual principal components nor the cumulative variance differed between groups at any component (independent permutation tests, 10,000 two-tailed resamples, BH-FDR-corrected across the 50 components, all *p* ≥ 0.87 for both metrics), and the number of PCs required to reach 85% cumulative variance was also comparable (independent permutation test, 10,000 two-tailed resamples. Diff is fear memory minus context-only. Diff = −0.1 PCs, *p* = 1.000; context-only: n = 8, median 15.5, mean 15.4 ± 1.6; fear memory: n = 9, median 15, mean 15.3 ± 2.1). Left, mean explained variance ratio per principal component. Right, mean cumulative variance ratio as a function of the number of PCs included. Shading shows ± s.d. across animals. The dashed vertical line marks 10 dimensions, the number of PCs retained for the CCA analyses. PCA was performed on a random subsample of 100 neurons per session to control for differences in the number of detected neurons across groups and sessions, and repeated over 50 independent subsamples. Variance spectra were summarized as the median across subsampling iterations within each session, then the median across sessions within each animal, and finally the mean ± s.d. across animals within each group. (**c**) Canonical correlations between Ctx-A pre and Ctx-B day1 latent dynamics were higher in fear memory than context-only animals, both for the leading 4 modes and averaged across all 10 modes (independent permutation tests, 10,000 two-tailed resamples, BH-FDR-corrected across the two comparisons. Diff is fear memory minus context-only. Leading 4 modes: context-only 0.575 ± 0.024, fear memory 0.625 ± 0.034, diff = +0.050, *p* = 0.007. All 10 modes: context-only 0.330 ± 0.012, fear memory 0.362 ± 0.020, diff = +0.032, *p* = 0.007). Left, canonical correlation as a function of mode; lines show the group mean and shading the 95% CI across animals (t-based). Right, distribution across animals of the mean canonical correlation for the leading 4 modes (filled) and all 10 modes (unfilled); values are mean ± s.d. across animals. CCA was performed on the 10 leading principal components of each session, computed over spatially co-visited bins. Per iteration, 100 neurons were randomly subsampled without replacement within each session, independently for the two sessions being compared; each animal contributes the median canonical correlation across iterations. The leading 4 modes were analyzed separately following (Safaie et al., 2023). (**d**) Similarity between latent dynamics across session pairs, shown for context-only and fear memory animals. Color indicates the mean canonical correlation between the two sessions of each pair, with warmer colors indicating greater similarity; both heatmaps share a common color scale. Only the early and late sessions of ach context are shown (Ctx-A: A1–A3 and A7–A10; Ctx-B: B1–B3 and B7–B9), and the thick lines separate the 2 contexts. Only the lower triangle is shown; the diagonal is omitted, as within-session comparisons were not included. Subsampling and CCA were performed as in **c**. For each session pair, all ten canonical components were averaged into a single similarity value per animal, and heatmaps show the mean of these values across animals within each group. (**e**) Context-only animals increased CCA alignment across days more steeply during familiarization in Ctx-B than in Ctx-A, whereas fear memory animals showed the opposite pattern (independent permutation tests, 10,000 two-tailed resamples. Diff is fear memory minus context-only. Slope in Ctx A: context-only 0.0051 ± 0.0060, fear memory 0.0098 ± 0.0039, diff = +0.0047, d = +0.95, *p* = 0.071. Slope in Ctx-B: context-only 0.0101 ± 0.0042, fear memory 0.0038 ± 0.0038, diff = −0.0063, d = −1.58, *p* = 0.035. Δ: context-only 0.0063 ± 0.0067, fear memory −0.0051 ± 0.0030, diff = −0.0115, d = −2.30, *p* = 0.009). The Δ result was confirmed with a linear mixed model fitted to the raw session-pair correlations, which allowed animals with partial session coverage to be included (correlation ∼ day × group × context, with a random intercept and random slope for day per animal; n = 17 animals, 231 observations), in which the three-way interaction was significant (*p* < 0.001). Left, per-animal slope of cross-session canonical correlation on session day within Ctx-A (context-only n = 8, fear memory n = 9). Middle, the same within Ctx-B (context-only n = 5, fear memory n = 6). Right their difference (Δ = slope in Ctx-B minus slope in Ctx-A; context-only n = 5, fear memory n = 6). Slopes are in units of canonical correlation per day, and values are mean ± s.d. across animals. Boxes show the group median and interquartile range, whiskers the full range, and each dot is one animal. Slopes were estimated per animal by weighted linear regression of session-pair-averaged canonical correlation (mean across the 10 canonical modes) on session day, over sequential session pairs, with each day bin weighted by the number of contributing session pairs. (**f**) Alignment to Ctx-A pre increased across days in Ctx-B in context-only animals, whereas in fear memory animals it did not change (linear mixed-effects model fitted within each group, with log(day) as fixed effect and animal as random intercept, ML; omnibus likelihood-ratio test against an intercept-only model. Context-only: χ²(1) = 27.5, *p* < 0.001, n = 8 animals, 53 observations. Fear memory: χ²(1) = 2.05, *p* = 0.15, n = 9 animals, 64 observations). In context-only animals, post-hoc contrasts of each day against day 1 (categorical-day model, REML; BH-FDR-corrected across the eight contrasts. Δ is in units of canonical correlation, positive values indicating higher alignment than on day 1) were significant from day 3 onwards (day 3, Δ = +0.027, *p* < 0.001; day 4, Δ = +0.029, *p* = 0.002; day 5, Δ = +0.023, *p* = 0.017; day 6, Δ = +0.045, *p* < 0.001; day 7, Δ = +0.035, *p* < 0.001; day 8, Δ = +0.034, *p* < 0.001; day 9, Δ = +0.042, *p* < 0.001) but not on day 2 (Δ = +0.014, *p* = 0.065). Per-day contrasts were not tested in fear memory animals, as the omnibus test was non-significant. Left, context-only animals; right, fear memory animals. Each dot is one animal, boxes show the group median and interquartile range, whiskers the full range. Animal counts vary across days because of incomplete session coverage (context-only, n = 4–8; fear memory, n = 6–9). For each animal and session pair, values are the median canonical correlation across the leading 10 modes over 50 random-subsampling iterations of 100 neurons each, computed as in **c**. (**g**) Alignment to Ctx-A pre increased across days in Ctx-B in context-only animals, but not in fear memory animals, so that the two groups differed on day 1 and converged thereafter (linear mixed-effects model, correlation ∼ log(day) × with animal as random intercept, ML; likelihood-ratio test of the day × group interaction. χ²(1) = 13.3, *p* < 0.001. The interaction term gives the difference in slopes between groups: alignment increased 0.015 per log-day more steeply in context-only than in fear memory animal, 95% CI [0.007, 0.023]). Groups differed on day 1 (a priori contrast, reported uncorrected. Diff is fear memory minus context-only. Diff = +0.032, *p* = 0.011) but not on any subsequent day (BH-FDR-corrected across days 2– 9, all *p* ≥ 0.86). Lines show the estimated marginal means and shading the 95% CI, from a linear mixed-effects model with day as a categorical fixed effect crossed with group and animal as random intercept, REML; confidence intervals are from a t-approximation with degrees of freedom equal to the number of observations minus the number of fixed-effect parameters. Stars denote the omnibus day × group interaction. Correlation values were aggregated as in **f**. (**h**) The group differences in cross-context CCA alignment were abolished when spatial bin identity was shuffled, indicating that they reflect the spatial geometry of the population code (independent permutation tests, 10,000 two-tailed resamples, BH-FDR-corrected across the two versions within each metric. Diff is fear memory minus context-only. Δ slope: normal, context-only 0.0063 ± 0.0067, fear memory −0.0051 ± 0.0030, diff = −0.0114, *p* = 0.017; shuffled, context-only 0.0004 ± 0.0010, fear memory 0.0005 ± 0.0009, diff = +0.0001, *p* = 0.82. Ctx-A pre → Ctx-B day1 correlation: normal, context-only 0.3295 ± 0.0127, fear memory 0.3617 ± 0.0216, diff = +0.0322, *p* = 0.008; shuffled, context-only 0.1661 ± 0.0033, fear memory 0.1644 ± 0.0023, diff = −0.0017, *p* = 0.27). Within animals, shuffling reduced the Ctx-A pre → Ctx-B day1 correlation in both groups (paired permutation tests, 10,000 two-tailed resamples, BH-FDR-corrected across the two groups for each metric. Δ is normal minus shuffled. Context-only, Δ = +0.1634, *p* = 0.008; fear memory, Δ = +0.1973, *p* = 0.008), whereas the Δ slope did not differ significantly between versions in either group (context-only, Δ = +0.0059, *p* = 0.13; fear memory, Δ = −0.0057, *p* = 0.062); these within-group comparisons are not shown on the plot. For the Δ slope, with five and six animals per group, the smallest attainable two-tailed *p* values are 0.063 and 0.031, so these paired comparisons are not powered to detect a difference between versions after correction. Left, Δ CCA alignment slope (Ctx-B minus Ctx-A; context-only n = 5, fear memory n = 6) as in **e**. Right, canonical correlation between Ctx-A pre and Ctx-B day1 (context-only n = 8, fear memory n = 9) as in **b**. Each dot is one animal, boxes show the group median and interquartile range, whiskers the full range. Values are mean ± s.d. across animals. In the shuffled version, spatial bin labels were shuffled after computing the spatial population vectors, which breaks the geometry of the spatial map while leaving the activity dynamics unchanged. (**i**) In context-only animals, alignment between Ctx-A pre and Ctx-B day1 exceeded the shuffled baseline but remained below within-session alignment of Ctx-A pre, indicating that latent dynamics were aligned across contexts above chance without reaching within-session stability (paired permutation tests, 10,000 two-tailed resamples, BH-FDR-corrected across the three comparisons, n = 8. Δ is the first minus the second condition named. Ctx-A pre → Ctx-B day1 versus within-session: Δ = −0.049, *p* = 0.007. Ctx-A pre → Ctx-B day1 versus shuffled: Δ = +0.163, *p* = 0.007. Within-session versus shuffled: Δ = +0.213, *p* = 0.007). Left, mean canonical correlation per neural mode across animals for three conditions: cross-session CCA between Ctx-A pre and Ctx-B day1 (group color), within-session CCA of Ctx-A pre between its first and second half (orange, upper bound), and cross-session CCA from spatial-bin-shuffled data (grey, lower bound); shading shows the 95% CI across animals. Right, distribution across animals of the mean canonical correlation over the 10 modes, per condition. Values are the median across 50 random-subsampling iterations of 100 neurons each, aggregated as in **b**, and the shuffle is as in **h**. (**j**) In fear memory animals, alignment between Ctx-A pre and Ctx-B day1 was indistinguishable from within-session alignment of Ctx-A pre (first versus second half) and both exceeded the shuffle baseline, indicating that latent dynamics were as well aligned across contexts as within a single session (paired permutation tests, 10,000 two-tailed resamples, BH-FDR-corrected across the three comparisons, n = 9. Δ is the first minus the second condition named. Ctx-A pre → Ctx-B day1 versus within-session: Δ = −0.006, *p* = 0.63. Ctx-A pre → Ctx-B day1 versus shuffled: Δ = +0.197, *p* = 0.004. Within-session versus shuffled: Δ = +0.203, *p* = 0.004). The gap between cross-context and within-session alignment was smaller in fear memory than in context-only animals (independent permutation test, 10,000 two-tailed resamples. Diff is fear memory minus context-only. Context-only −0.0492 ± 0.0462, fear memory −0.0055 ± 0.0333, diff = +0.0437, *p* = 0.039). Panels as in **i**, for fear memory animals.

We next asked how spatial information was organized within this shared low-dimensional structure by training a linear decoder to predict animal position from neural activity projected onto the first *n* principal components. Position could be decoded from the leading principal components in both groups (Figure S10f, g), with decoding error falling steeply as more dimensions were considered. In Ctx-B day1, decoding from low-dimensional activity was consistent with the enhanced spatial precision observed in the Fear memory group when using the Bayesian position decoder (Figure S10h, I and Figure 3b), suggesting that spatial information was embedded within the dominant population modes rather than in a separate high-dimensional component.

Since the leading 10 principal components captured most of the variance in our data (>70%), and the resulting low-dimensional structure was comparable between groups, we used these dimensions as a common basis for comparing the latent organization of population activity across sessions within animals. We applied canonical correlation analysis (CCA)^68,72^ to each session pair, aligning two low-dimensional subspaces by maximizing their mutual correlation through linear transformations (Figure 5a, S11), and then computed the mean of the top X canonical correlations to measure the strength of the alignment. This approach allowed us to ask whether, within individual animals, repeated experience increased the consistency of the latent embedding of space across days. Across the full session-by-session alignment matrix, naïve animals showed an experience-dependent increase in latent similarity during familiarization to an environment: as Ctx-A became familiar, consecutive Ctx-A representations became progressively more aligned to each other (Figure 5d, e). Although latent alignment within familiar Ctx-A evolved comparably between groups, and Ctx-B showed the same across-day increase in the context-only group, this increase failed to emerge in Ctx-B after associative memory formation. As a result, the relative Ctx-B-versus-Ctx-A alignment slope (the day-to-day change in alignment between representations of a context) differed significantly between fear memory and context-only groups (Figure 5d, e). Thus, associative memory formation selectively disrupted the normal experience-dependent refinement of the novel-context embedding. Importantly, this result also argued against a simple ceiling explanation for the decoder data: despite high within-session precision on day 1, the Ctx-B latent embedding in the fear memory cohort had not already reached a mature, cross-day-stabilized state.

We then asked what kind of latent organization characterized the novel map in the absence of this normal refinement. One possibility was that Ctx-B did not begin as an unconstrained novel representation, but was instead biased toward the latent regime already established in the familiar environment. Consistent with this idea, alignment between the familiar Ctx-A pre representation and the first Ctx-B session was significantly higher in the fear memory group than in the context-only group (Figure 5c, d). Moreover, when Ctx-B was followed across repeated exposure, context-only animals progressively increased their alignment to Ctx-A pre, whereas fear memory animals started at a higher alignment level and showed no further increase across days (Figure 5f, g). Thus, the fear memory group did not gradually develop familiar-like organization through experience; instead, the novel map was biased toward the familiar latent regime from its first expression. This initial familiar-like organization approached the upper bound defined by within-session Ctx-A pre-alignment (Figure 5i, j). In context-only animals, Ctx-A pre–Ctx-B day1 alignment was significantly above the spatial-bin-shuffled baseline but remained below the alignment between two halves of the Ctx-A pre session, indicating a meaningful but incomplete correspondence between familiar and novel latent structures. In fear memory animals, by contrast, Ctx-A pre–Ctx-B day1 alignment was indistinguishable from within-session Ctx-A pre alignment while remaining well above the shuffled baseline. Therefore, after fear memory formation, the first representation of a novel environment occupied a latent regime normally associated with the familiar map.

These effects were not explained by the overall dimensionality of the data, activity scale, behavioral sampling, or arbitrary choices in the alignment procedure. The group differences were preserved when neuronal activity was z-scored or when binarized events were used (Figure S12a-e), when behavioral sampling was controlled for unvisited bins (Figure S12f-h), and when the number of PCA dimensions entered into CCA was varied across a broad range (Figure S13a-d). CCA alignment was also unrelated to the absolute dimensionality of each session or to dimensionality mismatch between session pairs (Figure S13e-g). By contrast, randomly shuffling spatial-bin labels, which preserves the population vectors and their covariance structure while disrupting their relationship to physical space, drastically reduced the alignment between the latent embeddings and abolished the group differences in both Ctx-A pre–Ctx-B day1 alignment and experience-dependent Ctx-B refinement (Figure 5h, Figure S14). This disruption strongly supported the result that CCA alignment did not simply reflect similarity in covariance structure or dimensionality, but depended on a consistent mapping between neural activity and physical locations. Finally, CCA alignment had direct functional relevance: the strength of latent alignment predicted the improvement in cross-session spatial decoding of animal position obtained after aligning the two latent spaces (Figure S15).

Together, these results show that associative memory formation changes how the context-specific representation of a novel environment is embedded within the low-dimensional manifold of CA3 population activity, while exerting no effect on the dimensionality of this activity. After associative memory formation, a novel environment is represented from its first exposure in a familiar-like latent regime, approaching the alignment normally observed within a single familiar session. Yet this familiar-like organization is not equivalent to a mature representation: cross-day alignment remains low and fails to progressively increase with experience. Associative memory formation therefore uncouples immediate spatial precision and familiar-regime alignment from the experience-dependent refinement of a context-specific latent spatial embedding.

## Discussion

Our results reveal that encoding an associative memory in a specific context changes how the CA3 network represents an experience that follows in a different one, by decoupling two processes that normally unfold together as a cognitive map is learned. Under naive conditions, repeated experience both sharpens the spatial code, so that position becomes more precisely decodable, and organizes the latent embedding of this population activity across exposures, so that the geometry of such embedding becomes increasingly consistent. Following associative memory formation, a novel environment is already encoded with familiar-like precision from its first exposure, but neither of these experience-dependent processes proceeds. Decoding precision does not improve, and the latent embedding does not converge across days. Associative memory formation therefore shifts CA3 into a distinct representational regime, in which a cognitive map can be read out with high accuracy from the outset, while its context-specific population embedding does not converge to a consistent geometry across repeated exposures to the context.

This effect emerged within a network in which memory ensembles and cognitive maps were neither assigned to separate neuronal substrates nor randomly superimposed onto the same neurons. Instead, mnemonic and spatial representations were multiplexed within a shared population according to structured allocation biases. Individual neurons retained the capacity to participate in multiple representational systems across experiences, but their recruitment was constrained by stable relationships among neuronal availability, spatial tuning propensity, context, and memory-session engagement. Mnemonic and spatial codes therefore overlapped at the level of individual cells, while remaining partially orthogonal at the level of the rules governing their allocation across the population. This architecture was preserved across groups, indicating that fear conditioning did not overwrite the pre-existing relationship between memory ensembles and cognitive maps. The altered trajectory of the novel map therefore cannot be attributed to a redistribution of neurons between mnemonic and spatial representation; instead, it points to a change in the state of the network in which both representations are formed.

Fear memory formation changed how a novel cognitive map was initialized, with the decoding precision of the novel Ctx-B map approaching levels expressed by the familiar Ctx-A map. This effect on the novel map was specific to an experience where animals had performed an associative memory between a stimulus and a context. In the experiments reported here, that stimulus was aversive, but neither exposure to the context alone (context-only group) nor delivery of the aversive stimulus under conditions that prevent associative learning (immediate shock group) reproduced the effect. Furthermore, enhanced precision was selective for the new environment, since the established Ctx-A map remained comparable before and after memory acquisition. Several features of the data indicated that this increased precision reflected a genuine change in the spatial code. Fixing decoder input to the same number of neurons preserved the precision advantage, and using the full recorded population strengthened it, indicating that the expanded recruitment observed after fear memory was not merely a sampling confound but carried additional spatial information. At the same time, the effect was not carried exclusively by conventionally defined place cells, nor did the novel map inherit the familiar Ctx-A place-field structure: place cell overlap across contexts remained limited and retained cells remapped to a similar extent across groups. Associative memory formation therefore increased the spatially informative capacity available to a new map while preserving its contextual specificity.

The novel context map not only was initialized with high precision, but it also failed to undergo the refinement that accompanies repeated experience. During familiarization with Ctx-A, and during Ctx-B familiarization in the context-only group, repeated exploration improved decoder precision and increased latent alignment across days. Thus, while in naïve conditions spatial accuracy within a session and consistency of the population embedding across sessions refined together, after associative memory formation both measures were flat across familiarization, but from different starting points. Decoding precision began elevated and remained consistently so, while latent alignment began at low levels and did not rise with experience. Associative memory formation therefore decoupled a cognitive map’s precision and the latent structure of its population dynamics from experience.

At the level of individual place cells, plasticity was preserved after associative memory formation. Place cells retained their identity across days with comparable probability; persistent place cells continued to remap; highly informative neurons still showed preferential stabilization; and place cells continued to increase spatial information and to reduce their out-of-field activity with repeated experience. What changed was the relationship between these cellular processes and the population code they support. During normal familiarization, changes in recruitment, tuning, and stabilization aggregate into a progressively more precise population representation and an increasingly consistent latent embedding. After fear memory formation, the cellular changes continued but no longer had these consequences. This dissociation also argues against the possibility that associative memory simply accelerated the maturation of the novel map, bringing it to a familiar-like state earlier. The effect of associative memory formation is therefore tied to hippocampal population dynamics, more than to single-neuron tuning.

A possible explanation is that the formation of an associative memory biases subsequent representations toward a pre-existing, familiar-like CA3 scaffold. Such a scaffold could provide an efficient set of population dimensions through which spatial position can be decoded accurately from the outset, explaining both the high initial precision of Ctx-B and its preferential alignment with the familiar Ctx-A representation. However, this familiar-like organization was not equivalent to a mature Ctx-B map. If Ctx-B had simply been advanced to a mature state, cross-session Ctx-B alignment should have started high or increased normally with repeated exposure. Instead, early Ctx-B alignment was low and comparable between cohorts, and increased only in context-only animals. In fear memory animals, alignment remained near its initial level despite repeated exploration. The novel map was therefore precise without being mature. This resembles, at a different level of description, the history dependence observed when environments are morphed between familiar shapes: when the sequence of environments does not support incremental modification, population activity can fall back into pre-established states rather than forming new ones^54, 59,64^. What our data suggest is a constrain that sits at a different computational level: rather than determining which stored representation is expressed as in the morphing experiments, prior experience appears to constrain the dynamical regime within which a new representation is constructed. In this model, associative memory formation preserves the CA3 network’s flexible access to a familiar latent scaffold, but the mapping of Ctx-B locations onto that scaffold remains labile across exposures. New spatial experiences can therefore be embedded in a familiar-like population regime without being forced to stabilize into a dedicated, context-specific latent organization.

One candidate circuit substrate for this regime is learning-induced plasticity in local inhibitory networks. Contextual fear conditioning shifts hippocampal circuits toward a high-PV, high-GAD67 network configuration^42–49^. Such a state could sharpen ongoing population activity by constraining weak or inconsistent patterns, increasing the immediate decodability of spatial position, while simultaneously limiting the slower experience-dependent remodeling through which a novel map would otherwise acquire a stable latent embedding. A single inhibitory reconfiguration could therefore account for both signs of the dissociation, unifying enhanced initial precision and absent refinement. This proposal remains untested here, but it offers a tractable circuit-level bridge between associative memory formation, initial map precision, and subsequent refinement.

Taken together, our findings indicate that cognitive maps are shaped by the state in which prior learning left the circuit that constructs them. That influence is legible not only in the precision of the map itself, but in how the map evolves with experience.

## Acknowledgements

We thank the animal facility, the research instrumentation facility, and the mechanical and electrical workshops of the Biozentrum of the University of Basel for excellent technical assistance. We are grateful to Jual Alvaro Gallego for hosting NI in his lab at ICL, and for inspiration, input, assistance, and feedback on the manifold analysis. We thank the whole Donato lab for the productive discussions and feedback on the project. We thank Kadjita Asumbisa and Maria Luce Rizzolo for feedback on the manuscript.

## Author contributions

F.D. and N.I. conceived the study. N.I. acquired data, with help from L.M. N.I. analysed the data, with input and help from F.D., C.G.C, and C.M. N.I. visualized the results. F.D. and N.I. wrote the manuscript, with feedback from all authors.

## Data and materials availability

All data will be made available in the manuscript, the supplementary material, or upon request.

**Figure S1:**
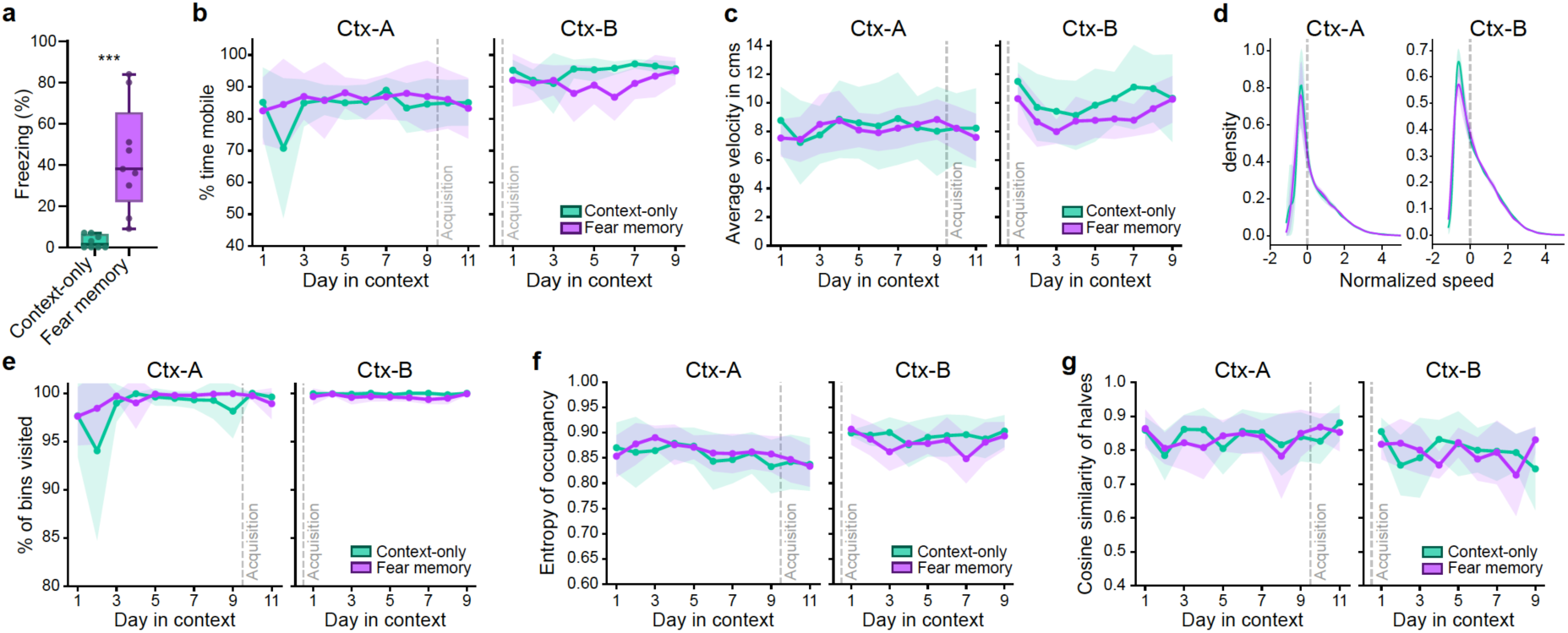
Freezing but not exploratory or locomotor behavior differed between groups. (**a**) Freezing during recall was higher in fear memory than context-only animals (independent permutation test, 10,000 two-tailed resamples. Diff is fear memory minus context-only. Fear memory 43.22 ± 25.91, context-only 2.75 ± 3.20, diff = +40.47, *p* < 0.001). Each dot is one animal (fear memory n = 9, context-only n = 8); boxes show the group median and interquartile range, whiskers the full range. (**b**) Overall activity level, measured as the percentage of mobile frames in a session, did not differ between context-only (n = 8) and fear memory animals (n = 9) in either context (linear mixed-effects models with log(day) as fixed effect and animal as random intercept, ML; trajectory divergence tested by likelihood-ratio test of the log(day) × group interaction. Ctx-A: interaction *p* = 0.50, day 1 *p* = 0.57. Ctx-B: interaction *p* = 0.92, day 1 *p* = 0.27). Per-day group contrasts (saturated categorical-day model, BH-FDR-corrected across days within each context) were examined only where the interaction was significant, and the a priori day 1 baseline is reported uncorrected; neither applied here, as no interaction reached significance. Lines show the group mean and shading the ± s.d. across animals. Ctx-A spans days 1–11 and Ctx-B days 1–9; in both panels the dashed lien marks the moment of memory acquisition (between Ctx-A days 9 and 10, i.e. before entry into Ctx-B). Missing data were assumed missing at random, and a sensitivity analysis restricted to animals present at all timepoints confirmed the result. (**c**) Movement speed during mobile frames did not differ between context-only and fear memory animals in either context (statistics as in **b**. Ctx-A: interaction *p* = 0.44, day 1 *p* = 0.24. Ctx-B: interaction *p* = 0.85, day 1 *p* = 0.18). Lines show the group mean and shading the ± s.d. across animals; the dashed line marks memory acquisition as in **b**. Values are the median speed of movement during mobile frames. (**d**) The distribution of movement speeds during mobile periods did not differ between context-only and fear memory animals in either Ctx-A post or Ctx-B day1 (animal-level permutation test on the proportion of frames in each quartile of the pooled speed distribution, 10,000 two-tailed resamples, BH-FDR-corrected across the four quartiles, all *p* ≥ 0.39). Instantaneous speeds were z-scored per animal (subtracting each animal’s median, dividing by its s.d.), removing individual speed differences while preserving distribution shape. Per-group distributions were estimated by kernel density estimation with bootstrap confidence bands (1,000 iterations, resampling animals with replacement); the dashed line marks the median (normalized speed = 0). Ctx-A post and Ctx-B day1 are the first exploration session of each context following acquisition. (**e**) Arena occupancy, measured as the percentage of spatial bins visited, did not differ between context-only and fear memory animals in either context (statistics as in **b**. Ctx-A: interaction *p* = 0.31, day 1 *p* = 0.97. Ctx-B: interaction *p* = 0.73, day 1 *p* = 0.14). Lines show the group mean and shading the ± s.d. across animals; the dashed line marks memory acquisition as in **b**. (**f**) Uniformity of space use, measured as the entropy of occupancy, did not differ between context-only and fear memory animals in either context (statistics as in **b**. Ctx-A: interaction *p* = 0.51, day 1 *p* = 0.43. Ctx-B: interaction *p* = 0.31, day 1 *p* = 0.64). Lines show the group mean and shading the ± s.d. across animals; the dashed line marks memory acquisition as in **b**. (**g**) Consistency of exploration, measured as the cosine similarity of occupancy across session halves, did not differ between context-only and fear memory animals in either context (statistics as in **b**. Ctx-A: interaction *p* = 0.95, day 1 *p* = 0.89. Ctx-B: interaction *p* = 0.95, day 1 *p* = 0.32). Lines show the group mean and shading the ± s.d. across animals; the dashed line marks memory acquisition as in **b**. Session halves are defined by equal numbers of mobile frames rather than by time.

**Figure S2:**
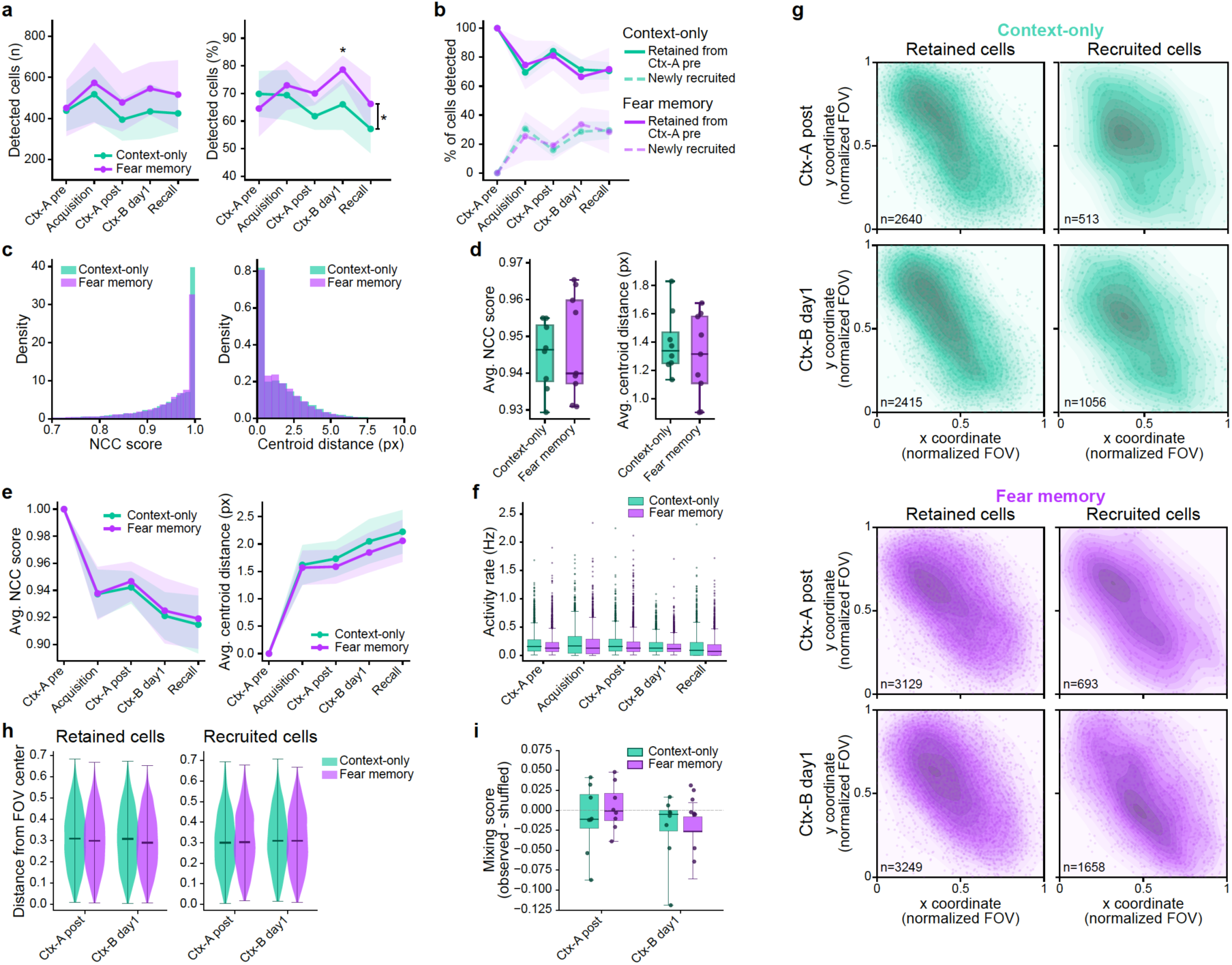
Higher post-acquisition cell detection in Fear memory animals is not accounted for by tracking quality, activity rate, or field-of-view sampling. (**a**) Cell detection was comparable between groups before acquisition but tended to be higher in fear memory animals from acquisition onward, reaching significance for the detection percentage at Ctx-B day1 (linear mixed modes, metric ∼ session × group with animal as random intercept; omnibus session × group interaction by likelihood-ratio test, ML; per-session contrasts from the fitted model, REML, BH-FDR-corrected across sessions. Diff is fear memory minus context-only. Detected cells (n): χ²(4) = 9.47, *p* = 0.0503; no individual session differed, all *p* ≥ 0.46, though the difference grew across sessions from +13 at Ctx-A pre to +111 at Ctx-B day1. Detected cells (%): χ²(4) = 11.80, *p* = 0.019; higher in fear memory at Ctx-B day1, diff = +12.58, *p* = 0.014, with no other session differing individually, all *p* ≥ 0.16). Left, number of detected neurons per session. Right, the same as a percentage of all tracked neurons. Lines show the group mean and shading the 95% CI across animals. This difference in detection is controlled for in the population analyses by subsampling to 100 neurons per session. (**b**) The balance of retained versus newly recruited cells did not differ between groups at any session (linear mixed models as in **a**; omnibus session × group interaction by likelihood-ratio test. Diff is fear memory minus context-only. χ²(4) = 4.03, *p* = 0.40; no session differed, all *p* ≥ 0.85). Lines show the proportion of detected cells retained from Ctx-A pre (solid) versus newly recruited (dashed), with the group mean and 95% CI across animals. As the two proportions sum to 100%, their contrasts are identical in magnitude with opposite sign. (**c**) Distributions of longitudinal tracking quality, shown per group and pooled across sessions, cohorts and animals. Left, normalized cross-correlation (NCC) score; right, centroid distance (pixels). Both are computed by the Inscopix longitudinal registration algorithm for all tracked cell pairs, where each value reflects how well a cell matches the cell assigned to the same global ID in another session, so higher NCC and lower centroid distance indicate better tracking. Distributions are normalized to density (n = 32,428 cell pairs from 17 animals; 13,896 from context-only and 18,532 from fear memory). (**d**) Per-animal tracking quality did not differ between groups (independent permutation tests, 10,000 two-tailed resamples. Diff is fear memory minus context-only. NCC score: diff = +0.002, *p* = 0.69. Centroid distance: diff = −0.10, *p* = 0.45). Per-animal means of NCC score (left) and centroid distance (right, pixels), collapsed across sessions and cohorts. Each dot is one animal (context-only n = 8, fear memory n = 9); boxes show the group median and interquartile range, whiskers the full range. (**e**) Tracking quality did not diverge between groups across sessions (linear mixed models, metric ∼ session × group with animal as random intercept, ML; likelihood-ratio test of the session × group interaction. NCC score: χ²(4) = 0.52, *p* = 0.97; centroid distance: χ²(4) = 2.26, *p* = 0.69). Lines show the group mean and shading the ± s.d. across animals (n = 9 animals: 4 Context-only, 5 Fear memory). Only data from animals recorded in all five longitudinally tracked sessions (cohort 1) are shown. The same pattern was observed for cohort 2 (see Methods; n = 8 animals: 4 context-only, 4 fear memory), which underwent the same behavioral protocol but was not imaged during Acquisition and Recall (NCC score: χ²(2) = 4.32, *p* = 0.12; centroid distance: χ²(2) = 4.97, *p* = 0.083). (**f**) Activity rates of detected neurons did not differ between groups at any session, by any of three analyses (cell-level linear mixed models, activity rate ∼ group with animal as random intercept, REML, BH-FDR-corrected across the five sessions, all *p* ≥ 0.09; and, because activity-rate residuals were strongly right-skewed (Shapiro–Wilk *p* < 0.001 at every session), independent permutation tests on per-animal rates, 10,000 two-tailed resamples, BH-FDR-corrected across sessions, using per-animal means (all *p* ≥ 0.19) and per-animal medians (all *p* ≥ 0.38); per-animal differences were at most 0.04 Hz). Nominally lower uncorrected *p*-values at acquisition and recall under the LMM (*p* = 0.037 and 0.030) and the mean-based permutation test (*p* = 0.048 and 0.079) were absent when animals were summarized by median rate (*p* = 0.19 and 0.37), consistent with these trends reflecting a minority of high-rate cells rather than a shift in the typical neuron. Boxes show the cell-level distribution of activity rates per group (median, interquartile range, whiskers to 1.5 × IQR, outlier cells as dots). Acquisition and recall include cohort 1 only (context-only n = 4, fear memory n = 5), as cohort 2 was not imaged during these sessions; the remaining sessions include both cohorts. (**g**) Location of retained and recruited cells in the recording field of view (FOV). Kernel density estimates of normalized cell locations in Ctx-A post (top) and Ctx-B day1 (bottom), shown separately for retained and recruited cells in each group. Retained cells were detected in both Ctx-A pre and the session shown; recruited cells were detected in that session but not Ctx-A pre. Per animal, x and y coordinates were scaled to [0, 1] using the extremes of all accepted cells in that session, so the detected population spans the normalized space regardless of absolute FOV size or position. Cell counts per group and session are shown in each panel’s lower-left corner. (**h**) The distance of retained and recruited cells from the FOV center did not differ between groups at either session (cell-level linear mixed models, distance from center ∼ group with animal as random intercept, REML, BH-FDR-corrected across the four comparisons (2 cell statuses × 2 sessions), all *p* ≥ 0.29; repeated at the animal level on per-animal mean distances by independent permutation test, 10,000 two-tailed resamples, BH-FDR-corrected across the same four comparisons, all *p* ≥ 0.33; per-animal differences at most 0.025 in normalized FOV units). Violin plots of Euclidean distance from the normalized FOV center (0.5, 0.5); lines show the median and extrema. Analyses pool both cohorts (context-only n = 8, fear memory n = 9; Ctx-A post: 5,769 retained and 1,206 recruited cells; Ctx-B day1: 5,664 retained and 2,714 recruited cells). (**i**) Newly recruited cells were spatially interspersed with retained cells at chance levels in both groups, with no group difference (animal-level permutation tests on the mixing delta, 10,000 two-tailed resamples, BH-FDR-corrected across the two sessions. Diff is fear memory minus context-only. Ctx-A post: diff = +0.014; Ctx-B day1: diff = −0.007; both *p* = 0.74). For each recruited cell, the fraction of its k = 5 nearest neighbors (in normalized FOV space) that were retained cells gives a per-animal mixing score (near 1 = recruited cells surrounded by retained cells; lower = recruited cells clustering together). A null was generated by shuffling retained/recruited labels within each animal (1,000 iterations), and the mixing delta (observed − null mean) measures integration relative to chance given each animal’s retained-to-recruited ratio (positive = recruited cells interleaved with retained cells beyond chance; negative = recruited cells spatially segregated). In both groups and sessions, recruited cells were surrounded predominantly by retained cells (mixing score 0.64– 0.83) and the mixing delta was near zero (−0.028 to 0.004; mean |z| ≤ 0.03 against the within-animal null), indicating no spatial clustering.

**Figure S3:**
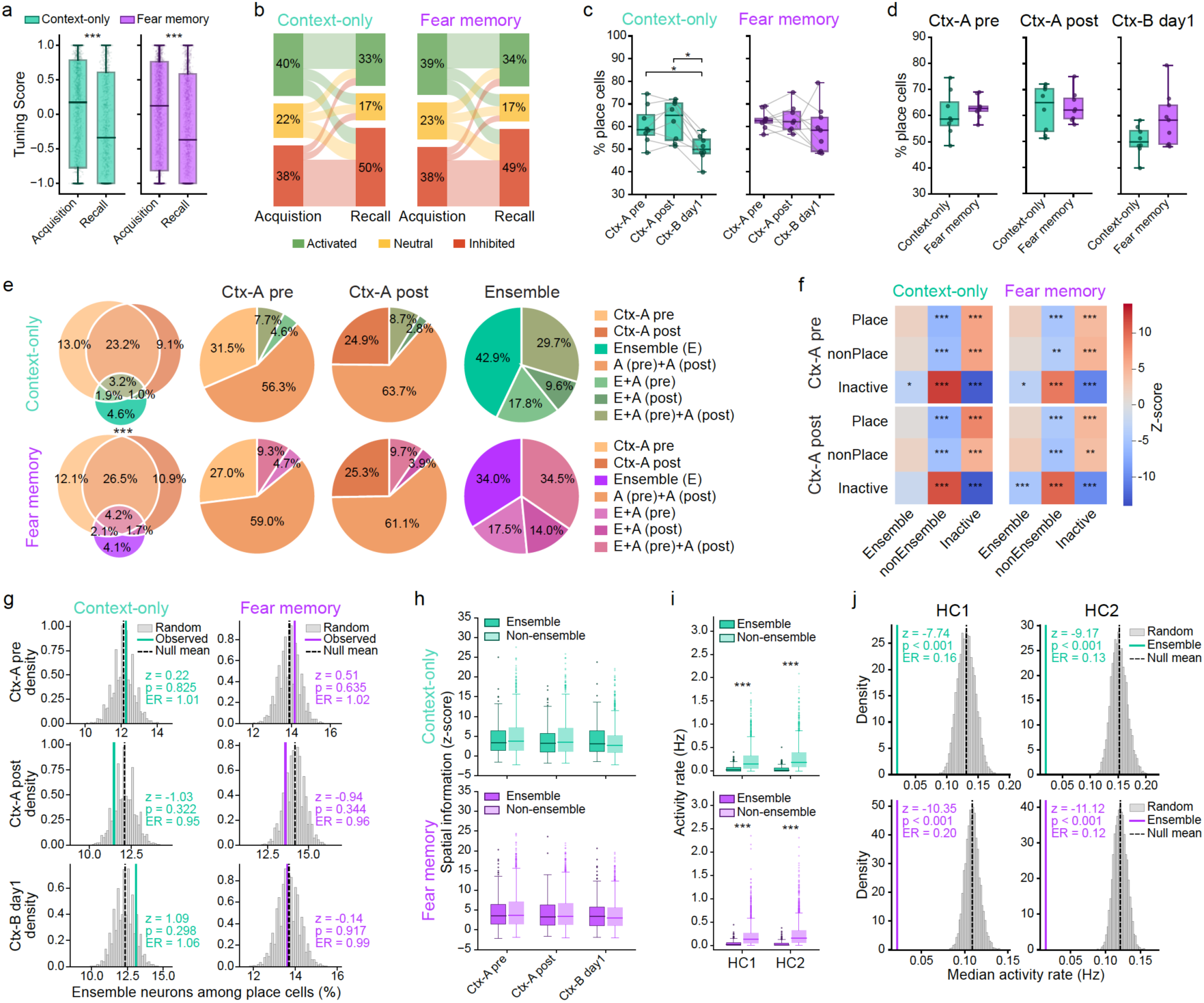
Characterization of memory ensemble and place cell populations. (**a**) Tuning scores were lower at recall than at acquisition in both groups, and did not differ between groups at either session (cell-level linear mixed models with animal as random intercept, REML; within-group session effects run separately per group, between-group effects BH-FDR-corrected across the two sessions. Within groups: context-only, coef = −0.216, *p* < 0.001, n = 3,769 cells from 4 animals; fear memory, coef = −0.217, *p* < 0.001, n = 5,447 cells from 5 animals. Between groups: acquisition, coef = +0.02, *p* = 0.97; recall, coef = −0.005, *p* = 0.97). The within-group decline was near-identical in magnitude in the two groups (−0.216 vs −0.217). Boxes show the cell-level distribution (median, interquartile range, whiskers to 1.5 × IQR, outliers not shown). (**b**) Transition of neurons between functional classes from acquisition to recall, restricted to cells detected in both sessions. Each node is a functional class: activated (green), neutral (yellow), or inhibited (red). Each flow represents the number of cells transitioning from their acquisition class (left) to their recall class (right); flow width is proportional to cell count. Only cells detected in both acquisition and recall are included; the Inactive class shown in **Figure 1e** therefore does not appear here. This complements **Figure 1e**, which uses the broader criterion of detection in at least one session. (**c**) The size of the cognitive map decreased at Ctx-B day1 in context-only animals but did not change across sessions in fear memory animals (animal-level paired permutation tests, 10,000 two-tailed resamples, BH-FDR-corrected across the three comparisons within each group. Δ is the earlier minus the later session, positive values indicating a smaller map at the later session. Context-only: Ctx-A pre → Ctx-A post, Δ = −2.05, *p* = 0.31; Ctx-A pre → Ctx-B day1, Δ = +10.21, *p* = 0.012; Ctx-A post → Ctx-B day1, Δ = +12.26, *p* = 0.012. Fear memory: Ctx-A pre → Ctx-A post, Δ = 0.00, *p* = 1.00; Ctx-A pre → Ctx-B day1, Δ = +4.42, *p* = 0.38; Ctx-A post → Ctx-B day1, Δ = +3.67, *p* = 0.68). Boxes show the group median and interquartile range, whiskers the full range; each dot is one animal (Context-only n = 8, Fear memory n = 9) and lines connect the same animal across sessions. Cognitive map size is the percentage of place cells among neurons detected in each session. (**d**) The percentage of place cells did not differ significantly between groups at any session (independent permutation tests on per-animal place cell percentages, 10,000 two-tailed resamples. Diff is fear memory minus context-only. Ctx-A pre: diff = +2.35, *p* = 0.48; Ctx-A post: diff = +0.73, *p* = 0.86; Ctx-B day1: diff = +8.14, *p* = 0.06). Each dot is one animal (context-only n = 8, fear memory n = 9); boxes show the group median and interquartile range, whiskers the full range. Values are the percentage of place cells per animal, among neurons detected in each exploration session. (**e**) Ctx-A pre place cells, Ctx-A post place cells, and ensemble neurons converge on partially overlapping neuronal populations (cell-level chi-square tests of independence. Overall distribution: χ²(6) = 14.35, *p* = 0.03; no individual population’s conditional distribution differed between groups, all *p* ≥ 0.06. At the animal level, the proportion of Ctx-A pre place cells also classified as Ctx-A post place cells was comparable between groups: context-only 63.8 ± 7.0% (n = 4), fear memory 67.0 ± 7.7% (n = 4), independent permutation test, diff = +3.2 percentage points, *p* = 0.57). Left, Venn diagram of the three populations; right, pie charts showing, within each population, the fraction that also belongs to the other two. Rows are the two groups (context-only, top; fear memory, bottom). The denominator is all cells detected in at least one of the sessions underlying these populations (Ctx-A pre and Ctx-A post for the place-cell populations, acquisition and recall for ensemble membership). One animal was excluded from this panel, as its Ctx-A post session did not pass quality control. Asterisks between the Venn diagrams indicate a significant difference in the overall overlap distribution between groups. (**f**) Both groups showed the same pattern of enrichment and depletion of spatial map classes within ensemble categories (within-animal Monte Carlo permutation). For each combination of ensemble category (Ensemble, nonEnsemble, Inactive) and exploration session (Ctx-A pre, Ctx-A post), we tested whether the fraction of cells also classified as Place, nonPlace or Inactive differed from chance given the marginal proportions. Spatial map labels were shuffled within each animal, 10,000 permutations, preserving per-animal cell counts and spatial class proportions, and aggregated across animals to form a population-level null. Z-scores give the observed value’s deviation from the null in s.d. units, positive = enrichment, negative = depletion; two-tailed permutation *p*-values are the fraction of shuffles at least as extreme as observed, Holm-corrected across the 18 tests per group, 2 sessions × 3 spatial classes × 3 ensemble categories). Per comparison, the denominator is all cells detected in at least one relevant session (acquisition, recall, or the exploration session), excluding globally silent cells. (**g**) Place cells were no more likely to be ensemble neurons than expected by chance, in any session in either group (within-animal shuffle test. For each session (Ctx-A pre, Ctx-A post, Ctx-B day1) the percentage of place cells that were also ensemble neurons was compared against a within-animal shuffled null, separately for each group. Nulls were generated by shuffling place cell labels within each animal, 10,000 permutations, preserving each animal’s cell count and place cell proportion, and aggregating overlap percentages across animals; two-tailed permutation *p*-values are the fraction of shuffles at least as extreme as observed. All |z| ≤ 1.09, all *p* ≥ 0.30, enrichment ratios 0.95–1.06). Grey histograms show the null distribution, the colored line the observed percentage, the dashed line the null mean. Insets give the z-score, *p*-value, and enrichment ratio (observed ÷ null mean; 1.0 = chance). Cells not detected in the exploration session are excluded. (**h**) Spatial information did not differ between ensemble and non-ensemble neurons in either group after correction (cell-level linear mixed models, z-score ∼ ensemble status with animal as random intercept, REML, BH-FDR-corrected across the three sessions within each group; ensemble is the reference level, so a positive coefficient indicates higher z-scores in non-ensemble neurons. Context-only: Ctx-A pre, coef = +0.61, *p* = 0.074; Ctx-A post, coef = +0.70, *p* = 0.074; Ctx-B day1, coef = −0.29, *p* = 0.26. Fear memory: all sessions *p* ≥ 0.31). Boxes show the single-cell z-score distribution (median, interquartile range, whiskers to 1.5 × IQR, outliers as points) for ensemble (dark) versus non-ensemble (light) neurons within each session and group. All cells detected in the relevant session are included (place and non-place cells). (**i**) Non-ensemble neurons were more active than ensemble neurons during home cage sessions, in both groups (cell-level linear mixed models, activity rate ∼ ensemble status with animal as random intercept, REML, BH-FDR-corrected across the two sessions within each group; ensemble is the reference level, so a positive coefficient indicates higher rates in non-ensemble neurons. Context-only: HC1, coef = +0.17; HC2, coef = +0.24. Fear memory: HC1, coef = +0.15; HC2, coef = +0.19. All *p* < 0.001). Boxes show the single-cell activity rate distribution (median, interquartile range, whiskers to 1.5 × IQR, outliers as dots) for ensemble (dark) versus non-ensemble (light) neurons within each group. HC1 and HC2 are the home cage sessions recorded immediately before acquisition and before recall, respectively. Statistical inference was performed with animal as a random intercept, whereas the boxes show the pooled cell-level distribution. (**j**) Ensemble neurons were less active than the population expectation during home-cage sessions, in both groups (within-animal Monte Carlo test. The observed statistic is each animal’s median ensemble-neuron activity rate, averaged across animals; nulls were generated by drawing, within each animal, the same number of neurons as that animal’s ensemble from its full recorded population, 10,000 iterations without replacement, taking the median of each draw and averaging across animals as for the observed statistic. Context-only: HC1, z = −7.74, enrichment ratio = 0.16; HC2, z = −9.17, ER = 0.13. Fear memory: HC1, z = −10.35, ER = 0.20; HC2, z = −11.12, ER = 0.12. All *p* < 0.001). Grey histograms show the null distribution, the colored line the observed statistic, the dashed line the null mean; insets give the z-score, enrichment ratio (observed ÷ null mean; 1.0 = chance), and exact two-tailed permutation p-value. HC1 and HC2 are the home-cage sessions before acquisition and before recall, as in **h**.

**Figure S4:**
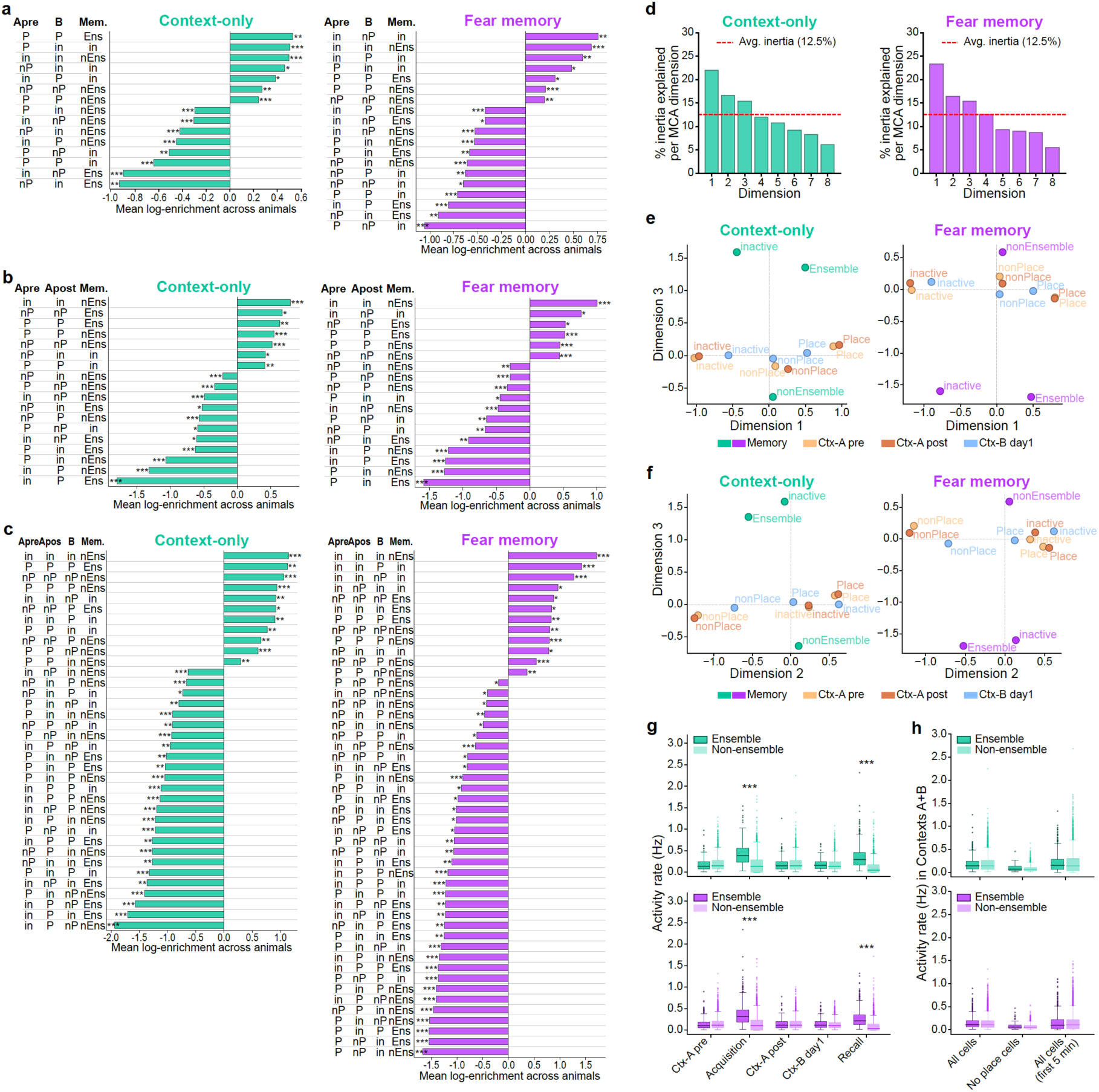
Supporting analyses for the memory-spatial profiling: MCA dimensionality and ensemble activity rates. (**a**) Significantly enriched and depleted co-occurrence profiles of ensemble identity and spatial-map class in Ctx-A pre and Ctx-B day1, shown separately for each group. Each neuron is assigned a profile combining memory status (Ensemble/Ens, nonEnsemble/nEns, inactive/in) with spatial map class in Ctx-A pre and Ctx-B day1 (Place/P, nonPlace/nP, inactive/in), displayed in the order Ctx-A pre (Apre), Ctx-B day1 (B), Memory (Mem.). Each bar is one significant profile; bars right of zero are enriched (occur more than expected by chance), left of zero depleted, with significance stars on the outer side of each bar. Bar length is the mean log-enrichment across animals, log((observed + 0.5) / (expected + 0.5)), with expected counts computed per animal under independence of memory status and spatial class and +0.5 Laplace smoothing for zero counts. Significance was assessed by permutation (10,000 iterations): condition labels shuffled independently within each animal, preserving marginal distributions, observed and expected counts recomputed, and the observed mean log-enrichment tested against the null; exact two-tailed *p*-values were BH-FDR-corrected across the 26 possible profiles (27 minus the all-inactive profile, which cannot occur given the denominator). Only profiles with *p* < 0.05 are shown. The denominator includes all neurons detected in at least one of the relevant sessions (Acquisition, Recall, and the two exploration sessions). (**b**) As in **a**, for Ctx-A pre and Ctx-A post (profiles ordered Apre, Apost, Mem.). (**c**) As in **a**, extend to four conditions: memory status and spatial map class in Ctx-A pre, Ctx-A post and Ctx-B day1 (profiles ordered Apre, Apost, B, Mem.). With four conditions × three labels, 81 combinations are possible; the all-inactive profile cannot occur, leaving 80, across which BH-FDR-correction is applied. Only profiles with *p* < 0.05 are shown. (**d**) Scree plots of the multiple correspondence analysis: percentage of total inertia explained by each MCA dimension, shown separately for context-only and fear memory animals. The red dashed line marks the average inertia per dimension (100% ÷ number of dimensions); dimensions above this threshold were retained for the clustering and compactness analysis in **Figure 2f**, yielding three dimensions for context-only and four for fear memory. (**e**) Multiple correspondence analysis of neuron profiles, dimensions 1 and 3. The 12-label condition points from the MCA in **Figure 2d**, projected onto MCA dimensions 1 and 3, shown separately for context-only and fear memory animals and colored by condition. As in the main figure, a point’s position reflects how strongly its label co-occurs with the others, so points lying close together tend to appear in the same neurons. (**f**) As in **e**, projected onto MCA dimensions 2 and 3. (**g**) Ensemble neurons were more active than non-ensemble neurons specifically during acquisition and recall, but not during the exploration sessions, in both groups (cell-level linear mixed models, activity rate ∼ ensemble status with animal as random intercept, REML, BH-FDR-corrected across the five sessions within each group; ensemble is the reference level, so a negative coefficient indicates higher rates in ensemble neurons. Context-only: Acquisition, coef = −0.231, *p* < 0.001; Recall, coef = −0.237, *p* < 0.001; Ctx-A pre, coef = +0.021, *p* = 0.15; Ctx-A post, coef = +0.016, *p* = 0.30; Ctx-B day1, coef = −0.002, *p* = 0.80. Fear memory: Acquisition, coef = −0.215, *p* < 0.001; Recall, coef = −0.183, *p* < 0.001; Ctx-A pre, coef = +0.014, *p* = 0.099; Ctx-A post, coef = +0.010, *p* = 0.23; Ctx-B day1, coef = −0.011, *p* = 0.068). Boxes show the cell-level distribution (median, interquartile range, whiskers to 1.5 × IQR, outliers as dots). Statistical inference was performed with animal as a random intercept, whereas the boxes show the pooled cell-level distribution. (**h**) The absence of an activity rate difference between ensemble and non-ensemble neurons during exploration was robust to spatial tuning and to novelty-induced arousal (cell-level linear mixed models, rate ∼ ensemble status with animal as random intercept, REML, BH-FDR-corrected across the three conditions within each group; ensemble is the reference level, so a negative coefficient indicates higher rates in ensemble neurons. Context-only: all cells, coef = +0.011, *p* = 0.15; non-place cells, coef = −0.015, *p* = 0.098; first 5 min, coef = +0.001, *p* = 0.81. Fear memory: all cells, coef = +0.003, *p* = 0.64; non-place cells, coef = −0.010, *p* = 0.23; first 5 min, coef = −0.001, *p* = 0.87). Three conditions are shown: all cells pooled across Ctx-A pre, Ctx-A post, and Ctx-B day1 (left); non-place cells only, never classified as place cells in any session (middle); and all cells restricted to the first 5 minutes of each recording (right), controlling respectively for place-cell spatial tuning and for novelty-induced arousal at placement. Activity rates were concatenated across sessions per cell, so each cell contributes up to three observations; the animal random intercept only partially accounts for this, and some pseudoreplication remains. Boxes show the cell-level distribution.

**Figure S5:**
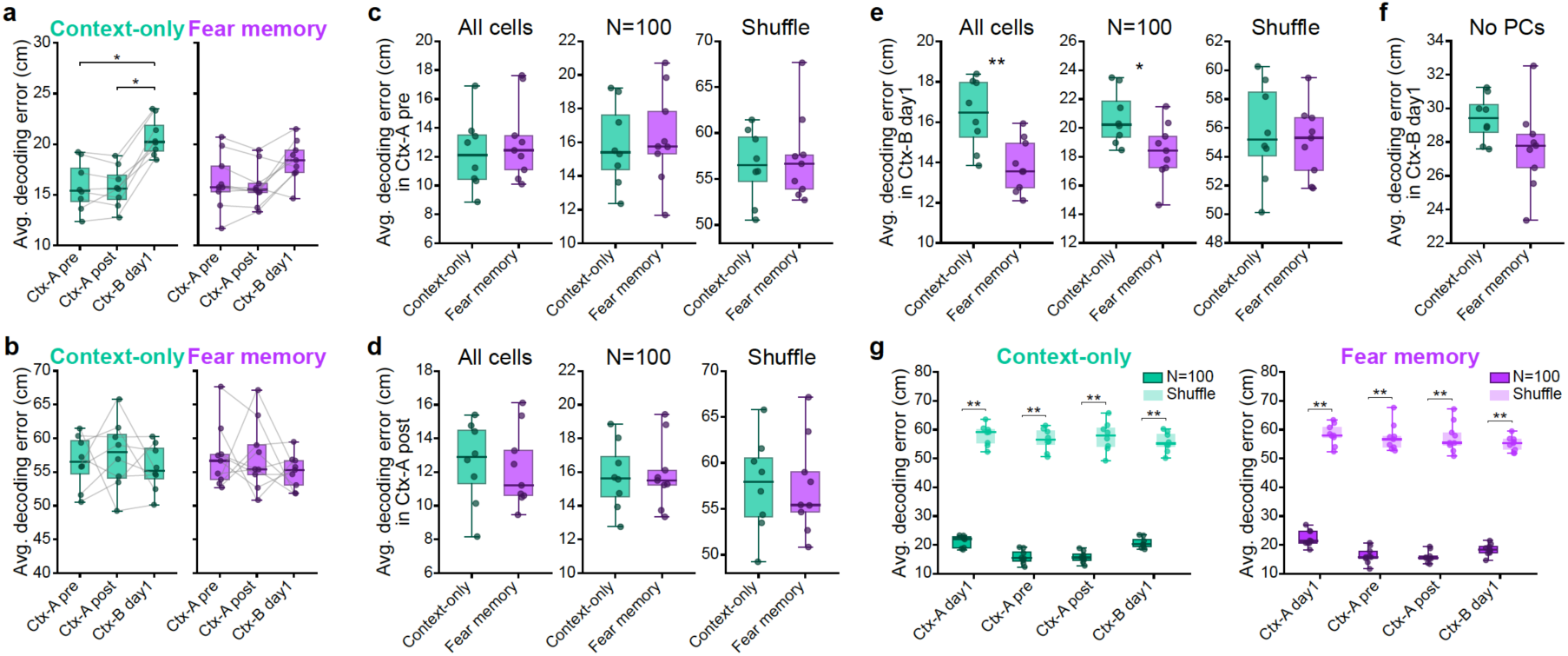
Population decoding of spatial precision across session and contexts, with subsampling and shuffle controls. (**a**) Bayesian population vector decoding error increased in the novel context in context-only animals but showed no significant change across Ctx-A pre, Ctx-A post and Ctx-B day1 in fear memory animals (paired permutation tests, 10,000 two-tailed resamples, BH-FDR-corrected across the three comparisons per group. Diff is the second minus the first session. Context-only: Ctx-A pre vs Ctx-A post, diff = −0.087, *p* = 0.82; Ctx-A pre vs Ctx-B day1, diff = +4.842, *p* = 0.012; Ctx-A post vs Ctx-B day1, diff = +4.930, *p* = 0.012. Fear memory: Ctx-A pre vs Ctx-A post, diff = −0.418, *p* = 0.24; Ctx-A pre vs Ctx-B day1, diff = +2.053, *p* = 0.076; Ctx-A post vs Ctx-B day1, diff = +2.471, *p* = 0.059). Decoding was performed on a random subsample of 100 neurons, shown separately for context-only and fear memory animals. Each box shows the group median and interquartile range of the per-animal median decoding error, whiskers the full range; dots are individual animals and lines connect the same animal across sessions. (**b**) Decoding error did not differ between Ctx-A pre, Ctx-A post and Ctx-B day1 in either group when position was circularly shuffled relative to neural activity (paired permutation tests, 10,000 two-tailed resamples, BH-FDR-corrected across the three comparisons per group. Context-only: Ctx-A pre vs Ctx-A post, diff = +1.045; Ctx-A pre vs Ctx-B day1, diff = −0.851; Ctx-A post vs Ctx-B day1, diff = −1.896; all *p* = 0.73. Fear memory: Ctx-A pre vs Ctx-A post, diff = +0.097, *p* = 0.97; Ctx-A pre vs Ctx-B day1, diff = −2.233, *p* = 0.53; Ctx-A post vs Ctx-B day1, diff = −2.330, *p* = 0.53). Circular shuffling preserves the temporal structure of both signals while destroying their alignment, giving the chance-level decoding error. Layout as in **a**. (**c**) Bayesian population vector decoding error in Ctx-A pre did not differ between context-only and fear memory animals under any decoding condition (independent permutation tests, 10,000 two-tailed resamples. Diff is fear memory minus context-only. All detected neurons: diff = +0.820, *p* = 0.52; random subsample of 100 neurons: diff = +0.470, *p* = 0.70; shuffle control: diff = +0.768, *p* = 0.73). Three decoding conditions are shown: all detected neurons (left), a random subsample of 100 neurons (middle), and a shuffle control in which position labels were circularly shuffled relative to neural activity (right), which provides an estimate of chance-level decoding error. Each box shows the group median and interquartile range of the per-animal median decoding error, whiskers the full range; dots are individual animals. (**d**) Bayesian population vector decoding error in Ctx-A post did not differ between context-only and fear memory animals under any decoding condition (independent permutation tests, 10,000 two-tailed resamples. All detected neurons: diff = −0.366, *p* = 0.76; random subsample of 100 neurons: diff = +0.140, *p* = 0.87; shuffle control: diff = −0.180, *p* = 0.94). Layout and statistics as in **c**. (**e**) In Ctx-B day1, fear memory animals showed lower decoding error than context-only animals when decoding from all detected neurons and from a random subsample of 100 neurons, with no difference in the shuffle control (independent permutation tests, 10,000 two-tailed resamples. All detected neurons: diff = −2.533, *p* = 0.006; random subsample of 100 neurons: diff = −2.319, *p* = 0.027; shuffle control: diff = −0.614, *p* = 0.68). Layout and statistics as in **c**. (**f**) Removing all place cells reduced but did not abolish the difference in Bayesian population vector decoding error between context-only and fear memory animals in Ctx-B day1, which remained in the same direction as when place cells were included (independent permutation test, 10,000 two-tailed resamples. Diff = −1.769, *p* = 0.10, compared with diff = −2.319 for the random subsample of 100 neurons in **e**. Decoding was performed on a random subsample of 100 non-place cells. Layout and statistics as in **c**. (**g**) In both groups, decoding error from a random subsample of 100 neurons was far below the shuffle baseline in every session, and varied across Ctx-A day1, Ctx-A pre, Ctx-A post and Ctx-B day1 in a way that the shuffle condition did not (paired permutation tests, 10,000 two-tailed resamples, BH-FDR-corrected across the four sessions for the within-session comparisons and across all six pairwise combinations for the cross-session comparisons, separately per group and condition. Diff is the second minus the first session. Context-only: N = 100 below shuffle in all sessions, all diff > 34, all *p* = 0.008. Fear memory: all diff > 36, all *p* = 0.004). In context-only animals, decoding error was higher in Ctx-A day1 than in Ctx-A pre (diff = −5.327, *p* = 0.012) and Ctx-A post (diff = −5.415, *p* = 0.012), and higher in Ctx-B day1 than in Ctx-A pre (diff = +4.842, *p* = 0.012) and Ctx-A post (diff = +4.930, *p* = 0.012), with no difference between Ctx-A day1 and Ctx-B day1 (diff = −0.485, *p* = 0.64) or between Ctx-A pre and Ctx-A post (diff = −0.087, *p* = 0.82). In fear memory animals, decoding error was higher in Ctx-A day1 than in Ctx-A pre (diff = −5.883, *p* = 0.012), Ctx-A post (diff = −6.301, *p* = 0.012) and Ctx-B day1 (diff = −3.830, *p* = 0.016), and higher in Ctx-B day1 than in Ctx-A post (diff = +2.471, *p* = 0.029), while Ctx-A pre did not differ from Ctx-A post (diff = −0.418, *p* = 0.24) or from Ctx-B day1 (diff = +2.053, *p* = 0.061). Shuffle decoding error did not differ across sessions in either group (Context-only: all diff < 2.41, all *p* > 0.68. Fear memory: all diff < 3.24, all *p* > 0.47). N = 100 (dark) is decoding from a random subsample of 100 neurons; shuffle (light) is decoding after circularly shifting position relative to neural activity, giving the chance-level baseline. Each box shows the group median and interquartile range of the per-animal median decoding error, whiskers the full range; dots are individual animals.

**Figure S6:**
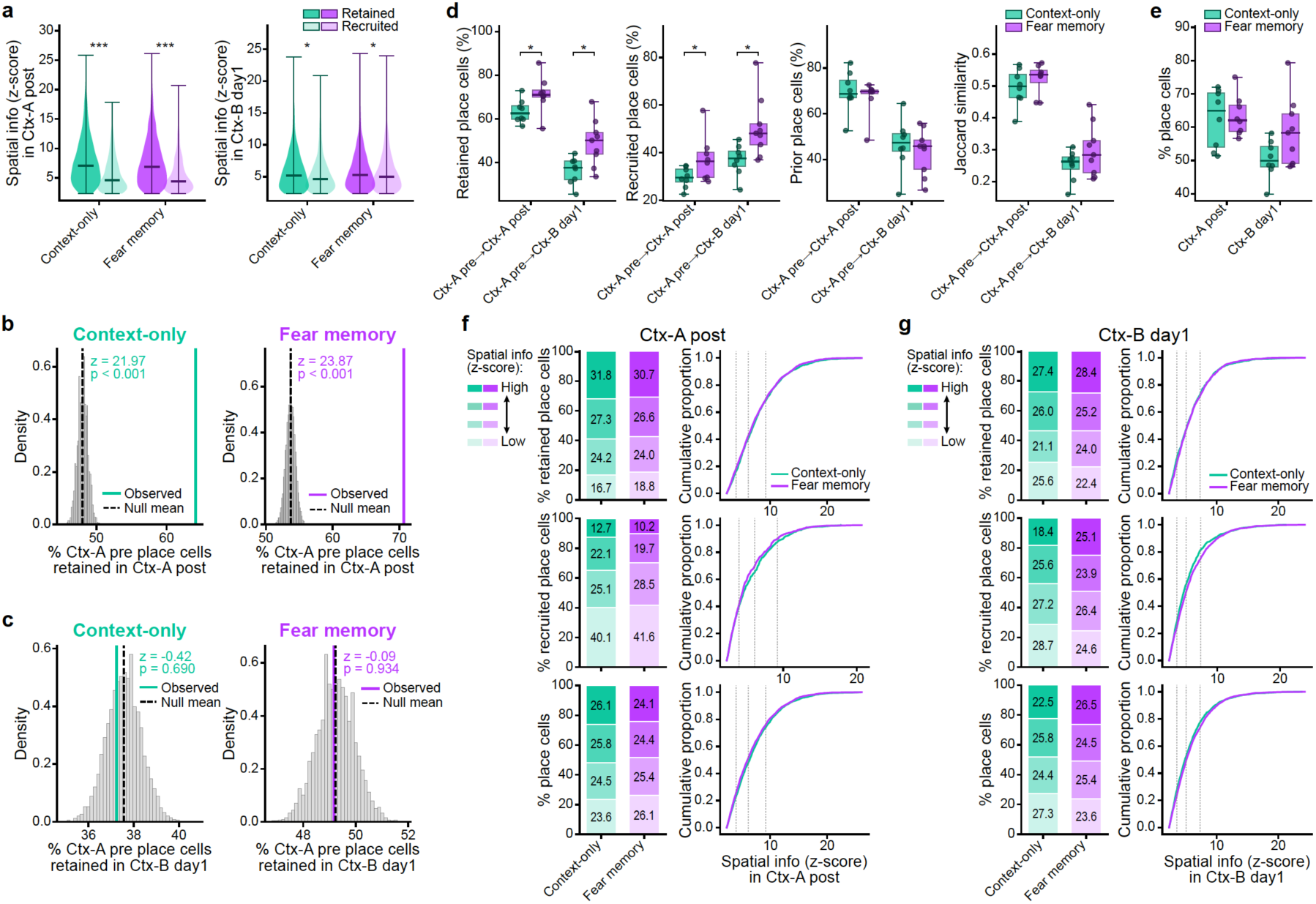
Retention and recruitment of place cells across repeated exposures to a familiar context and on entry to a novel context. (**a**) Place cells retained from Ctx-A pre carried more spatial information than newly recruited place cells in both Ctx-A post and Ctx-B day1, in both groups (cell-level linear mixed models, spatial information z-score ∼ cell type, animal as random intercept, REML, BH-FDR-corrected across the two groups. Coefficients are retained minus recruited. Ctx-A post: context-only, coef = +2.284, *p* < 0.001; fear memory, coef = +2.375, *p* < 0.001. Ctx-B day1: context-only, coef = +0.407, *p* = 0.017; fear memory, coef = +0.235, *p* = 0.048). The difference was substantially smaller in Ctx-B day1 than in Ctx-A post. Place cells in Ctx-A post (left) and Ctx-B day1 (right) are classified as retained if they were also place cells in Ctx-A pre, and as newly recruited if they were not; the spatial information z-score of that session is compared between these two populations. Darker violins indicate retained cells, lighter violins recruited cells. (**b**) Place cell identity was retained from Ctx-A pre to Ctx-A post far more often than expected by chance in both groups (Context-only: z = 21.97, *p* < 0.001. Fear memory: z = 23.87, *p* < 0.001). Among cells classified as place cells in Ctx-A pre, the histogram shows the observed fraction also classified as place cells in Ctx-A post (colored line) against a within-animal null (grey histogram; dashed line, null mean) generated by shuffling Ctx-A post spatial class labels independently within each animal (10,000 permutations, preserving each animal’s spatial class proportions). Z-scores and exact two-tailed permutation *p*-values are shown per panel, for context-only (left) and fear memory (right) animals. (**c**) Retention of place cell identity from Ctx-A pre to Ctx-B day1 was indistinguishable from chance in both groups (Context-only: z = −0.42, *p* = 0.69. Fear memory: z = −0.09, *p* = 0.93). As in **b**, for retention from Ctx-A pre to Ctx-B day1: among Ctx-A pre place cells, the fraction also classified as place cells in Ctx-B day1, compared against a within-animal shuffled null. (**d**) Fear memory animals showed higher retention of place cell identity and higher recruitment of new place cells than context-only animals, from Ctx-A pre to both Ctx-A post and Ctx-B day1, whereas the proportion of prior place cells and Jaccard similarity did not differ (independent permutation tests, 10,000 two-tailed resamples, BH-FDR-corrected across the two session pairs per metric. Diff is fear memory minus context-only. Ctx-A pre → Ctx-A post: retention rate, diff = +7.937, *p* = 0.037; recruitment rate, diff = +7.715, *p* = 0.038; reciprocal retention, diff = −2.495, *p* = 0.56; Jaccard index, diff = +0.026, *p* = 0.35. Ctx-A pre → Ctx-B day1: retention rate, diff = +12.947, *p* = 0.014; recruitment rate, diff = +13.672, *p* = 0.012; reciprocal retention, diff = −3.349, *p* = 0.56; Jaccard index, diff = +0.047, *p* = 0.33). Four metrics are shown: retained place cells, the percentage of Ctx-A pre place cells also classified as place cells in the second session; recruited place cells, the percentage of cells that were non-place or not detected in Ctx-A pre and became place cells in the second session; prior place cells, the percentage of second-session place cells that were also place cells in Ctx-A pre; and the Jaccard index, the intersection of the two sessions’ place cell populations divided by their union, giving a symmetric measure of population overlap. Boxes show the group median and interquartile range, whiskers the full range, and each dot is one animal. (**e**) The percentage of place cells did not differ between groups in either Ctx-A post or Ctx-B day1 (independent permutation tests, 10,000 two-tailed resamples, BH-FDR-corrected across the two sessions. Diff is fear memory minus context-only. Ctx-A post: diff = +0.729, *p* = 0.84. Ctx-B day1: diff = +8.142, *p* = 0.109). Statistics and layout as in **d**. (**f**) For the Ctx-A pre → Ctx-A post session pair, the distribution of spatial information z-scores across quartiles did not differ between context-only and fear memory animals for any place cell type (cell-level chi-square tests of independence, all *p* > 0.13; animal-level independent permutation tests on per-animal quartile proportions, BH-FDR-corrected across the four quartiles, all *p* > 0.32). Quartile boundaries were computed from the pooled place cell distribution across both groups and both cell types. Three rows are shown: retained place cells (place cells in both Ctx-A pre and Ctx-A post), newly recruited place cells (place cells in Ctx-A post only), and all place cells combined. Stacked bars (left) show the proportion of cells in each spatial information quartile (Q1 low to Q4 high) per group. Cumulative distribution functions (right) show the distribution of spatial information z-scores per group with quartile boundaries marked (dotted lines), shown for visualization only. The chi-square tests are exploratory, as cells from the same animal are not independent. (**g**) For the Ctx-A pre → Ctx-B day1 session pair, quartile distributions differed between groups at the cell level for newly recruited place cells (χ² = 16.3, df = 3, *p* = 0.001) and for all place cells combined (χ² = 13.8, df = 3, *p* = 0.003) but not for retained place cells (χ² = 4.11, df = 3, *p* = 0.25), whereas no per-animal quartile proportions survived FDR correction for any cell type (all *p* > 0.18). Layout and statistics as in **f**, with retained defined as place cells in both Ctx-A pre and Ctx-B day1, and recruited as place cells in Ctx-B day1 only.

**Figure S7:**
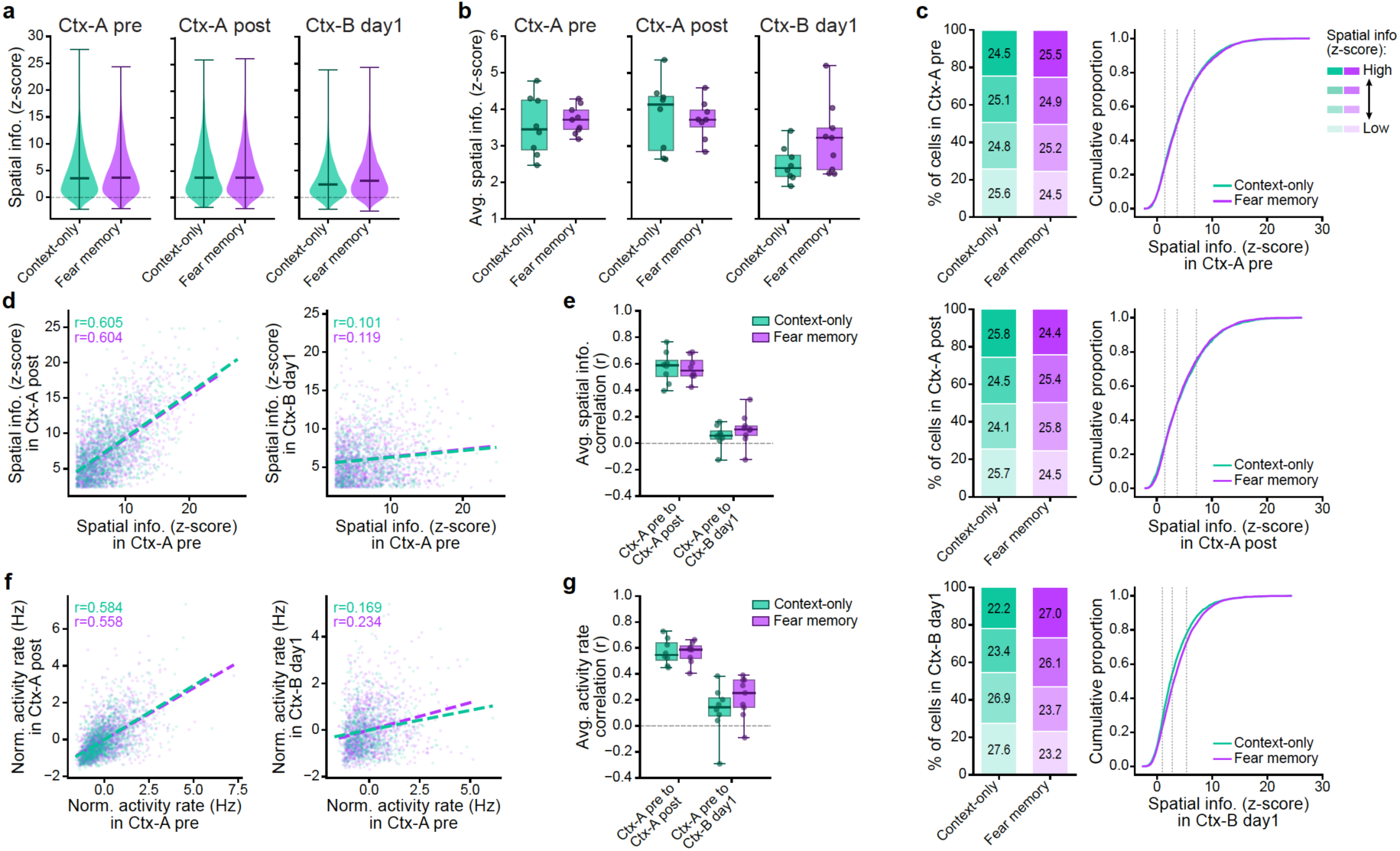
Population spatial information and cross-session stability of place cell tuning levels. (**a**) Across all detected cells, the cell-level distribution of spatial information z-scores did not differ significantly between context-only and fear memory animals in any session (cell-level linear mixed models, z-score ∼ group, animal as random intercept, REML, one per session. Coefficient is fear memory minus context-only. Ctx-A pre: coef = +0.166, *p* = 0.59. Ctx-A post: coef = −0.060, *p* = 0.87. Ctx-B day1: coef = +0.525, *p* = 0.13. Violins show the cell-level distribution per group across all detected cells, with the horizontal line marking the median. (**b**) Per-animal median spatial information z-scores did not differ significantly between groups in any session (independent permutation tests on per-animal medians, 10,000 two-tailed resamples, one per session. Diff is fear memory minus context-only. Ctx-A pre: 3.71 ± 0.36 vs 3.55 ± 0.77, diff = +0.16, *p* = 0.60. Ctx-A post: 3.72 ± 0.51 vs 3.83 ± 0.92, diff = −0.11, *p* = 0.77. Ctx-B day1: 3.20 ± 0.91 vs 2.50 ± 0.46, diff = +0.69, *p* = 0.084). Boxes show the group median and interquartile range, whiskers the full range, and each dot is one animal. (**c**) Quartile and cumulative distributions of spatial information z-scores per group. In Ctx-A pre and Ctx-A post the distributions did not differ between groups (Ctx-A pre: χ² = 1.89, *p* = 0.60; KS = 0.020, *p* = 0.46. Ctx-A post: χ² = 4.75, *p* = 0.19; KS = 0.028, *p* = 0.15; all per-animal quartile proportions *p* > 0.93). In Ctx-B day1, both the cell-level chi-square (χ² = 48.6, df = 3, *p* < 0.001) and the KS test (KS = 0.080, *p* < 0.001) were significant: fear memory animals had relatively more cells in the high quartile (27.8% vs 22.7%) and fewer in the low quartile (22.3% vs 26.8%), a small rightward shift in spatial information. No per-animal quartile proportion survived FDR correction in any session (Ctx-B day1: all *p* > 0.11). Quartile boundaries were computed per session from the pooled distribution across both groups. Stacked bars show the proportion of cells per quartile per group, tested by cell-level chi-square of independence and by animal-level permutation tests on per-animal quartile proportions (BH-FDR-corrected across the four quartiles). The CDF panels show the cumulative distribution per group, with quartile boundaries marked; the chi-square and KS tests are exploratory, as cells from the same animal are not independent. (**d**) Cell-level correlation of place cell spatial information z-scores across sessions, for context-only and fear memory animals. Scatter plots show the spatial information z-score in session 1 versus session 2, pooled across animals, with regression lines per group; Pearson r values are computed from pooled cell-level data and shown for descriptive purposes only, as cells from the same animal are not independent. Only cells classified as place cells in both sessions are included. (**e**) Per-animal correlations of place cell spatial information z-scores were substantially lower across contexts than across repeated sessions in the same context, in both groups, and did not differ between groups for either session pair (paired sign-flip permutation test on per-animal r values for the within-versus-across comparison, and independent permutation tests for the between-group comparison, 10,000 two-tailed resamples, BH-FDR-corrected. Within versus across: context-only, mean Δ = −0.522, *p* = 0.009; fear memory, mean Δ = −0.452, *p* = 0.009. Between groups: Ctx-A pre → Ctx-A post, context-only r = 0.57 vs fear memory r = 0.56, *p* = 0.82; Ctx-A pre → Ctx-B day1, context-only r = 0.05 vs fear memory r = 0.10, *p* = 0.69). Δ is the across-context minus the within-context correlation. Each dot is one animal’s Pearson r for that session pair. Inclusion criteria as in **d**. Boxes show the group median and interquartile range, whiskers the full range. (**f**) Cell-level correlation of normalized place cell activity rates across sessions, for context-only and fear memory animals. Scatter plots show the normalized activity rate in session 1 versus session 2, pooled across animals; with regression lines per group; Pearson r values are computed from pooled cell-level data and shown for descriptive purposes only, as cells from the same animal are not independent. Only cells classified as place cells in both sessions are included. Activity rates were z-score normalized within each animal before computing correlations, removing animal-specific differences in overall activity level. (**g**) Per-animal correlations of normalized place cell activity rates were likewise substantially lower across contexts than across repeated sessions in the same context, in both groups, and did not differ between groups for either session pair (statistics as in **e**. Within versus across: context-only, mean Δ = −0.449, *p* = 0.009; fear memory, mean Δ = −0.307, *p* = 0.009. Between groups: Ctx-A pre → Ctx-A post, context-only r = 0.57 vs fear memory r = 0.56, *p* = 0.91; Ctx-A pre → Ctx-B day1, context-only r = 0.12 vs fear memory r = 0.22, *p* = 0.58. Each dot is one animal’s Pearson r for that session pair. Inclusion criteria and normalization as in **f**. Boxes show the group median and interquartile range, whiskers the full range.

**Figure S8:**
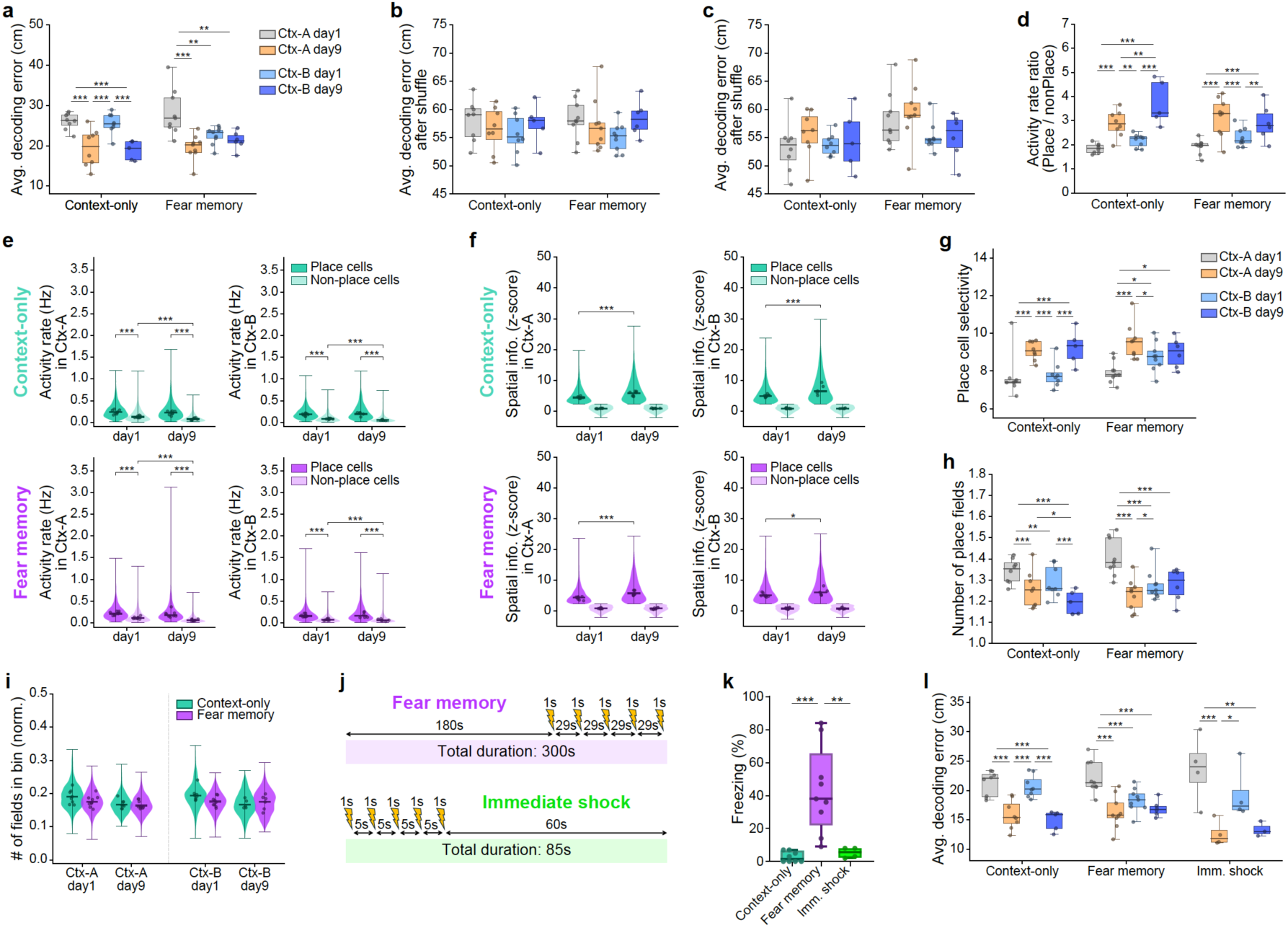
Within-session map refinement across familiarization: additional measures and controls. (**a**) Under a sequential (rather than random) train/test split, Bayesian population vector decoding error also decreased from the first to the last day of familiarization in both contexts in context-only animals, but in fear memory animals this improvement was absent in Ctx-B (statistical approach as in **Figure 4a**. Δ is in cm, positive values indicating lower error on the later session. Context-only: Ctx-A day1 → day9, Δ = +6.60, *p* < 0.001; Ctx-B day1 → day9, Δ = +5.86, *p* < 0.001; Ctx-A day9 → Ctx-B day1, Δ = −6.11, *p* < 0.001; Ctx-A day1 → Ctx-B day9, Δ = +6.26, *p* < 0.001. Fear memory: Ctx-A day1 → day9, Δ = +8.62, *p* < 0.001; Ctx-A day1 → Ctx-B day1, Δ = +5.69, *p* = 0.002; Ctx-A day1 → Ctx-B day9, Δ = +8.34, *p* = 0.001; Ctx-B day1 → day9, Δ = +0.73, *p* = 0.52, not significant). The absence of improvement across Ctx-B in fear memory animals contrasts with the corresponding improvement in Ctx-A of the same animals (Δ = +8.62) and with Ctx-B in context-only animals (Δ = +5.86). All remaining contrasts had *p* ≥ 0.14. Because the split is sequential (training on the first 80% mobile frames of each session and testing on the last 20%), this analysis additionally reflects within-session stability of the map, not only decoding precision. Each dot is one animal; boxes show the group median and interquartile range, whiskers the full range. (**b**) With position circularly shuffled relative to neural activity, decoding error did not differ across sessions in either group under the random split, confirming that the session differences in the unshuffled analysis (**Figure 4a**) reflect genuine spatial coding (statistical approach as in **Figure 4a**. No contrast reached significance: context-only, all *p* ≥ 0.76; fear memory, all *p* ≥ 0.25). Decoding used a random split of train and test frames during mobility, with position circularly shuffled to give the chance-level baseline. Each dot is one animal; boxes show the group median and interquartile range, whiskers the full range. (**c**) The same as in **b** held under the sequential split: with position circularly shuffled, decoding error did not differ across sessions in either group (statistical approach as in **Figure 4a**. No contrast reached significance: context-only, all *p* ≥ 0.83; fear memory, all *p* ≥ 0.16). Layout as in **a**, with position circularly shuffled to give the chance-level baseline. Each dot is one animal; boxes show the group median and interquartile range, whiskers the full range. (**d**) The ratio of place cell to non-place cell median activity rate increased from the first to the last day of familiarization across Ctx-A and Ctx-B in both groups, indicating that place cells became increasingly selective relative to the background population (statistical approach as in **Figure 4a**. Δ is in ratio units, negative values indicating a higher ratio on the later session. Context-only: Ctx-A day1 → day9, Δ = −1.03, *p* < 0.001; Ctx-B day1 → day9, Δ = −1.55, *p* < 0.001; Ctx-A day9 → Ctx-B day1, Δ = +0.69, *p* = 0.007; Ctx-A day9 → Ctx-B day9, Δ = −1.04, *p* = 0.004; Ctx-A day1 → Ctx-B day9, Δ = −1.84, *p* < 0.001. Fear memory: Ctx-A day1 → day9, Δ = −1.23, *p* < 0.001; Ctx-A day9 → Ctx-B day1, Δ = +0.86, *p* < 0.001; Ctx-A day1 → Ctx-B day9, Δ = −1.09, *p* < 0.001; Ctx-B day1 → day9, Δ = −0.63, *p* = 0.003). The cross-context drop in the ratio from Ctx-A day9 to Ctx-B day1 was significant in both groups (context-only Δ = +0.69; fear memory Δ = +0.86). The increase in this ratio across Ctx-B was present in both groups. All remaining contrasts had *p* ≥ 0.060. Each dot is one animal; boxes show the group median and interquartile range, whiskers the full range. (**e**) Place cells were more active than non-place cells at both day 1 and day 9 of each context, in both groups; across days, non-place cell activity rates decreased while place cel rates were unchanged (animal-level linear mixed-effects models on per-animal medians, with cell type or day as a categorical fixed effect and animal as a random intercept, REML; four comparisons per panel BH-FDR-corrected. In every panel: place vs non-place at day1 and at day9, both *p* < 0.001; non-place cells day1 vs day9, *p* < 0.001; place cells day1 vs day9, non-significant in all four panels, *p* = 0.30, 0.45, 0.20 and 0.62 for context-only Ctx-A, context-only Ctx-B, fear memory Ctx-A and fear memory Ctx-B respectively). Activity rates are show at day1 and day9 of Ctx-A (left) and Ctx-B (right), for context-only (top) and fear memory (bottom) animals. Violins show the full distributions across all cells pooled within each group; dots show the per-animal median. Dark violins are place cells, light violins are non-place cells. Per-animal medians were used as model input to account for the unequal numbers of place and non-place cells per animal; cell-level linear mixed models (one observation per cell, animal as random intercept) were run as a sanity check and agreed with the animal-level tests except for the place cell day1 vs day9 comparison, which reached significance at the cell level in three of the four panels (all but context-only Ctx-A) while remaining non-significant at the animal level, reflecting the greater power of the cell-level test. (**f**) Spatial information of place cells increased from the first to the last day of familiarization in both contexts and both groups, whereas spatial information of non-place cells was unchanged (animal-level linear mixed-effects models on per-animal medians, with day as a categorical fixed effect and animal as a random intercept, REML; two comparisons per panel BH-FDR-corrected. Place cells day1 vs day9: context-only Ctx-A, *p* < 0.001; context-only Ctx-B, *p* < 0.001; fear memory Ctx-A, *p* < 0.001; fear memory Ctx-B, *p* = 0.020. Non-place cells day1 vs day9: non-significant in all four panels, *p* = 0.64, 0.97, 0.25 and 0.62 respectively). Spatial information z-scores are shown at day1 and day9 of Ctx-A (left) and Ctx-B (right), for context-only (top) and fear memory (bottom) animals. Violins show the full distribution across all cells pooled within each group; dots show the per-animal median. Dark violins are place cells, light violins non-place cells. Place and non-place cells are defined by a spatial information z-score threshold, so the two populations are not compared within a session; only the day 1 versus day 9 chang within each population is tested. Per-animal medians were used as model input to account for the unequal numbers of place and non-place cells per animal, and cell-level linear mixed models (one observation per cell, animal as random intercept) run as a sanity check agreed with the animal-level tests throughout. (**g**) Place cell selectivity increased from the first to the last day of familiarization in both contexts in context-only animals, but in fear memory animals this increase was absent in Ctx-B (statistical approach as in **Figure 4a**. Δ is in selectivity units, negative values indicating higher selectivity on the later session. Context-only: Ctx-A day1 → day9, Δ = −1.35, *p* < 0.001; Ctx-B day1 → day9, Δ = −1.15, *p* < 0.001; Ctx-A day9 → Ctx-B day1, Δ = +1.28, *p* < 0.001; Ctx-A day1 → Ctx-B day9, Δ = −1.18, *p* < 0.001. Fear memory: Ctx-A day1 → day9, Δ = −1.60, *p* < 0.001; Ctx-A day1 → Ctx-B day1, Δ = −0.82, *p* = 0.025; Ctx-A day9 → Ctx-B day1, Δ = +0.78, *p* = 0.026; Ctx-A day1 → Ctx-B day9, Δ = −0.93, *p* = 0.020; Ctx-B day1 → day9, Δ = −0.04, *p* = 0.61, not significant). The absence of an increase across Ctx-B in fear memory animals contrasts with the corresponding increase in Ctx-A of the same animals (Δ = −1.60) and with Ctx-B in context-only animals (Δ = −1.15). All remaining contrasts had *p ≥* 0.13. Each dot is one animal; boxes show the group median and interquartile range, whiskers the full range. (**h**) The median number of place fields per place cell decreased from the first to the last day of familiarization in both contexts in context-only animals, but in fear memory animals this decrease was absent in Ctx-B (statistical approach as in **Figure 4a**. Δ is in fields per cell, positive values indicating fewer fields on the later session. Context-only: Ctx-A day1 → day9, Δ = +0.084, *p* < 0.001; Ctx-B day1 → day9, Δ = +0.083, *p* < 0.001; Ctx-A day1 → Ctx-B day1, Δ = +0.052, *p* = 0.008; Ctx-A day9 → Ctx-B day9, Δ = +0.050, *p* = 0.013; Ctx-A day1 → Ctx-B day9, Δ = +0.147, *p* < 0.001. Fear memory: Ctx-A day1 → day9, Δ = +0.176, *p* < 0.001; Ctx-A day1 → Ctx-B day1, Δ = +0.130, *p* < 0.001; Ctx-A day9 → Ctx-B day1, Δ = −0.046, *p* = 0.050; Ctx-A day1 → Ctx-B day9, Δ = +0.159, *p* < 0.001; Ctx-B day1 → day9, Δ = +0.015, *p* = 0.50, not significant). The absence of a decrease across Ctx-B in fear memory animals contrasts with the corresponding decrease in Ctx-A of the same animals (Δ = +0.176) and with Ctx-B in context-only animals (Δ = +0.083). All remaining contrasts had *p* ≥ 0.085. Each dot is one animal; boxes show the group median and interquartile range, whiskers the full range. (**i**) The number of place fields per spatial bin did not differ between groups at any timepoint (animal-level permutation tests on per-animal medians, independent samples, 10,000 permutations, BH-FDR-corrected across the four timepoints. Ctx-A day1, *p* = 0.33; Ctx-A day9, *p* = 0.69; Ctx-B day1, *p* = 0.071; Ctx-B day9, *p* = 0.69). The largest difference, at Ctx-B day1, was nominally significant before correction (*p* = 0.018) but did not survive it. Violins show the full distribution of spatial bins pooled within each group; dots show the per-animal median. Cell-level linear mixed-effects models with group as a categorical fixed effect (reference: context-only) and animal as a random intercept (REML), which account for the nested structure of bins within animals, were run as a sanity check and gave the same pattern throughout (Ctx-B day1, *p* = 0.019 uncorrected; all others non-significant). (**j**) Schematic of the immediate shock protocol compared with the standard fear conditioning protocol. In the standard protocol, animals explored the conditioning context for three minutes before receiving the first footshock, allowing a context representation to be formed and associated with the shock. In the immediate shock protocol, the first footshock was delivered within two seconds of placement, leaving insufficient time to form a context representation, so animals receive the same aversive stimulus without acquiring a contextual fear memory. (**k**) Fear memory animals froze substantially more than both context-only and immediate shock animals, which did not differ detectably from each other, confirming that the footshocks alone are not sufficient to produce contextual freezing without prior exploration of the context (omnibus permutation test across the three groups, 10,000 label shuffles, *p* = 0.001; pairwise permutation tests, 10,000 two-tailed resamples, BH-FDR-corrected across the three comparisons. Fear memory 43.2 ± 25.9%, n = 9; context-only 2.8 ± 3.2%, n = 8; immediate shock 5.3 ± 3.2%, n = 4. Fear memory vs context-only, diff = +40.5, *p* < 0.001; fear memory vs immediate shock, diff = +38.0, *p* = 0.004; context-only vs immediate shock, diff = −2.5, *p* = 0.25). Diff is the first minus the second group, in percentage points. Values are mean ± s.d. Each dot is one animal; boxes show the group median and interquartile range, whiskers the full range. (**l**) With the immediate shock group included, decoding error decreased across Ctx-A familiarization in all three groups. Across Ctx-B familiarization it decreased in context-only animals, while the comparison did not reach significance in either fear memory or immediate shock animals. These two null results differ in kind: in fear memory animals the effect was near zero despite adequate sampling (Δ = +0.84 cm, dz = 0.60, n = 6, *p* = 0.21), whereas in immediate shock animals the improvement was of comparable magnitude and effect size to that in context-only animals (Δ = +3.73 cm, dz = 2.04, n = 3, *p* = 0.077, versus Δ = +4.84 cm, dz = 4.40, n = 5, *p* < 0.001 in context-only) but did not reach significance with only three animals contributing (statistical approach as in **Figure 4a**. Δ is in cm, positive values indicating lower error on the later session. Context-only: Ctx-A day1 → day9, Δ = +5.33, *p* < 0.001; Ctx-B day1 → day9, Δ = +4.84, *p* < 0.001; Ctx-A day9 → Ctx-B day1, Δ = −4.84, *p* < 0.001; Ctx-A day1 → Ctx-B day9, Δ = +5.16, *p* < 0.001. Fear memory: Ctx-A day1 → day9, Δ = +5.88, *p* < 0.001; Ctx-A day1 → Ctx-B day1, Δ = +3.83, *p* < 0.001; Ctx-A day1 → Ctx-B day9, Δ = +5.92, *p* < 0.001; Ctx-B day1 → day9, Δ = +0.84, *p* = 0.21, not significant. Immediate shock: Ctx-A day1 → day9, Δ = +11.05, *p* < 0.001, n = 4; Ctx-A day9 → Ctx-B day1, Δ = −6.77, *p* = 0.012, n = 4; Ctx-A day1 → Ctx-B day9, Δ = +9.89, *p* = 0.0014, n = 3; Ctx-B day1 → day9, Δ = +3.73, *p* = 0.077, n = 3). All remaining contrasts had *p ≥* 0.050. Group sizes for the immediate shock group are given per contrast, as one animal did not complete Ctx-B day9. Each dot is one animal; boxes show the group median and interquartile range, whiskers the full range.

**Figure S9:**
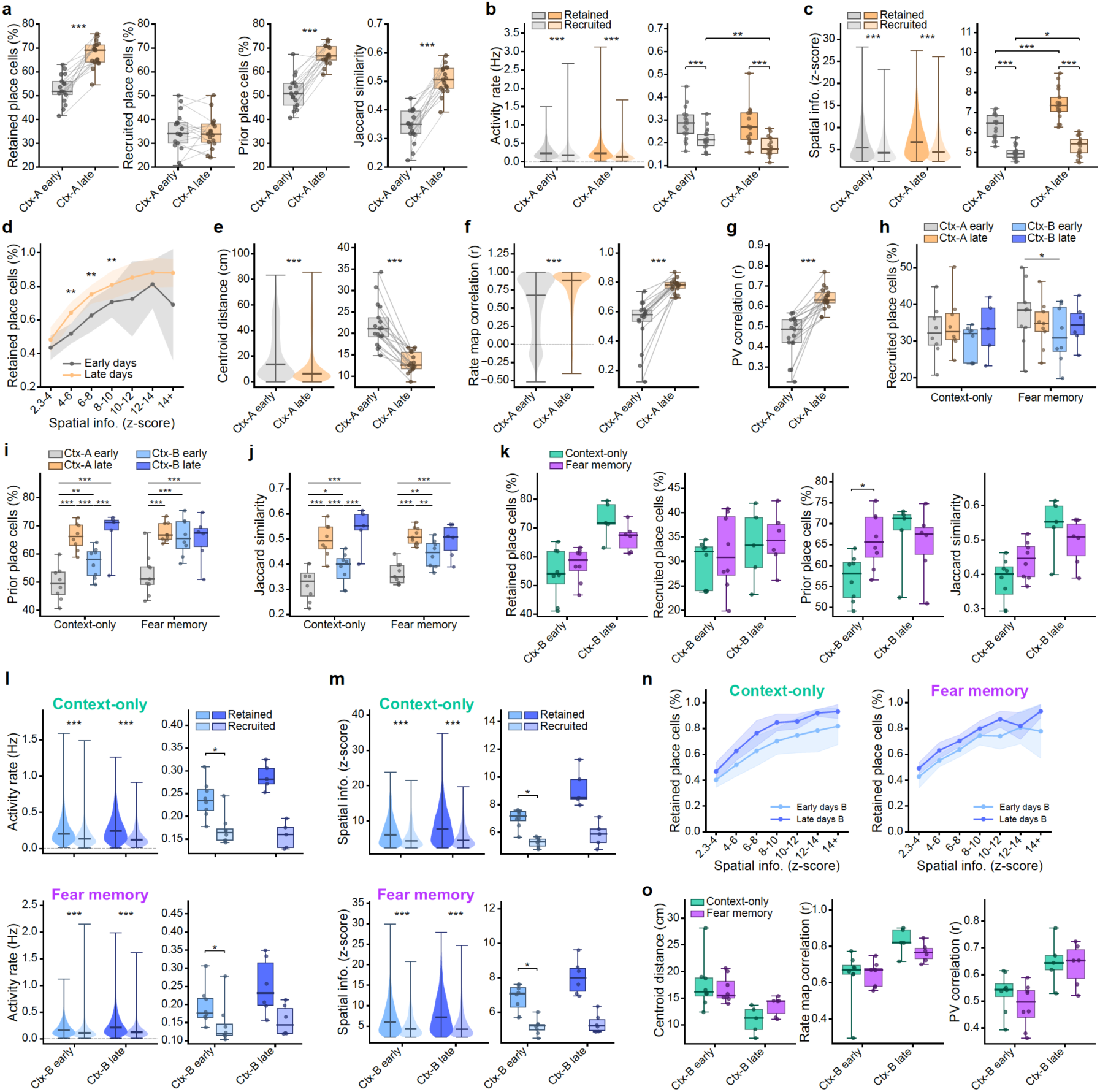
Across-day representational stability: place cell turnover, retention, and remapping across familiarization. (**a**) Place cell stability increased from early to late exposure in Ctx-A, with higher retention of place cells and greater population overlap (paired permutation tests, 10,000 two-tailed resamples. Diff is late minus early. Retained place cells, early 52.8% vs late 67.6%, diff = +14.75, *p* < 0.001; recruited place cells, 35.1% vs 34.6%, diff = −0.51, *p* = 0.77; prior place cells, 51.3% vs 67.0%, diff = +15.68, *p* < 0.001; Jaccard similarity, 0.34 vs 0.50, diff = +0.162, *p* < 0.001). Four metrics were computed for each consecutive day pair within each period and averaged: retained place cells, the percentage of place cells on day N that remain place cells on day N+1; recruited place cells, the percentage of non-place cells on day N that become place cells on day N+1; prior place cells, the percentage of place cells on day N+1 that were already place cells on day N; and Jaccard similarity, the intersection of the place cell populations on consecutive days divided by their union, giving a symmetric measure of overlap. Early is days 1–3 and late is days 7–9. Context-only and fear memory animals were combined into a single naive group, as all animals are experimentally naive at this stage; a linear mixed-effects model with a group × period interaction (animal as random intercept, REML) confirmed no group difference or group × period interaction for any metric (all *p ≥* 0.16), supporting this pooling. Boxes show the median and interquartile range, whiskers the full range; dots are individual animals and lines connect the same animal across periods. (**b**) Place cells retained from the previous day were more active than newly recruited place cells, in both the early and late periods, and this difference increased across exposure because recruited cell activity decreased while retained cell activity was unchanged (animal-level paired permutation tests, 10,000 two-tailed resamples, BH-FDR-corrected across the four comparisons. For the retained versus recruited comparisons, diff is retained minus recruited; for the early versus late comparisons, diff is late minus early. Retained vs recruited, early days, diff = +0.067, *p* < 0.001; retained vs recruited, late days, diff = +0.095, *p* < 0.001; retained cells early vs late, diff = −0.004, *p* = 0.80; recruited cells early vs late, diff = −0.032, *p* = 0.006). For each consecutive day pair within each period (as in **a**), place cells in the second session were classified as retained if they were also place cells in the first session, and as recruited if they were not. Left, cell-level distribution of activity rate for retained and recruited place cells, pooled across both consecutive pairs within each period; a cell active across all three sessions within a period contributes to both pairs. Cell-level linear mixed models (activity rate ∼ cell type, animal as random intercept, REML) were run as a sanity check and agreed with the animal-level tests (early, retained 0.276 vs recruited 0.216; late, 0.287 vs 0.184, both *p* < 0.001). Right, per-animal means, obtained by averaging across cells within each cell type and session pair and then across the two consecutive pairs within each period, giving one value per animal, cell type and period. Boxes show the median and interquartile range, whiskers the full range; dots are individual animals. (**c**) Place cells retained from the previous day carried more spatial information than newly recruited place cells, in both the early and late periods, and spatial information increased across exposure in both populations, more strongly in retained cells (animal-level paired permutation tests, 10,000 two-tailed resamples, BH-FDR-corrected across the four comparisons; conventions and layout as in **b**. Retained vs recruited, early days, diff = +1.29, *p* < 0.001; retained vs recruited, late days, diff = +2.10, *p* < 0.001; retained cells early vs late, diff = +1.13, *p* < 0.001; recruited cells early vs late, diff = +0.32, *p* = 0.011). Cell-level linear mixed models were run as a sanity check and agreed with the animal-level tests (early, retained 6.41 vs recruited 5.01; late, 7.57 vs 5.37, both *p* < 0.001). Left, cell-level distribution of spatial information z-score for retained and recruited place cells; right, per-animal means. Boxes show the median and interquartile range, whiskers the full range; dots are individual animals. (**d**) The likelihood that a place cell was retained the following day increased with its spatial information, and this relationship strengthened with experience (linear mixed-effects model, retained proportion ∼ period × z-score bin, animal as random intercept, REML; period × bin interaction *p* = 0.0026. Per-bin paired permutation tests comparing early with late, 10,000 two-tailed resamples, BH-FDR-corrected across the seven bins: retention increased from early to late in the 4–6, 6–8 and 8–10 bins, diff = +0.125, +0.126 and +0.101, all *p* ≤ 0.002; all other bins *p* ≤ 0.060). Place cells in the first session of each consecutive day pair were binned by spatial information z-score, and the fraction remaining place cells in the second session was computed per animal and bin, then averaged across the two pairs within each period. Lines show mean ± s.d. Context-only and fear memory animals were combined into a single naive group, as all animals are experimentally naive at this stage; a three-way model showed no significant group × period × bin interaction (*p* = 0.077). Significance stars reflect FDR-corrected per-bin *p*-values. (**e**) Place field centroids shifted less between consecutive days in the late than the early period (paired permutation test on per-animal means, 10,000 two-tailed resamples, diff = −8.68 cm, *p* < 0.001. Diff is late minus early). Only cells that remained place cells on consecutive days were included. Violins show the cell-level distribution and boxes the per-animal means; a cell-level linear mixed model (centroid displacement ∼ period, animal as random intercept, REML) agreed (early 21.30 vs late 12.98 cm, *p* < 0.001). Values were averaged across the two consecutive day pairs within each period. Context-only and fear memory animals were combined into a single naive group, as all animals are experimentally naive at this stage; a model including a group × period interaction showed none (*p* = 0.71). Dots are individual animals and lines connect the same animal across periods. (**f**) Rate map correlation between consecutive days was higher in the late than the early period (paired permutation test on per-animal means, 10,000 two-tailed resamples, diff = +0.250, *p* < 0.001. Diff is late minus early). Rate map correlation is the Pearson r between consecutive days’ activity rate maps, computed over spatial bins visited in both sessions. Violins show the cell-level distribution and boxes the per-animal means; a cell-level linear mixed model agreed (early 0.536 vs late 0.782, *p* < 0.001). Inclusion, average and group pooling as in **e**; the group × period interaction was not significant (*p* = 0.23). (**g**) Population vector (PV) correlation between consecutive days was higher in the late than the early period (paired permutation test on per-animal means, 10,000 two-tailed resamples, diff = +0.185, *p* < 0.001. Diff is late minus early). PV correlation was computed per spatial bin across all retained place cells per animal and averaged across bins visited in both sessions, giving one value per animal per period. Inclusion, averaging and group pooling as in **e**; the group × period interaction was not significant (*p* = 0.36). (**h**) Place cell recruitment did not change from early to late exposure in either context or group (statistical approach, period definitions and sign convention as in **Figure 4e**. Context-only: no contrast reached significance, all *p* ≥ 0.28. Fear memory: recruitment was lower in early Ctx-B than in early Ctx-A, Δ = +6.28, *p* = 0.013; no other contrast reached significance, all *p* ≥ 0.23). Each dot is one animal; boxes show the group median and interquartile range, whiskers the full range. (**i**) The proportion of prior place cells increased from early to late exposure in both contexts in context-only animals, but in fear memory animals this increase was absent in Ctx-B (statistical approach, period definitions and sign convention as in **Figure 4e**. Context-only: Ctx-A early → late, Δ = −16.59, *p* < 0.001; Ctx-B early → late, Δ = −10.28, *p* = 0.0010; Ctx-A late → Ctx-B early, Δ = +9.24, *p* = 0.0006; Ctx-A early → Ctx-B early, Δ = −7.35, *p* = 0.005; Ctx-A early → Ctx-B late, Δ = −17.64, *p* < 0.001. Fear memory: Ctx-A early → late, Δ = −14.87, *p* < 0.001; Ctx-A early → Ctx-B early, Δ = −12.87, *p* < 0.001; Ctx-A early → Ctx-B late, Δ = −16.28, *p* < 0.001; Ctx-B early → late, Δ = +3.73, *p* = 0.82, not significant). The absence of an increase across Ctx-B in fear memory animals contrasts with the corresponding increase in Ctx-A of the same animals (Δ = −14.87) and with Ctx-B in context-only animals (Δ = −10.28). All remaining contrasts had *p* ≥ 0.50. Each dot is one animal; boxes show the group median and interquartile range, whiskers the full range. (**j**) Jaccard similarity of the place cell population across consecutive days increased from early to late exposure in both contexts in context-only animals, but in fear memory animals this increase was absent in Ctx-B (statistical approach, period definitions and sign convention as in **Figure 4e**. Context-only: Ctx-A early → late, Δ = −0.175, *p* < 0.001; Ctx-B early → late, Δ = −0.151, *p* < 0.001; Ctx-A late → Ctx-B early, Δ = +0.114, *p* < 0.001; Ctx-A early → Ctx-B early, Δ = −0.061, *p* = 0.029; Ctx-A early → Ctx-B late, Δ = −0.221, *p* < 0.001. Fear memory: Ctx-A early → late, Δ = −0.150, *p* < 0.001; Ctx-A early → Ctx-B early, Δ = −0.076, *p* = 0.002; Ctx-A late → Ctx-B early, Δ = +0.067, *p* = 0.005; Ctx-A early → Ctx-B late, Δ = −0.156, *p* < 0.001; Ctx-B early → late, Δ = −0.022, *p* = 0.065, not significant). The absence of an increase across Ctx-B in fear memory animals contrasts with the corresponding increase in Ctx-A of the same animals (Δ = −0.150) and with Ctx-B in context-only animals (Δ = −0.151). All remaining contrasts had p ≥ 0.22. Each dot is one animal; boxes show the group median and interquartile range, whiskers the full range. (**k**) Place cell stability across consecutive days in Ctx-B did not differ between context-only and fear memory animals, except that early in Ctx-B, place cells in fear memory animals were more likely to have already been place cells on the preceding day (independent permutation tests, 10,000 two-tailed resamples, BH-FDR-corrected across the two Ctx-B periods within each metric. Diff is fear memory minus context-only. Prior place cells: early Ctx-B, diff = +9.10, *p* = 0.027; late Ctx-B, diff = −2.17, *p* = 0.65. Retained place cells: early, diff = +3.05, *p* = 0.45; late, diff = −6.03, *p* = 0.25. Recruited place cells: early, diff = +2.24, *p* = 0.77; late, diff = +1.17, *p* = 0.77. Jaccard similarity: early, diff = +0.058, *p* = 0.15; late, diff = −0.047, *p* = 0.33). The four metrics are show in separate plots, each comparing the two groups in the early and late Ctx-B periods; metric definitions, period definitions and averaging as in **a**. Each dot is one animal; boxes show the group median and interquartile range, whiskers the full range. (**l**) In Ctx-B, place cells retained from the previous day were more active than newly recruited place cells in both groups, and as in Ctx-A this difference was larger in the late period, though only the early-period comparison survived correction (animal-level paired permutation tests, 10,000 two-tailed resamples, BH-FDR-corrected across the four comparisons within each group; conventions and layout as in **b**. Context-only: retained vs recruited, early diff = +0.067, *p* = 0.031; late diff = +0.129, *p* = 0.083; retained cells early vs late, diff = +0.054, *p* = 0.083; recruited cells early vs late, diff = −0.003, *p* = 0.63. Fear memory: retained vs recruited, early diff = +0.050, *p* = 0.031; late diff = +0.094, *p* = 0.063; retained cells early vs late, diff = +0.041, *p* = 0.083; recruited cells early vs late, diff = +0.001, *p* = 1.00). The per-group sample sizes limit the resolution of the paired permutation tests in the late period. Cell-level linear mixed models agreed in direction and were significant throughout (Context-only: early, retained 0.243 vs recruited 0.175; late, 0.287 vs 0.163. Fear memory: early, 0.198 vs 0.146; late, 0.264 vs 0.153, all *p* < 0.001). Context-only is shown on top and fear memory below; left, cell-level distributions; right, per-animal means. (**m**) In Ctx-B, place cells retained from the previous day carried more spatial information than newly recruited place cells in both groups, and as in Ctx-A this difference was larger in the late period, though only the early-period comparison survived correction (animal-level paired permutation tests, 10,000 two-tailed resamples, BH-FDR-corrected across the four comparisons within each group; conventions and layout as in **b**. Context-only: retained vs recruited, early diff = +1.74, *p* = 0.031; late diff = +3.34, *p* = 0.083; retained cells early vs late, diff = +2.07, *p* = 0.083; recruited cells early vs late, diff = +0.55, *p* = 0.25. Fear memory: retained vs recruited, early diff = +1.66, *p* = 0.031; late diff = +2.68, *p* = 0.063; retained cells early vs late, diff = +1.36, *p* = 0.17; recruited cells early vs late, diff = +0.27, *p* = 0.25). As in **l**, the per-group sample sizes limit the resolution of the paired permutations tests in the late period. Cell-level linear mixed models agreed in direction and were significant throughout (Context-only: early, retained 7.03 vs recruited 5.29; late, 8.83 vs 5.59. Fear memory: early, 6.93 vs 5.18; late, 8.18 vs 5.30; all *p* < 0.001). Context-only is shown on top and fear memory below; left, cell-level distributions; right, per-animal means. (**n**) In Ctx-B, the likelihood that a place cell was retained the following day increased with its spatial information in both groups, but unlike in Ctx-A this relationship did not strengthen detectably from early to late exposure (linear mixed-effects models, retained proportion ∼ period × z-score bin, animal as random intercept, REML, run separately per group. Main effect of z-score bin: context-only *p* = 0.013, fear memory *p* = 0.013. Period × bin interaction: context-only *p* = 0.30, fear memory *p* = 0.24). Retention increased from early to late in nearly every bin in both groups, but only five animals contribute to each bin, so per-bin comparisons were not tested. Binning, averaging and plotting as in **d**. Context-only is shown left and fear memory right; lines show mean ± s.d. (**o**) Place field remapping across consecutive days in Ctx-B did not differ between context-only and fear memory animals in either period, for any of the three remapping measures (independent permutation test, 10,000 two-tailed resamples, BH-FDR-corrected across the two Ctx-B periods within each metric. Diff is fear memory minus context-only. Centroid displacement: early, diff = −0.96 cm, *p* = 0.68; late, diff = +2.64 cm, *p* = 0.17. Rate map correlation: early, diff = +0.007, *p* = 0.96; late, diff = −0.064, *p* = 0.26. PV correlation: early, diff = −0.047, *p* = 0.46; late, diff = −0.011, *p* = 0.84). The three metrics are shown in separately plots, each comparing the two groups in the early and late Ctx-B periods; metric definitions and averaging as in **e–g**. Each dot is one animal; boxes show the group median and interquartile range, whiskers the full range.

**Figure S10:**
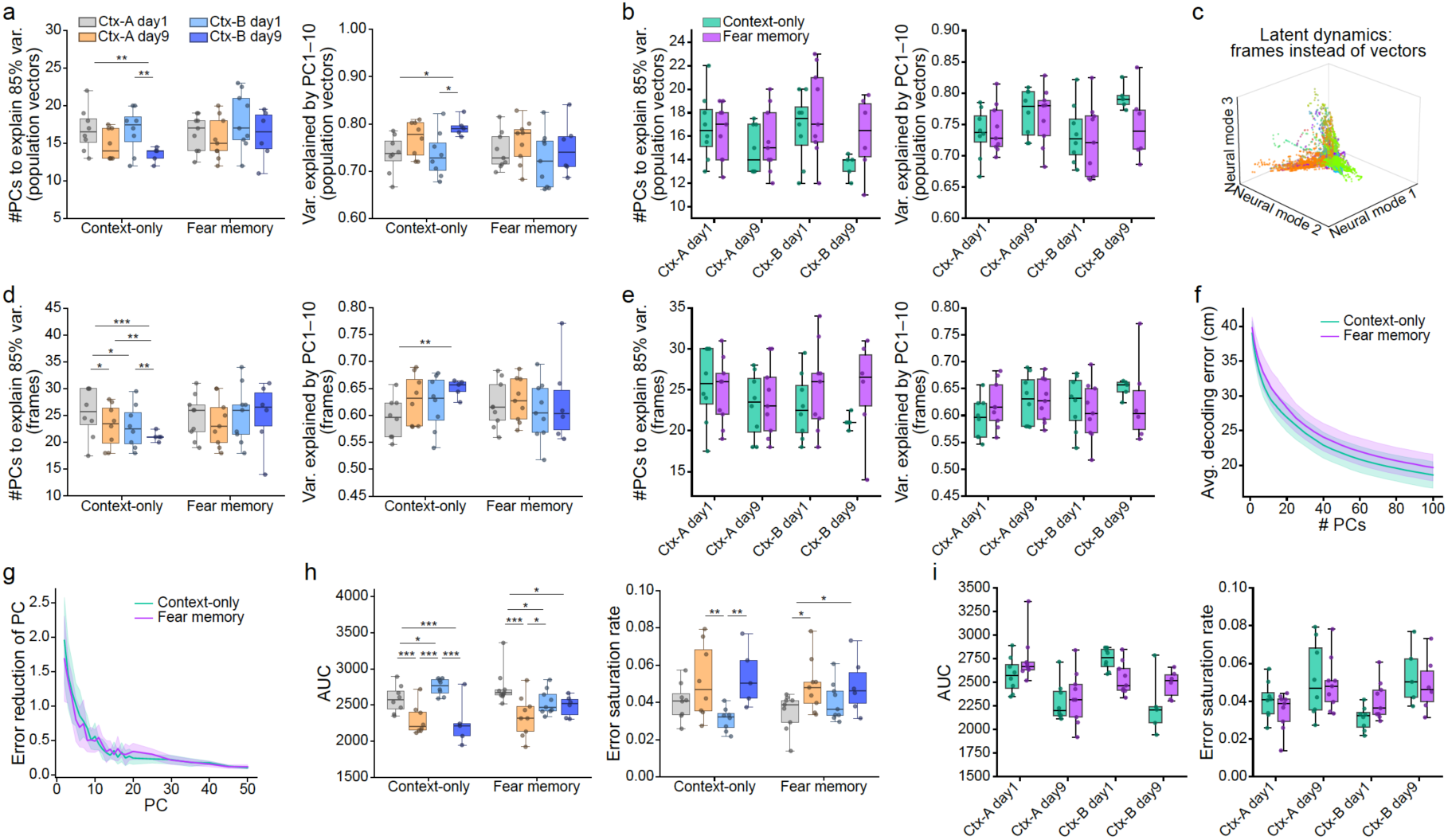
PCA dimensionality of spatially binned and per-frame population activity, and decoding performance as a function of dimensionality. (**a**) The dimensionality of spatial population activity decreased across familiarization with Ctx-B in context-only animals, but this decrease was absent in Ctx-A and in fear memory animals (linear mixed-effects model with session as a categorical fixed effect and animal as random intercept, REML; pairwise contrasts BH-FDR-corrected across all comparisons. Δ is the earlier minus the later session, in number of PCs for the left panel and in cumulative variance explained for the right panel; lower dimensionality on the later session gives a positive Δ on the left and a negative Δ on the right. Number of PCs to reach 85% variance, Context-only: Ctx-B day1 → day9, Δ = +4.10, *p* = 0.009; Ctx-A day1 → Ctx-B day9, Δ = +3.80, *p* = 0.009; Ctx-A day1 → day9, Δ = +2.00, *p* = 0.098, not significant; Ctx-A day9 → Ctx-B day1, Δ = −1.94, *p* = 0.098, not significant. Fear memory: all contrasts *p* ≥ 0.064. Variance explained by the first 10 PCs, Context-only: Ctx-B day1 → day9, Δ = −0.070, *p* = 0.025; Ctx-A day1 → Ctx-B day9, Δ = −0.059, *p* = 0.025; all remaining contrasts *p* ≥ 0.14. Fear memory: all contrasts *p* ≥ 0.16). The absence of a decrease across Ctx-B in fear memory animals contrasts with the corresponding decrease in context-only animals, and both metrics give the same result: fewer PCs were needed to reach 85% of the variance and more variance was captured by the first 10PCs on the later session. Left, number of PCs to explain 85% of the variance. Right, cumulative variance explained by the first 10 PCs. The model was chosen to handle the repeated-measures structure and missing data at later timepoints (attrition due to logistical constraints; missing-at-random assumed); sensitivity analyses (bootstrap, permutation, paired t-tests, Wilcoxon signed-rank) on complete-case pairs were consistent for the well-powered comparisons. Each dot is one animal; boxes show the group median and interquartile range, whiskers the full range. (**b**) The dimensionality of spatial population activity did not differ between context-only and fear memory animals at any session (independent permutation tests, 10,000 two-tailed resamples, BH-FDR-corrected across the four sessions. Number of PCs to reach 85% variance: all *p* ≥ 0.61. Variance explained by the first 10 PCs: all *p* ≥ 0.48). Left, number of PCs required to explain 85% of the variance. Right, cumulative variance explained by the first 10 PCs. Sessions are Ctx-A day1, Ctx-A day9, Ctx-B day1 and Ctx-B day9. Each dot is one animal; boxes show the group median and interquartile range, whiskers the full range. (**c**) The first three principal components of per-frame neuronal activity in an example session. PCA was applied directly to per-frame activity rather than to spatially binned population vectors as elsewhere, capturing the moment-to-moment population state. For visualization, 6,000 randomly selected frames during mobility are shown, corresponding to five minutes of recording time; the corresponding analyses used all frames during mobility. (**d**) The dimensionality of per-frame population activity decreased across sessions in context-only animals but did not change in fear memory animals (statistical approach as in **a**. Δ is the earlier minus the later session, in number of PCs for the left panel and in cumulative variance explained for the right panel; lower dimensionality on the later session gives a positive Δ on the left and a negative Δ on the right. Number of PCs to reach 85% variance, Context-only: Ctx-A day1 → day9, Δ = +2.38, *p* = 0.024; Ctx-A day1 → Ctx-B day1, Δ = +2.56, *p* = 0.018; Ctx-A day1 → Ctx-B day9, Δ = +6.60, *p* < 0.001; Ctx-A day9 → Ctx-B day9, Δ = +4.60, *p* = 0.003; Ctx-B day1 → day9, Δ = +3.60, *p* = 0.004; Ctx-A day9 → Ctx-B day1, Δ = +0.19, *p* = 0.85, not significant. Fear memory: all contrasts *p* ≥ 0.62. Variance explained by the first 10 PCs, Context-only: Ctx-A day1 → Ctx-B day9, Δ = −0.072, *p* = 0.001; all remaining contrasts *p* ≥ 0.064. Fear memory: all contrasts *p* ≥ 0.32). Both metrics give the same result: in context-only animals fewer PCs were needed to reach 85% of the variance and more variance was captured by the first 10 PCs on later sessions, whereas in fear memory animals dimensionality was unchanged throughout. Left, number of PCs required to explain 85% of the variance. Right, cumulative variance explained by the first 10 PCs. PCA was applied to per-frame activity during mobility rather than to spatially binned population vectors, as in **c**. Each dot is one animal; boxes show the group median and interquartile range, whiskers the full range. (**e**) The dimensionality of per-frame population activity did not differ between context-only and fear memory animals at any session (independent permutation tests, 10,000 two-tailed resamples, BH-FDR-corrected across the four sessions. The number of PCs to reach 85% variance: all *p* ≥ 0.69. Variance explained by the first 10 PCs: all *p* ≥ 0.76). Left, number of PCs required to explain 85% of the variance. Right, cumulative variance explained by the first 10 PCs. Sessions are Ctx-A day1, Ctx-A day9, Ctx-B day1 and Ctx-B day9. PCA was applied to per-frame activity during mobility rather than to spatially binned population vectors, as in **c**. Each dot is one animal; boxes show the group median and interquartile range, whiskers the full range. (**f**) Decoding error decreased with the number of principal components (PCs) included and did not differ between context-only and fear memory animals at any dimensionality (independent permutation tests, 10,000 two-tailed resamples, BH-FDR-corrected across the 26 tested dimensions, all *p* ≥ 0.28). Lines show the group mean decoding error as a function of the number of PCs included, and shading the ± s.d. across animals. PCA was applied to spatially binned population vectors, as in **a**. Neural population activity was mean-centered and projected onto the first d principal components, with d swept across a dense range (1– 20) and a sparse range (25–100, in steps of 5). At each dimensionality, position was decoded by linear regression with an 80/20 train/test split, and decoding error was quantified as the median Euclidean distance (cm) between predicted and actual position. Decoding was repeated across all sessions and subsampling iterations, and per-animal decoding error was summarized as the median across sessions and iterations. Group comparisons were performed at dimensionalities up to 50 PCs. (**g**) The marginal reduction in decoding error per additional principal component did not differ between context-only and fear memory animals at any dimensionality (statistical approach as in **f**, BH-FDR-corrected across the 25 tested dimensions, all *p* ≥ 0.50). Marginal error reduction (cm per PC) was computed per animal as the finite difference of the decoding error curve in **f** and is shown for PCs 1–50. Lines show the group mean and shading the ± s.d. across animals. (**h**) Decoding performance across the full range of PCA dimensionalities improved across familiarization with Ctx-B in context-only animals, but this improvement was absent in fear memory animals (statistical approach as in **a**. Δ is the earlier minus the later session, in AUC units for the left panel and in decay constant for the right panel; better decoding on the later session gives a positive Δ on the left and a negative Δ on the right. AUC, Context-only: Ctx-A day1 → day9, Δ = +285.7, *p* < 0.001; Ctx-A day1 → Ctx-B day1, Δ = −167.6, *p* = 0.012; Ctx-A day1 → Ctx-B day9, Δ = +286.4, *p* < 0.001; Ctx-A day9 → Ctx-B day1, Δ = −453.3, *p* < 0.001; Ctx-B day1 → day9, Δ = +472.4, *p* < 0.001; Ctx-A day9 → Ctx-B day9, Δ = +26.3, *p* = 0.79, not significant. Fear memory: Ctx-A day1 → day9, Δ = +403.0, *p* < 0.001; Ctx-A day1 → Ctx-B day1, Δ = +207.2, *p* = 0.038; Ctx-A day1 → Ctx-B day9, Δ = +333.5, *p* = 0.012; Ctx-A day9 → Ctx-B day1, Δ = −195.8, *p* = 0.040; Ctx-B day1 → day9, Δ = +67.2, *p* = 0.40, not significant; Ctx-A day9 → Ctx-B day9, Δ = −56.7, *p* = 0.34, not significant. Saturation rate, Context-only: Ctx-A day9 → Ctx-B day1, Δ = +0.020, *p* = 0.001; Ctx-B day1 → day9, Δ = −0.020, *p* = 0.008; all remaining contrasts *p* ≥ 0.092. Fear memory: Ctx-A day1 → day9, Δ = −0.014, *p* = 0.029; Ctx-A day1 → Ctx-B day9, Δ = −0.017, *p* = 0.048; Ctx-B day1 → day9, Δ = −0.005, *p* = 0.23, not significant; all remaining contrasts *p* ≥ 0.17). The absence of an improvement across Ctx-B in fear memory animals contrasts with the corresponding improvement in context-only animals on both metrics (AUC, Δ = +472.4; saturation rate, Δ = −0.020), mirroring the decoding and place-cell results in **Figure 4**. Left, area under the decoding error curve, computed as the numerical integral of decoding error over the tested dimensionality range (1–100 PCs) by the trapezoidal rule; lower values indicate better decoding across all dimensionalities. Right, saturation rate, the decay constant b of an exponential model (error = a·e^(−b·d) + c) fitted to each animal’s decoding curve; higher values indicate that performance saturates at lower dimensionality. Each dot is one animal; boxes show the group median and interquartile range, whiskers the full range. (**i**) Decoding performance across the full range of PCA dimensionalities did not differ significantly between context-only and fear memory animals at any session (independent permutation tests, 10,000 two-tailed resamples, BH-FDR-corrected across the four sessions. AUC: Ctx-B day1, *p* = 0.054; all remaining sessions *p* ≥ 0.20. Saturation rate: all *p* ≥ 0.12). Left, area under the decoding error curve (AUC), computed as the numerical integral of decoding error over the tested dimensionality range (1–100 PCs) by the trapezoidal rule. Right, saturation rate, the decay constant b of an exponential model (error = a·e^(−b·d) + c) fitted to each animal’s decoding curve. Sessions are Ctx-A day1, Ctx-A day9, Ctx-B day1 and Ctx-B day9. Each dot is one animal; boxes show the group median and interquartile range, whiskers the full range.

**Figure S11:**
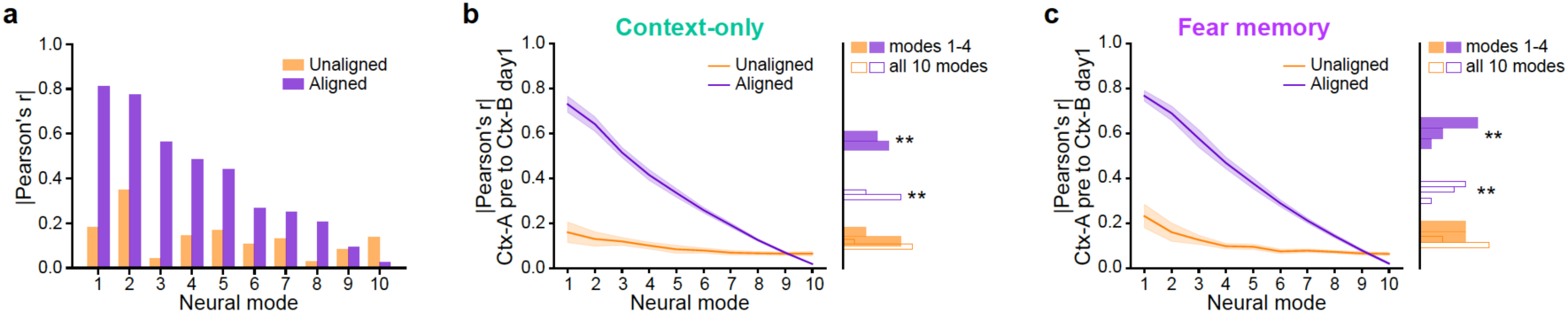
Cross-session correspondence of latent dynamics before and after CCA alignment. (**a**) Correlation between corresponding neural modes of Ctx-A pre and Ctx-B day1 before and after CCA alignment, for the example session pair in **Figure 5a**. Bars show the absolute Pearson correlation (|r|) for each of the 10 modes, unaligned (orange) and aligned (purple). Unaligned correlations are computed between corresponding principal components of the two sessions, which need not share a coordinate frame. Aligned correlations are the canonical variates: Pearson correlations between the session axes optimally rotated by CCA to maximize correspondence, giving the maximal achievable cross-session correspondence. Alignment increased the correlation across all 10 modes, with the largest gains in the leading modes. (**b**) In context-only animals, CCA alignment increased the correlation between Ctx-A pre and Ctx-B day1 latent dynamics for both the leading 4 modes and all 10 modes (paired permutation tests, 10,000 two-tailed resamples, BH-FDR-corrected across the two comparisons, n = 8. Δ is aligned minus unaligned. Leading 4 modes: unaligned 0.127 ± 0.020, aligned 0.575 ± 0.024, Δ = +0.448, *p* = 0.007. All 10 modes: unaligned 0.093 ± 0.008, aligned 0.330 ± 0.012, Δ = +0.236, *p* = 0.007). Left, absolute Pearson correlations between corresponding principal components (unaligned, orange) and canonical variates following CCA (aligned, purple) across the 10 neural modes, defined as in **a**; lines show the group mean and shading the 95% CI across animals (t-based). Right, distribution across animals of the mean correlation for the leading 4 modes (filled) and all 10 modes (unfilled); values are mean ± s.d. across animals. Each value is the median across 50 random-subsampling iterations of 100 neurons each, aggregated as in **Figure 5c**. (**c**) In fear memory animals, CCA alignment likewise increased the correlation for both the leading 4 modes and all 10 modes (paired permutation tests, 10,000 two-tailed resamples, BH-FDR-corrected across the two comparisons, n = 9. Δ is aligned minus unaligned. Leading 4 modes: unaligned 0.153 ± 0.024, aligned 0.625 ± 0.034, Δ = +0.471, *p* = 0.003. All 10 modes: unaligned 0.106 ± 0.011, aligned 0.362 ± 0.020, Δ = +0.256, *p* = 0.003). Panels as in **b**, for fear memory animals.

**Figure S12:**
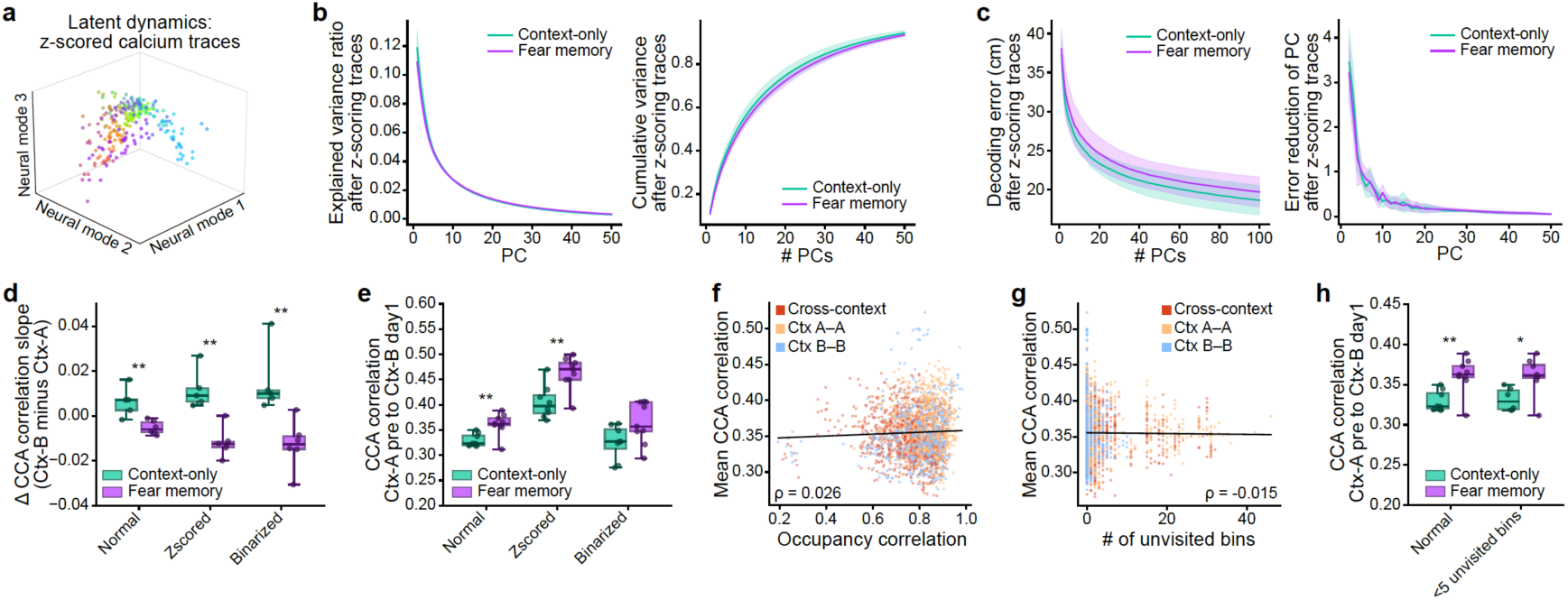
Neural activity and behavioral controls for the CCA analyses. (**a**) The first three neural modes of spatial population vectors in an example recording session, computed after z-scoring neuronal activity. Activity was z-scored across time within the session before construction of the spatial population vectors, controlling for the possibility that differences in activity scale across neurons drive the principal components and alignment. (**b**) The dimensionality of spatial population activity did not differ between context-only and fear memory animals after z-scoring neuronal activity. Neither the explained variance of individual principal components nor the cumulative variance differed between groups at any component (independent permutation tests, 10,000 two-tailed resamples, BH-FDR-corrected across the 50 components, all *p* ≥ 0.13 for both metrics), and the number of PCs required to reach 85% cumulative variance was also comparable (independent permutation test, 10,000 two-tailed resamples. Diff is fear memory minus context-only. Diff = +2.44 PCs, d = +0.89, *p* = 0.11; context-only: median 30.5, mean 31.00 ± 3.42; fear memory: median 33, mean 33.44 ± 2.01). Left, mean explained variance ratio per principal component. Right, mean cumulative variance ratio as a function of the number of PCs included. Shading shows ± s.d. across animals. Activity was z-scored as in **a**; subsampling and aggregation as in **figure 5b**. (**c**) Decoding error decreased with the number of principal components included and did not differ between context-only and fear memory animals at any dimensionality after z-scoring neuronal activity, either in decoding error itself or in the marginal reduction per additional component (independent permutation tests, 10,000 two-tailed resamples, BH-FDR-corrected across the tested dimensionalities. Decoding error: 26 dimensions tested, all *p* ≥ 0.26. Marginal error reduction: 25 dimensions tested, *p* ≥ 0.56). Left, group mean decoding error as a function of the number of neural modes included. Right, marginal reduction in decoding error per additional neural mode (cm per mode), computed per animal as the finite difference of the decoding error curve and shown for the first 50 modes. Lines show the group mean and shading the ± s.d. across animals. Population activity was z-scored as in **a** and projected onto the first d principal components (d = 1–20 in steps of 1, and 25–100 in steps of 5) before training a linear regression decoder; decoding error was quantified as the median Euclidean distance (cm) between predicted and true position. Group comparisons were performed at dimensionalities up to 50 PCs. (**d**) The change in CCA alignment slope from Ctx-A to Ctx-B differed between groups under all three analysis approaches, normal activity, z-scored activity, and binarized traces, with context-only animals increasing alignment more steeply in Ctx-B and fear memory animals showing the opposite (independent permutation tests, 10,000 two-tailed resamples, BH-FDR-corrected across the three approaches. Diff is fear memory minus context-only. Normal: context-only 0.0063 ± 0.0067, fear memory −0.0051 ± 0.0030, diff = −0.0114, *p* = 0.009. Z-scored: context-only 0.0119 ± 0.0088, fear memory −0.0117 ± 0.0065, diff = −0.0236, *p* = 0.007. Binarized: context-only 0.0151 ± 0.0148, fear memory −0.0128 ± 0.0108, diff = −0.0279, *p* = 0.007). Within each group, Δ slope did not differ significantly between approaches (paired permutation tests, 10,000 two-tailed resamples, BH-FDR-corrected across the three approach pairs within each group, all *p* ≥ 0.19); these comparisons are not shown on the plot. With five and six animals per group, the smallest attainable two-tailed *p* values are 0.063 and 0.031, so these paired comparisons are not powered to detect a difference between approaches after correction. Δ slope was compute per animal as the alignment slope in Ctx-B minus that in Ctx-A, estimated from CCA correlations across same-context session pairs as in **figure 5e**. Z-scoring is as in **a**. Each dot is one animal (context-only n = 5, fear memory n = 6); boxes show the group median and interquartile range, whiskers the full range. Values are mean ± s.d. across animals. (**e**) The canonical correlation between Ctx-A pre and Ctx-B day1 was higher in fear memory than context-only animals under the normal and z-scored approaches, and showed the same direction under binarization without reaching significance (independent permutation tests, 10,000 two-tailed resamples, BH-FDR-corrected across the three approaches. Diff is fear memory minus context-only. Normal: context-only 0.3295 ± 0.0127, fear memory 0.3617 ± 0.0216, diff = +0.0322, *p* = 0.006. Z-scored: context-only 0.4051 ± 0.0331, fear memory 0.4639 ± 0.0316, diff = +0.0588, *p* = 0.006. Binarized: context-only 0.3256 ± 0.0335, fear memory 0.3645 ± 0.0412, diff = +0.0389, *p* = 0.051). Within each group, z-scoring raised the correlation relative to both the normal and binarized approaches, which did not differ from each other (paired permutation tests, 10,000 two-tailed resamples, BH-FDR-corrected across the three approach pairs within each group. Context-only: normal versus z-scored, Δ = −0.0756, *p* = 0.012; z-scored versus binarized, Δ = +0.0795, *p* = 0.012; normal versus binarized, Δ = +0.0039, *p* = 0.73. Fear memory: normal versus z-scored, Δ = −0.1021, *p* = 0.006; z-scored versus binarized, Δ = +0.0993, *p* = 0.006; normal versus binarized, Δ = −0.0028, *p* = 0.78); these comparisons are not shown on the plot. Data presentation and analysis as in **d**, with n = 8 context-only and n = 9 fear memory animals. (**f**) Occupancy similarity showed a weak positive relationship with CCA alignment strength within animals, confined to Ctx-A → Ctx-A session pairs (sign-flip permutation test on per-animal correlations, 10,000 two-tailed resamples, n = 17 animals. Overall: mean within-animal Spearman ρ = 0.129, *p* = 0.036. Per session-pair type, BH-FDR-corrected across the three types: Ctx-A → Ctx-A: mean ρ = +0.217, *p* = 0.003. Cross-context: mean ρ = −0.100, *p* = 0.21. Ctx-B → Ctx-B: mean ρ = −0.079, *p* = 0.53). Each point is one session pair from one animal (n = 2,314 pairs), colored by session-pair type (Ctx-A → Ctx-B, Ctx-A → Ctx-A, Ctx-B → Ctx-B). The x-axis shows the occupancy correlation (Pearson r) between the two sessions, computed over bins visited in both, and the y-axis the mean canonical correlation across the first 10 modes aggregated as in **figure 5c**. A linear fit is shown for visualization. The Spearman ρ displayed in the panel is based on pooled session-pair data (ρ = 0.026), whereas statistical inference was performed on the per-animal correlations. (**g**) The number of unvisited spatial bins showed a weak negative relationship with CCA alignment strength within animals, present for both cross-context and Ctx-A → Ctx-A session pairs (sign-flip permutation test on per-animal correlations, 10,000 two-tailed resamples. Overall: mean within-animal Spearman ρ = −0.223, *p* = 0.002, n = 15 animals. Per session-pair type, BH-FDR-corrected across the three types: Cross-context: mean ρ = −0.199, *p* = 0.002, n = 15. Ctx-A → Ctx-A: mean ρ = −0.373, *p* = 0.002, n = 13. Ctx-B → Ctx-B: mean ρ = +0.105, *p* = 0.46, n = 11. Animals were excluded from a comparison where coverage was complete across the relevant sessions, leaving no variance to correlate). Each point is one session pair from one animal, colored by session-pair type as in **f**. The x-axis shows the number of bins left unvisited across the two sessions of each pair, and the y-axis the mean canonical correlation across the first 10 modes. A linear fit is shown for visualization. The Spearman ρ displayed in the panel is based on pooled session-pair data (ρ = −0.015), whereas statistical inference was performed on the per-animal correlations. (**h**) The group difference in canonical correlation between Ctx-A pre and Ctx-B day1 was preserved when session pairs with ≥ 5 unvisited spatial bins were excluded (independent permutation tests, 10,000 two-tailed resamples, BH-FDR-corrected across the two approaches. Diff is fear memory minus context-only. Normal: context-only 0.3295 ± 0.0127, n = 8; fear memory 0.3617 ± 0.0216, n = 9; diff = +0.0322, *p* = 0.008. Filtered: context-only 0.3317 ± 0.0142, n = 6; fear memory 0.3613 ± 0.0231, n = 8; diff = +0.0296, *p* = 0.019). The filter retained 1,962 of 2,314 session pairs across all animals, and removed the Ctx-A pre → Ctx-B day1 pair entirely for two context-only and one fear memory animal; values for all retained animals were unchanged. Each dot is one animal; boxes show the group median and interquartile range, whiskers the full range. Values are mean ± s.d. across animals.

**Figure S13:**
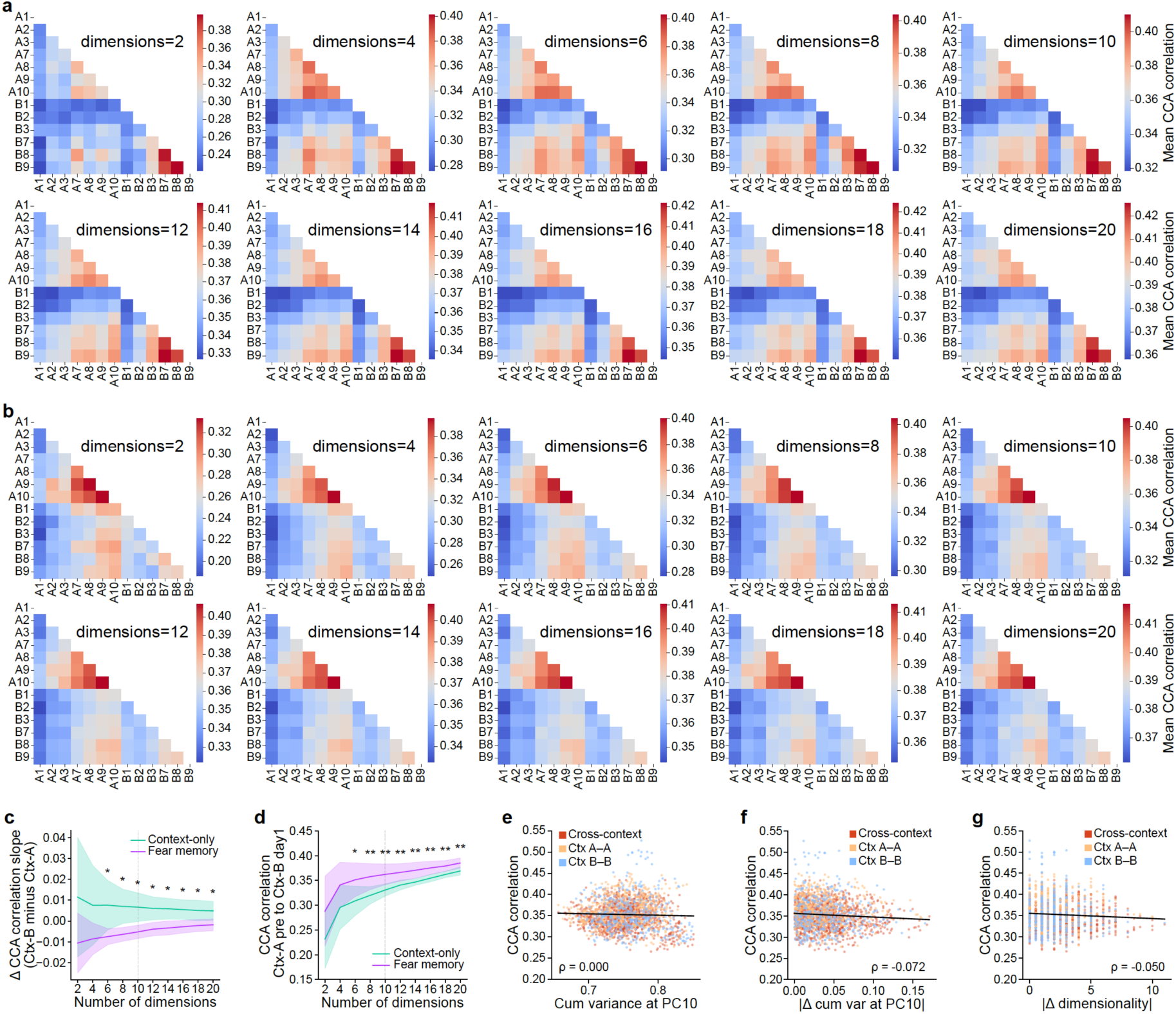
Effect of CCA input dimensionality and relationship between session dimensionality and alignment. (**a**) Similarity between latent dynamics across session pairs in context-only animals (n = 8), with CCA performed on a range of input dimensionalities (dimensions, panel titles), sweeping from 2 to 20. PCA was performed as in the main analyses, and the leading d principal components (panel titles) of each session were then passed to CCA, so that each panel is a separate CCA fitted on a different number of components rather than a truncation of a single higher-dimensional solution. Each heatmap shows the same comparison as in **Figure 5d**. Color indicates the mean canonical correlation between the two sessions of each pair, with warmer colors indicating greater similarity; each heatmap is scaled to its own range, so that the relative structure across session pairs, rather than the absolute correlation, is comparable across dimensionalities. Only the early and late sessions of each context are shown (Ctx-A: A1–A3 and A7–A10; Ctx-B: B1–B3 and B7–B9), and the grey lines separate the two contexts. Only the lower triangle is shown; the diagonal is omitted, as within-session comparisons were not included. At each d, the canonical correlations were taken as the median across 50 subsampling iterations per mode and then averaged across the d modes, giving one similarity value per session pair per animal, and heatmaps show the mean of these values across animals. (**b**) The same, for fear memory animals (n = 9). Panels as in **a**. (**c**) The difference in CCA alignment slope between Ctx-B and Ctx-A separated the groups across the tested dimensionality range, with context-only animals increasing alignment more steeply in Ctx-B and fear memory animals showing the opposite at every dimensionality (independent permutation tests, 10,000 two-tailed resamples, BH-FDR-corrected across the 10 tested dimensionalities. Diff is fear memory minus context-only. Significant from 6 dimensions onwards, all *p* ≤ 0.027; not significant at 2 and 4 dimensions, both *p* ≥ 0.086. At the 10 dimensions used in the main analyses: context-only +0.0063 ± 0.0067, fear memory −0.0051 ± 0.0030, diff = −0.0114, *p* = 0.019). Lines show the group mean and shading the ± s.d. across animals (context-only n = 5, fear memory n = 6). The dashed vertical line marks 10 dimensions, the dimensionality used in the main analyses. The dimensionality sweep and aggregation are as in **a**; per-animal Δ slopes were computed as the alignment slope across days in Ctx-B minus that in Ctx-A, using the same sessions and procedure as in **Figure 5e**. (**d**) The group difference in canonical correlation between Ctx-A pre and Ctx-B day1 was robust to the choice of input dimensionality, with fear memory animals showing higher correlations than context-only animals at every dimensionality tested (independent permutation tests, 10,000 two-tailed resamples, BH-FDR-corrected across the 10 tested dimensionalities. Diff is fear memory minus context-only. Significant from 4 dimensions onwards, all *p* ≤ 0.048, with Cohen’s d ranging from 1.10 to 2.00 over that range; not significant at 2 dimensions, *p* = 0.074. At the 10 dimensions used in the main analyses: context-only 0.3295 ± 0.0127, fear memory 0.3617 ± 0.0216, diff = +0.0322, *p* = 0.007). Lines show the group mean and shading the ± s.d. across animals (context-only n = 8, fear memory n = 9). The dashed vertical line marks 10 dimensions, the dimensionality used in the main analyses. The dimensionality sweep and aggregation are as in **a**. (**e**) Overall representational dimensionality did not predict CCA alignment strength within animals (sign-flip permutation test on per-animal correlations, 10,000 two-tailed resamples, n = 17 animals. Overall: mean within-animal Spearman ρ = 0.138, *p* = 0.13. Per session-pair type, BH-FDR-corrected across the three types: Cross-context, mean ρ = +0.118, *p* = 0.28; Ctx-A → Ctx-A, mean ρ = +0.189, *p* = 0.11; Ctx-B → Ctx-B, mean ρ = −0.027, *p* = 0.86). Each point is one session pair from one animal, coloured by session-pair type (Ctx-A → Ctx-B, Ctx-A → Ctx-A, Ctx-B → Ctx-B). The x-axis shows the mean cumulative variance explained by the first 10 PCs, averaged across the two sessions of each pair, and the y-axis the mean canonical correlation across the first 10 modes, aggregated as in **Figure 5c**. A linear fit across all session-pair types is shown for visualization. The Spearman ρ displayed in the panel is based on pooled session-pair data (ρ = 0.00), whereas statistical inference was performed on the per-animal correlations. (**f**) Mismatch in the variance captured by the first 10 PCs between the two sessions of a pair did not predict CCA alignment strength within animals (sign-flip permutation test on per-animal correlations, 10,000 two-tailed resamples, n = 17 animals. Overall: mean within-animal Spearman ρ = −0.057, *p* = 0.15. Per session-pair type, BH-FDR-corrected across the three types: Cross-context, mean ρ = −0.008, *p* = 0.89; Ctx-A → Ctx-A, mean ρ = −0.026, *p* = 0.89; Ctx-B → Ctx-B, mean ρ = −0.226, *p* = 0.13). Each point is one session pair from one animal, colored by session-pair type (Ctx-A → Ctx-B, Ctx-A → Ctx-A, Ctx-B → Ctx-B). The x-axis shows the absolute difference in cumulative variance explained by the first 10 PCs between the two sessions of each pair, and the y-axis the mean canonical correlation across the first 10 modes, aggregated as in **Figure 5c**. A linear fit across all session-pair types is shown for visualization. The Spearman ρ displayed in the panel is based on pooled session-pair data (ρ = −0.07), whereas statistical inference was performed on the per-animal correlations. (**g**) Mismatch in overall dimensionality, defined as the number of PCs required to explain 85% of the variance, did not predict CCA alignment strength within animals (sign-flip permutation test on per-animal correlations, 10,000 two-tailed resamples, n = 17 animals. Overall: mean within-animal Spearman ρ = −0.066, *p* = 0.10. Per session-pair type, BH-FDR-corrected across the three types: Cross-context, mean ρ = −0.029, *p* = 0.62; Ctx-A → Ctx-A, mean ρ = −0.055, *p* = 0.54; Ctx-B → Ctx-B, mean ρ = −0.125, *p* = 0.54). The x-axis shows the absolute difference in dimensionality between the two sessions of each pair. Panel as in **f**. The Spearman ρ displayed in the panel is based on pooled session-pair data (ρ = −0.05).

**Figure S14:**
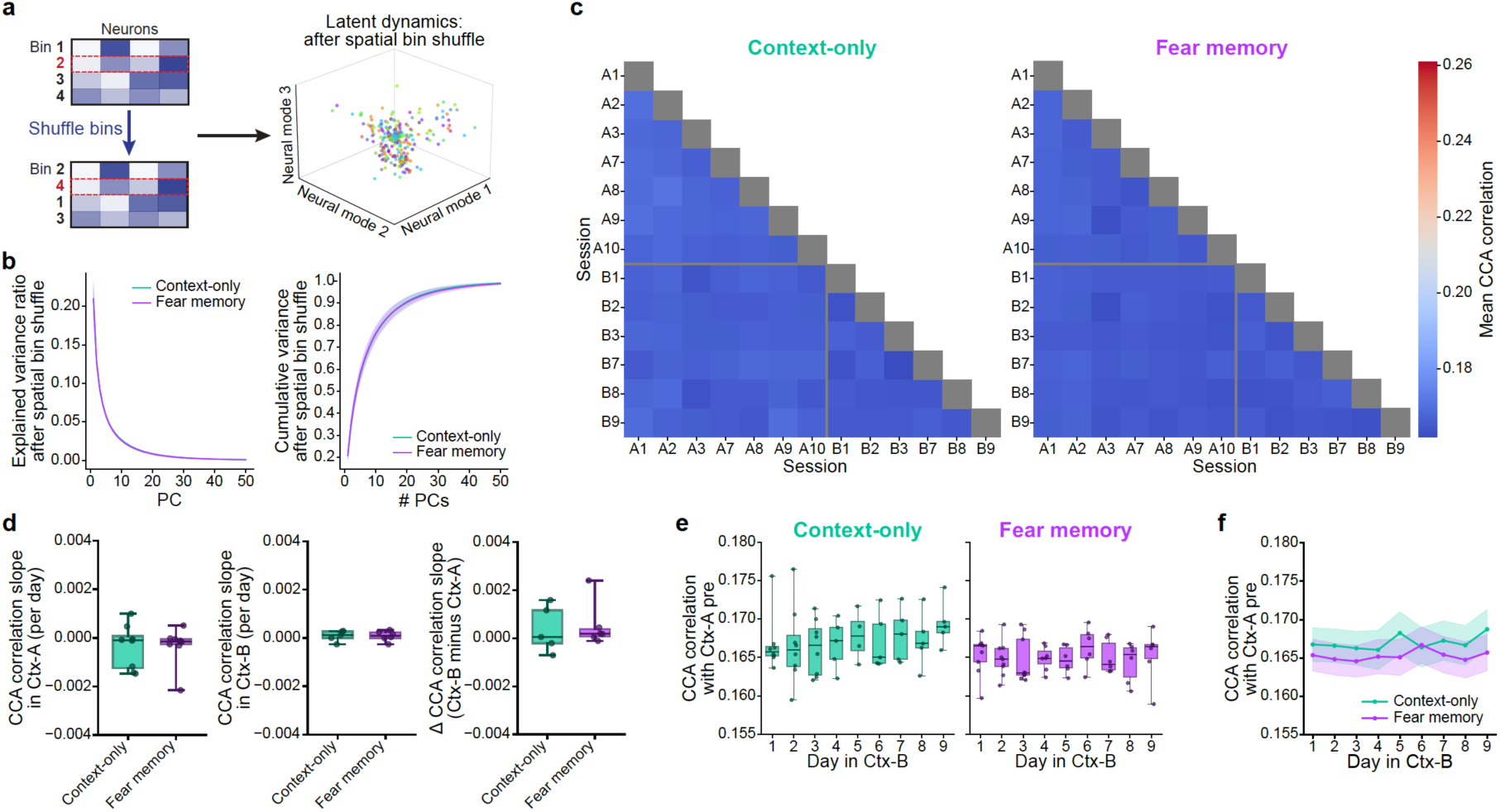
Shuffling spatial bin labels preserves dimensionality but abolishes cross-context alignment differences. (**a**) Schematic of the spatial bin shuffle (left) and the resulting latent dynamics for an example session (right). Left, spatial bin labels were randomly permuted within the session, so that each population vector and the neuron-to-neuron covariance structure are preserved while the correspondence between population activity and physical location is destroyed. Right, the first three neural modes (principal components) of the shuffled spatial population vectors, with each point colored by spatial bin (see colormap in **Figure 5a**). Shuffling removes spatial tuning while leaving overall activity statistics intact, isolating the contribution of spatially organized activity to the low-dimensional structure. (**b**) The dimensionality of spatial population activity after shuffling spatial bin labels did not differ between context-only and fear memory animals. Neither the explained variance of individual principal components nor the cumulative variance differed between groups at any component (independent permutation tests, 10,000 two-tailed resamples, BH-FDR-corrected across the 50 components, all *p* ≥ 0.88 for both metrics), and the number of PCs required to reach 85% cumulative variance was also comparable (independent permutation test, 10,000 two-tailed resamples. Diff is fear memory minus context-only. Diff = −0.16 PCs, d = −0.08, *p* =0.98; context-only: median 15, mean 15.38 ± 1.51; fear memory: median 15, mean 15.22 ± 2.05). Left, mean explained variance ratio per principal component. Right, mean cumulative variance ratio as a function of the number of PCs included. Shading shows ± s.d. across animals. Bin shuffling is as in **a**; subsampling and aggregation as in **Figure 5b**. (**c**) Similarity between latent dynamics across session pairs after shuffling spatial bin labels, shown for context-only (left, n = 8) and fear memory (right, n = 9) animals. Color indicates the mean canonical correlation between the two sessions of each pair, with warmer colors indicating greater similarity. Both heatmaps share a common color scale spanning the same dynamic range as the corresponding analysis in **Figure 5d** but anchored to the minimum correlation observed in the shuffled data, illustrating the marked reduction in similarity structure after shuffling. Only the early and late sessions of each context are shown (Ctx-A: A1–A3 and A7–A10; Ctx-B: B1–B3 and B7–B9), and the thick lines separate the two contexts. Only the lower triangle is shown; the diagonal is omitted, as within-session comparisons were not included. Bin shuffling is as in **a**; subsampling and CCA were performed as in **Figure 5c**, and for each session pair the ten canonical components were averaged into a single similarity value per animal, with heatmaps showing the mean across animals within each group. (**d**) After shuffling spatial bin labels, the change in CCA alignment slope from Ctx-A to Ctx-B no longer differed between groups (independent permutation tests, 10,000 two-tailed resamples. Diff is fear memory minus context-only. Slope in Ctx-A: context-only −0.00035 ± 0.00091, fear memory −0.00030 ± 0.00073, diff = +0.00005, d = +0.07, *p* = 0.91. Slope in Ctx-B: context-only 0.00008 ± 0.00023, fear memory 0.00008 ± 0.00021, diff = +0.00000, d = +0.003, *p* = 0.99. Δ: context-only 0.00038 ± 0.00096, fear memory 0.00053 ± 0.00094, diff = +0.00015, d = +0.16, *p* = 0.82). The absence of a group difference was confirmed with a linear mixed model fitted to the raw session-pair correlations, which allowed animals with partial session coverage to be included (correlation ∼ day × group × context, with a random intercept and a random slope for day per animal; n = 17 animals, 231 observations), in which the three-way interaction was not significant (*p* = 0.498). Left, per-animal slope of cross-session canonical correlation on session day with Ctx-A (context-only n = 8, fear memory, n = 9). Middle, the same within Ctx-B (context-only n = 5, fear memory n = 6). Right, their difference (Δ = slope in Ctx-B minus slope in Ctx-A; context-only n = 5, fear memory n = 6). Bin shuffling is as in **a**. Slopes were estimated and aggregated as in **Figure 5e**. Boxes show the group median and interquartile range, whiskers the full range, and each dot is one animal. (**e**) After shuffling spatial bins labels, alignment to Ctx-A pre across days in Ctx-B did not change in either group (linear mixed-effects model fitted within each group, with log(day) as fixed effect and animal as random intercept, ML; omnibus likelihood-ratio test against an intercept-only model. Context-only: χ²(1) = 0.83, *p* = 0.36, n = 8 animals. Fear memory: χ²(1) = 0.04, *p* = 0.83, n = 9 animals). Per-day contrasts were not tested, as neither omnibus test was significant. Left, context-only animals; right, fear memory animals. Bin shuffling is as in **a**. Each dot is one animal, boxes show the group median and interquartile range, whiskers the full range. Values are aggregated as in **Figure 5f**. (**f**) After shuffling spatial bin labels, alignment to Ctx-A pre across days in Ctx-B no longer differed between groups (linear mixed-effects model, correlation ∼ log(day) × group with animal as random intercept, ML; likelihood-ratio test of the day × group interaction. χ²(1) = 0.21, *p* = 0.65; the interaction term corresponds to a difference in slopes between groups of 0.0003 per log-day, 95% CI [−0.0009, 0.0015]). Groups did not differ on day 1 (a priori contrast, reported uncorrected. Diff is fear memory minus context-only. Diff = +0.0014, *p* = 0.36) or on any subsequent day (BH-FDR-corrected across days 2–9, all *p* ≥ 0.33). Lines show the estimated marginal means and shading the 95% CI, from a linear mixed-effects model with day as a categorical fixed effect crossed with group and animal as random intercept, REML; confidence intervals are from a t-approximation with degrees of freedom equal to the number of observations minus the number of fixed-effect parameters. Bin shuffling is as in **a**. Animal counts vary across days because of incomplete session coverage (n = 4–9 per group per day). Correlation values were aggregated as in **Figure 5f**.

**Figure S15:**
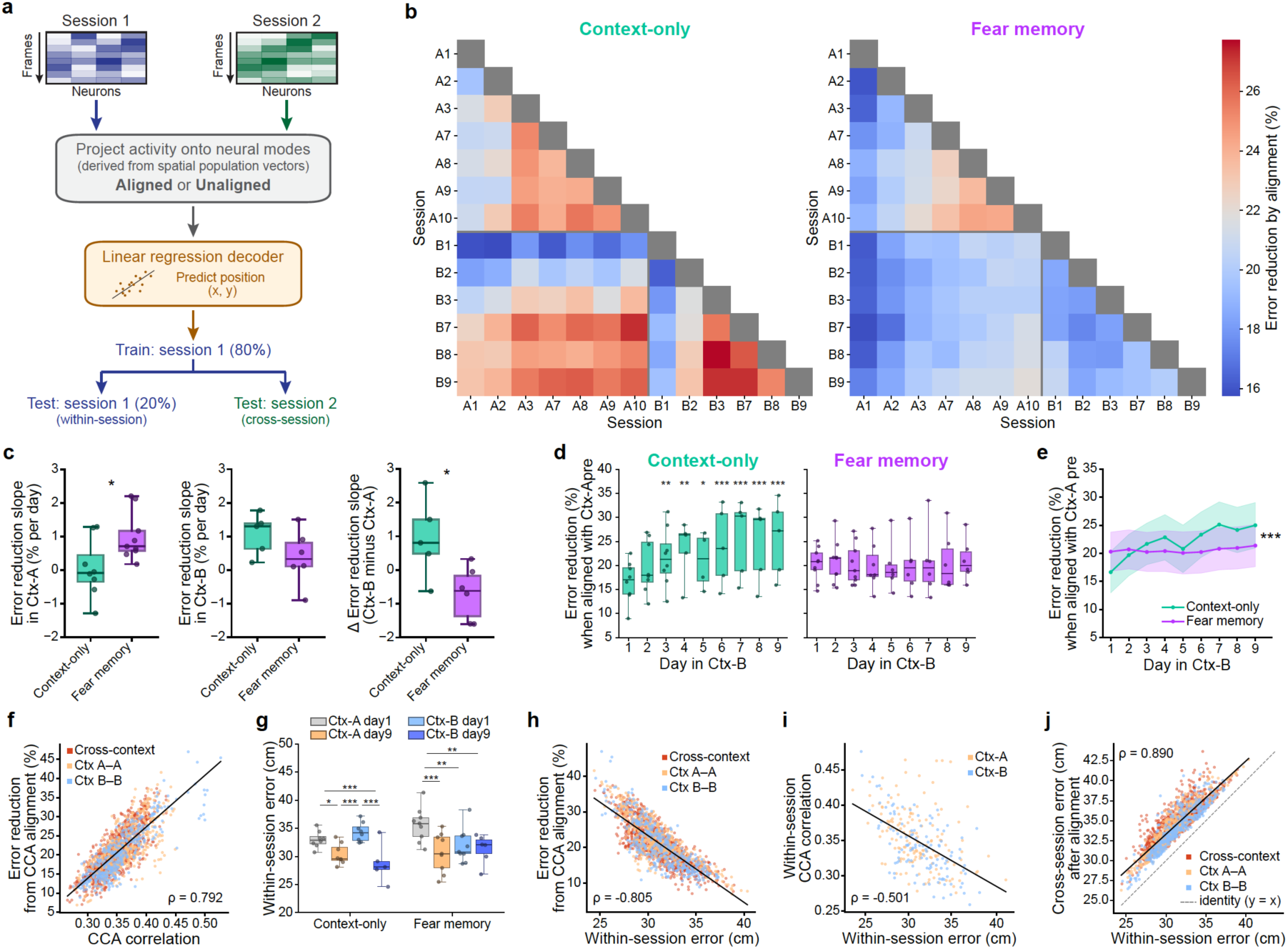
CCA-aligned latent dynamics support cross-session decoding that tracks within-session performance. (**a**) Schematic of the cross-session decoding analysis. For each pair of sessions, per-frame neural activity (frames × neurons) is projected onto neural modes derived from the binned spatial population vectors, either the unaligned modes (PCA only) or the CCA-aligned modes shared across the two sessions. A linear decoder is trained to predict two-dimensional position (x, y) from 80% of Session 1 frames and evaluated on held-out Session 1 frames (20%, within-session) and on all Session 2 frames (cross-session). The CCA alignment is computed from the binned spatial population vectors of the two sessions and uses no position labels; the decoder is trained on Session 1 positions only. (**b**) Improvement in cross-session position decoding from CCA alignment, shown for context-only (left, n = 8) and fear memory (right, n = 9) animals. Each cell shows the percentage reduction in cross-session decoding error (median Euclidean distance in cm between predicted and true position) when session modes were CCA-aligned rather than unaligned, as in **a**, computed as (unaligned − aligned) ÷ unaligned × 100 for each pair of sessions; warmer colors indicate a larger reduction, i.e. greater benefit from alignment. Both matrices share a common color scale. Decoding is directed (train → test); the two directions were computed and then averaged, so each matrix is symmetric. Only the early and late sessions of each context are shown (Ctx-A: A1–A3 and A7–A10; Ctx-B: B1–B3 and B7–B9), and the grey lines separate the two contexts. Only the lower triangle is shown; the diagonal is omitted, as within-session decoding was excluded. Each cell is the mean across animals of the per-animal value, itself the median across 50 subsampling iterations. (**c**) The across-day change in decoding improvement from CCA alignment differed between groups by context, mirroring the alignment-slope result in **Figure 5e** by an independent decoding-based measure (independent permutation tests, 10,000 two-tailed resamples. Diff is fear memory minus context-only, in percentage points of error reduction per day. Slope in Ctx-A: context-only 0.06 ± 0.87, fear memory 1.00 ± 0.72, diff = +0.94, *p* = 0.027. Slope in Ctx-B: context-only 1.07 ± 0.62, fear memory 0.38 ± 0.82, diff = −0.69, *p* = 0.16. Δ: context-only 0.95 ± 1.19, fear memory −0.69 ± 0.79, diff = −1.64, *p* = 0.026). The Δ result was confirmed with a linear mixed model fitted to the raw session-pair values, which allowed animals with partial session coverage to be included (percentage error reduction ∼ day × group × context, with a random intercept and a random slope for day per animal; n = 17 animals, 231 observations), in which the three-way interaction was significant (*p* = 0.020). Left, per-animal slope of decoding error reduction on session day within Ctx-A (context-only n = 8, fear memory n = 9). Middle, the same within Ctx-B (context-only n = 5, fear memory n = 6). Right, their difference (Δ = slope in Ctx-B minus slope in Ctx-A; context-only n = 5, fear memory n = 6). Slopes were estimated per animal by weighted linear regression across sequential session pairs (lag 1 day), each day bin weighted by the number of contributing pairs. Boxes show the group median and interquartile range, whiskers the full range, and each dot is one animal. (**d**) The reduction in cross-session decoding error from aligning Ctx-B sessions with Ctx-A pre increased across days in context-only animals but did not change in fear memory animals, mirroring the alignment result in **Figure 5f** (linear mixed-effects model fitted within each group, with log(day) as fixed effect and animal as random intercept, ML; omnibus likelihood-ratio test against an intercept-only mode. Context-only: χ²(1) = 27.4, *p* < 0.001, n = 8 animals, 53 observations. Fear memory: χ²(1) = 0.44, p = 0.51, *n* = 9 animals, 64 observations). In context-only animals, post-hoc contrasts of each day against day 1 (categorical-day model, REML; BH-FDR-corrected across the eight contrasts. Δ is in percentage points of error reduction relative to day 1) were significant from day 3 onwards (day 3, Δ = +5.04, *p* = 0.002; day 4, Δ = +6.21, *p* = 0.001; day 5, Δ = +4.16, *p* = 0.041; day 6, Δ = +6.68, *p* < 0.001; day 7, Δ = +8.48, *p* < 0.001; day 8, Δ = +7.52, *p* < 0.001; day 9, Δ = +8.36, *p* < 0.001) but not on day 2 (Δ = +3.03, *p* = 0.052). Per-day contrasts were not tested in fear memory animals, as the omnibus test was non-significant. Left, context-only animals; right, fear memory animals. For each animal and session pair, the percentage reduction in cross-session decoding error from CCA-aligning a Ctx-B session with Ctx-A pre, (unaligned − aligned) ÷ unaligned × 100, was averaged across both decoding directions and taken as the median across 50 subsampling iterations. Each dot is one animal, boxes show the group median and interquartile range, whiskers the full range. (**e**) The reduction is cross-session decoding error from aligning Ctx-B sessions with Ctx-A pre increased across days in context-only animals but not in fear memory animals, so that the groups diverged over days rather than differing on any single day, mirroring the alignment result in **Figure 5g** (linear mixed-effects model, reduction ∼log(day) × group with animal as random intercept, ML; likelihood-ratio test of the day × group interaction. χ²(1) = 21.4, *p* < 0.001. The interaction term gives the difference in slopes between groups: the error reduction increased 3.45 percentage points per log-day more steeply in context-only than in fear memory animals, 95% CI [2.02, 4.88]). Groups did not differ on day 1 (a priori contrast, reported uncorrected. Diff is Fear memory minus Context-only. Diff = +3.65, *p* = 0.15) or on any subsequent day (BH-FDR-corrected across days 2–9, all *p* ≥ 0.53). Lines show the estimated marginal means and shading the 95% CI, from a linear mixed-effects model with day as a categorical fixed effect crossed with group and animal as random intercept, REML. Reduction values were computed as in **d**. Stars denote the omnibus day × group interaction. (**f**) CCA alignment strength was strongly and positively correlated with the reduction in cross-session decoding error within animals, confirming that the two measures capture the same underlying alignment (sign-flip permutation test on per-animal correlations, 10,000 two-tailed resamples, n = 17 animals. Overall: mean within-animal Spearman ρ = 0.656, *p* < 0.001. Per session-pair type, BH-FDR-corrected across the three types: cross-context, mean ρ = +0.613, *p* < 0.001; Ctx-A → Ctx-A, mean ρ = +0.549, *p* < 0.001; Ctx-B → Ctx-B, mean ρ = +0.590, *p* < 0.001). Each point is one session pair from one animal (n = 2,046 pairs), colored by session-pair type (Ctx-A → Ctx-B, Ctx-A → Ctx-A, Ctx-B → Ctx-B). The x-axis shows CCA alignment strength, the mean canonical correlation across the first 10 modes, and the y-axis the percentage reduction in decoding error from CCA alignment, (unaligned − aligned) ÷ unaligned × 100, averaged across both decoding directions; both are taken as the median across 50 subsampling iterations. A linear fit across all session-pair types is shown for visualization. The Spearman ρ displayed in the panel is based on pooled session-pair data (ρ = 0.79), whereas statistical inference was performed on the per-animal correlations. (**g**) Within-session decoding error decreased across familiarization in both contexts in context-only animals, but in fear memory animals this improvement was absent in Ctx-B (linear mixed-effects model with session as a categorical fixed effect and animal as random intercept, REML; pairwise contrasts BH-FDR-corrected across all comparisons. Δ is in cm, positive values indicating lower error on the later session. Context-only: Ctx-A day1 → day9, Δ = +2.64, *p* = 0.012; Ctx-B day1 → day9, Δ = +5.05, *p* < 0.001; Ctx-A day9 → Ctx-B day1, Δ = −3.94, *p* < 0.001; Ctx-A day1 → Ctx-B day9, Δ = +4.41, *p* < 0.001. Fear memory: Ctx-A day1 → day9, Δ = +4.95, *p* < 0.001; Ctx-A day1 → Ctx-B day1, Δ = +3.78, *p* = 0.002; Ctx-A day1 → Ctx-B day9, Δ = +4.91, *p* = 0.001; Ctx-B day1 → day9, Δ = −0.02, *p* = 0.80, not significant). The absence of improvement across Ctx-B in fear memory animals contrasts with the corresponding improvement in Ctx-A of the same animals (Δ = +4.95) and with Ctx-B in context-only animals (Δ = +5.05). All remaining contrasts had *p* ≥ 0.19. The model was chosen to handle the repeated-measures structure and missing data at later timepoints (attrition due to logistical constraints; missing-at-random assumed); sensitivity analyses (bootstrap, permutation, paired t-tests, Wilcoxon signed-rank) on complete-case pairs were consistent for the well-powered comparisons. Each dot is one animal; boxes show the group median and interquartile range, whiskers the full range. (**h**) Within-session decoding error was strongly and negatively correlated with the reduction in cross-session decoding error from CCA alignment within animals: alignment helped most for session pairs whose own within-session decoding was already good, and least where within-session decoding was poor (sign-flip permutation test on per-animal correlations, 10,000 two-tailed resamples, n = 17 animals. Overall: mean within-animal Spearman ρ = −0.659, *p* < 0.001. Per session-pair type, BH-FDR-corrected across the three types: cross-context, mean ρ = −0.636, *p* < 0.001; Ctx-A → Ctx-A, mean ρ = −0.629, *p* < 0.001; Ctx-B → Ctx-B, mean ρ = −0.510, *p* = 0.001). Each point is one session pair from one animal (n = 2,046 pairs), colored by session-pair type (Ctx-A → Ctx-B, Ctx-A → Ctx-A, Ctx-B → Ctx-B). The x-axis shows the mean within-session decoding error across the two sessions of each pair, and the y-axis the percentage reduction in cross-session decoding error from CCA alignment. A linear fit across all session-pair types is shown for visualization. The Spearman ρ displayed in the panel is based on pooled session-pair data (ρ = −0.81), whereas statistical inference was performed on the per-animal correlations. (**i**) Within-session decoding error was negatively correlated with within-session CCA alignment within animals: sessions with better within-session decoding also showed higher alignment between their two halves (sign-flip permutation test on per-animal correlations, 10,000 two-tailed resamples, n = 17 animals. Overall: mean within-animal Spearman ρ = −0.442, *p* < 0.001. Per context, BH-FDR-corrected across the two contexts: Ctx-A, mean ρ = −0.185, *p* = 0.073; Ctx-B, mean ρ = −0.386, *p* = 0.020). Each point is one session from one animal (n = 268 sessions), colored by context (Ctx-A, Ctx-B). The x-axis shows the within-session decoding error (cm) and the y-axis the within-session CCA alignment, the mean canonical correlation across the first 10 modes between the first and second half of the same session; both are taken as the median across 50 subsampling iterations. A linear fit across both contexts is shown for visualization. The Spearman ρ displayed in the panel is based on pooled session-level data (ρ = −0.50), whereas statistical inference was performed on the per-animal correlations. (**j**) After CCA alignment, cross-session decoding error closely tracked each pair’s within-session decoding error, remaining a small consistent margin above it (sign-flip permutation test on per-animal correlations, 10,000 two-tailed resamples, n = 17 animals. Overall: mean within-animal Spearman ρ = 0.809, *p* < 0.001. Per session-pair type, BH-FDR-corrected across the three types: cross-context, mean ρ = +0.760, *p* < 0.001; Ctx-A → Ctx-A, mean ρ = +0.862, *p* < 0.001; Ctx-B → Ctx-B, mean ρ = +0.707, *p* < 0.001). Aligned cross-session error exceeded within-session error by a small margin in every pair (median 2.98 cm, above zero for all 2,046 pairs), so alignment approached but did not fully reach within-session performance. Each point is one session pair from one animal, colored by session-pair type (Ctx-A → Ctx-B, Ctx-A → Ctx-A, Ctx-B → Ctx-B); the dashed line is the identity (y = x). The x-axis shows the mean within-session decoding error across the two sessions of each pair, and the y-axis the cross-session decoding error after alignment, both averaged across decoding directions and taken as the median across 50 subsampling iterations. A linear fit across all session-pair types is shown for visualization. The Spearman ρ displayed in the panel is based on pooled session-pair data (ρ = 0.89), whereas statistical inference was performed on the per-animal correlations.

## Methods

### Animals

All procedures related to animal housing, surgery, behavioral experiments and euthanasia were conducted according to the guidelines of the Cantonal Veterinary Office Basel-Stadt, and in compliance with the Swiss Veterinary Law. Approval for all experiments that involved living animals were obtained before the start of the project under permit number 3018. Only male mice were used for this study. Experiments were performed on C57BL/6JRj (Janvier). Mice were initially group housed in temperature– and humidity-controlled cages and kept on a 12h light-dark cycle. They were then single-housed at least 1 week prior to the start of the experiment, and food-restricted ±5 days prior. After this and until the end of the experiment, mice underwent a daily health inspection, and their body weight was kept above 80% of their initial weight. The researchers responsible for the mice assigned them to experimental groups based on the quality of their calcium imaging recordings, taking into account viral expression and signal strength, so that each group contained animals with comparable recording quality. All experiments took place during the dark phase of the 12h light-dark cycle.

### Anesthesia and analgesia during surgeries

In adult mice, Buprenorphine (Bupax P, 0.1 mg/kg, Streuli) and Atropine Sulphate (0.05 mg/kg, Amino AG) were injected subcutaneously (s.c.) before the surgery as pre-emptive analgesia. Mice were then anesthetized with isoflurane in oxygen (Piramal, 3% at induction, 1-2% for maintenance) at an airflow of 0.5 l/min. Bupivacaine (1-5 mg/kg, Sintetica) and Lidocaine (< 7 mg/kg, Streuli) were administered locally through a s.c. injection at the start of the surgery. After surgery completion, mice could recover under a heat lamp and Buprenorphine was administered 4 to 6 hours later. Meloxicam (Metacam 5 mg/kg, Boehringer Ingelheim) was injected s.c. once a day on the following 3 days as postoperative analgesia.

### Viral injections

A viral vector to induce expression of the genetically encoded calcium indicator GCaMP6f (pAAV.Syn.GCaMP6f.WPRE.SV40, Cat. Number #100837-AAV1, titer 2 x 10^13^, Addgene^77^) was injected stereotaxically (Kopf) into the hippocampus of the right hemisphere. Injection took place between postnatal day 39 and 87 (±4 weeks before the start of the experiment), at stereotaxic coordinates: –1.58 mm AP, +1.65 mm ML and –1.7/-1.5 mm DV (276nL at each DV location) from Bregma. Mice were then allowed to recover for at least a week.

### Lens implants and baseplating

After stereotaxic alignment under isoflurane anesthesia, an endoscope comprised of a tubular, 1 mm wide GRIN lens glued to a 1 mm prism (custom microendoscope design, GRINtech GmbH, Jena, Germany (see Kveim et al., 2024^8^)) was inserted through a 1.5 mm wide circular craniotomy to gain optical access to the CA3 area of the hippocampus. Implantation took place between postnatal days 48 and 94 (at least 13 days before the start of the experiment). The imaging face of the prism was implanted at a 45° angle facing the posterior-medial part of the brain so that it follows the curvature of the hippocampus, with its most frontal edge at the following stereotaxic coordinates: –1.58 mm AP, +1.95 to +2.00 mm ML and –2.3 mm DV from Bregma. After insertion into the tissue, the endoscope was moved –0.1 mm ML to compress tissue against the imaging face of the prism, thereby limiting movement along the z-axis during imaging. UV-curable cement (Venus Diamond Flow, Kultzer) was used to secure the lens to the skull and dental cement (Paladur, Kulzer) mixed with charcoal powder (Graphite, Carl Roth) was applied over the exposed skull for protection and to secure a metal head bar that was leveled with the surface of the endoscope. After ±1.5 to 2 weeks of recovery, mice were imaged under isoflurane anesthesia using a 1 photon microscope (nVista 3.0 or nVue 1.0, Inscopix). When a mouse exhibited homogeneous GCaMP6f expression and appeared to have healthy tissue, a baseplate was permanently fixed to the implant with UV-curable cement and superglue (Superglue 415, Loctite, Loctite) to enable easy mounting of a miniaturized, 1-photon microscope (Inscopix). The baseplate was shielded with dental cement mixed with charcoal powder and a magnetic cap was placed on top to protect the lens in between imaging sessions.

### Calcium imaging

An integrated miniature fluorescence 1-photon microscope (nVista 3.0 or nVue 2.0, Inscopix) was used to perform time-lapse imaging of calcium activity in CA3 neurons from freely behaving mice. At the start of each imaging session, the miniscope was mounted on the baseplate while the animal was head-fixed to a stable head-post and could freely run on a planar wheel. Calcium imaging was performed at a rate of 20Hz and a resolution of 1200 x 800 pixels, thereby covering a field of view of 1000 x 625 μm (1 pixel = 0.765 μm^2^). On the first experimental day, imaging parameters such as LED brightness, gain and focus plane were empirically optimized (Supplementary Table 1) to capture the maximum number of neurons. LED brightness and gain remained constant throughout the experiment, while the focus plane was reassessed and when necessary, adjusted before each session to counteract small drifts along the z-axis in between days. To record mouse behavior, an overhead camera (Basler ace acA2040-55 μm, BASLER AG) was used that was synchronized with the integrated microscope.

### Behavioral task

Mice were food restricted and habituated to human handling and miniscope mounting for 5 days prior to the experiment. They were then imaged for 9 consecutive days, one 30-minute session per day, during which they randomly foraged for reward (cookie crumbs) in an open field arena of 80×80 cm (Ctx-A). On day 10, mice underwent fear conditioning in one of two protocols depending on the cohort they belonged to: 5 mice in cohort one were imaged with the miniscope during fear conditioning, while another 4 mice in cohort two were not imaged. In the first cohort, mice were first imaged in their home cage for 10 minutes and then, without removing the miniscope in between recordings, underwent an acquisition session (hardware: Fear Conditioning system, Ugo Basile; software: EthoVision XT, version 14/17, Noldus) in whey were first allowed to freely explore a circular arena of 39 cm diameter (Training Context, Conditioned Stimulus, CS) characterized by distinctive visual wall patterns and an electrifiable grid floor, but no specific odor. After 3 minutes, a series of 5 footshocks (1 s, 0.8 mA) with 30-second intervals was delivered (Unconditioned Stimulus, US). In the second cohort, mice underwent the same acquisition session in 25×25 cm square box (Training Context, Conditioned Stimulus, CS) that had a similar wall pattern and floor as the circular arena but additionally carried a specific odor (2% acetic acid). In addition to these two contextual fear conditioning (cFC) cohorts, separate cohorts of mice underwent the same acquisition session without foot shocks (context-only control; 4 mice in cohort one imaged with the miniscope and 4 mice in cohort two that were not imaged) or an immediate shock (IS) protocol that does not support hippocampus-dependent contextual learning (IS control; 4 mice imaged with the miniscope)^12^. During the IS protocol, mice received 5 footshocks (1 s, 0.8mA) at seconds 2, 8, 14, 20 and 26 and then remained in Training Context for another 60 seconds. Between 6 and 8 hours (Supplementary Table 1) after the acquisition session in the Training Context, on day 10, all mice were reintroduced to Ctx-A and imaged during 30 minutes of random foraging. They were then placed back into their home cage for 1 to 3 hours (Supplementary Table 1) before being introduced to a different open field arena (Ctx-B) that has the same size as Ctx-A but different wall color, spatial cue color, lightning and floor texture. Imaging in Ctx-B also lasted 30 minutes in which mice were randomly foraging for reward (sugary sprinkles). On day 11, all mice underwent a retrieval session, during which they were placed back into the Training Context and allowed to freely explore it for 5 minutes without presentation of the US. Memory retention and expression were quantified as the amount of time the animal spent freezing upon exposure to the CS during this retrieval session. Freezing bouts were systematically identified by the Ethovision XT software and defined as instances of complete immobility apart from breathing (>2 seconds of pixel change <1-3%, adjusted empirically for each mouse). Between 6 and 8 hours after the retrieval session, on day 11, mice were imaged again Ctx-A and Ctx-B, following the same imaging schedule for each mouse as on day 10. They were then similarly imaged in these 2 open field arenas for 7 more days.

### Processing of calcium imaging recordings

Raw miniscope recordings were processed using the Inscopix data analysis software Matlab API (IDAS, version 1.6.0, Inscopix), as well as custom Matlab and Python scripts (version R2021b, Mathworks; version 3.8.12, Python). Recordings acquired during the same session (i.e., when the miniscope was not unmounted in between recordings) were concatenated. Spatial down-sampling by a factor of 2 resulted in a final field of view of 640 x 400 pixels (1 pixel = 1.530 μm^2^), which was then cropped and aligned to improve cell registration across sessions by matching landmarks such as blood vessels and lens corners. Recorded frames were filtered with a Gaussian band-pass filter (high cut-off: 0.500/pixel, low cut-off: 0.005/pixel) and motion registered on a frame-by-frame basis.

*Cell detection.* Constrained Nonnegative Matrix Factorization for micro-endoscopic data (CNMFe, provided by IDAS) was used to identify regions of interest (ROIs) corresponding to individual neurons. Default IDAS values were used, except for the minimum pixel correlation (0.917 ± 0.006) and minimal peak-to-noise ratio (18.095 ± 0.843), which were empirically assessed on a mouse-to-mouse basis. The cell size was set to 12 pixels in diameter (18.36 μm^2^) after down-sampling. ROIs were obtained by delineating a boundary that induced only pixels with average fluorescence values within the highest 20% of the maximum intensity per ROI. For each ROI, IDAS extracted fluorescent traces expressed as ΔF/σ (i.e., the change in fluorescence relative to the baseline noise, σ).

*Event detection.* To identify calcium events, fluorescence traces were pre-processed by applying a first-order Butterworth low-pass filter (cut-off 0.5-1Hz) and first-order polynomial detrending to remove slow drifts and bleaching effects. All thresholds were applied to filtered and detrended fluorescence traces expressed in normalized units (scaled representations of the ΔF/σ signal). Cells with excessive baseline variability, defined as having a standard deviation exceeding a dataset-specific threshold (0.050 ± 0.002), were excluded from further analysis. The remaining cells were evaluated with an additional quality-control step based on a moment statistic (0.019 ± 0.003) that quantifies the shape of the fluorescence distribution. This criterion was used to identify cells with atypical signal properties (e.g., non-physiological skewness or heavy-tailed noise) that were not captured by variance-based exclusion. Event detection thresholds were computed individually for each cell by fitting a Gaussian distribution to the filtered trace and selecting a high percentile (e.g. 99.999^th^ percentile) to define candidate calcium transients relative to the cell-specific activity distribution. Parameter values were empirically optimized on a per-dataset basis to maximize detection of visually identifiable calcium transients while minimizing noise-driven false positives, and were validated for consistency across recordings. Calcium events were defined as periods during which the fluorescence trace exceeded both this percentile-based threshold and a minimum normalized fluorescence criterion (0.142 ± 0.010 for typical cells; for cells with atypical signal properties exceeding the moment threshold, a more conservative fixed threshold of 0.500 was applied). Detected events were further processed to distinguish the “up-phase” (rising phase of the event, defined as frames with a positive derivative of the filtered calcium signal) and the “on-phase” (entire duration of the event). Only events with continuous durations of at least 0.3 s for the up-phase and 0.5 s for the on-phase were retained. Calcium traces were binarized based on up-phase activity, with up-phase frames assigned a value of 1 and all other frames assigned a value of 0.

*Cell curation.* The signal-to-noise ratio (SNR) for each cell was calculated as the ratio of the mean event amplitude to the standard deviation of the noise. Noise statistics were estimated from the filtered ΔF/σ signal in temporal windows extending from 1 s before to 10 s after each up-phase event, excluding the up-phase itself, to capture local baseline fluctuations surrounding detected events. SNR thresholds for cell inclusion were defined based on the lowest 4-8% (6.95 ± 0.21) of SNR values within each recording and empirically optimized on a per-animal basis to exclude false positives and account for differences in recording quality. To prevent redundant sampling of spatially overlapping signals, pairwise distances between all ROIs were computed, and when two ROIs were separated by less than 3-5 pixels (4.05 ± 0.08), only the ROI with the higher SNR was retained for further analysis. Cells were additionally required to exhibit an average activity rate of at least 0.01Hz (i.e., active frames per second) to be included.

*Longitudinal tracking.* Longitudinal registration of neurons across sessions was conducted using dedicated algorithms provided by IDAS. A minimum correlation threshold of 0.6 was set to ensure reliable tracking between sessions. Tracking performance was evaluated by comparing the distributions of normalized cross-correlation scores and centroid distances across animals to verify consistency and exclude mismatched neuron assignments. Longitudinal registration was applied across all imaging sessions required for each analysis, including consecutive-day recordings (days 1–3 and days 7–9 in Ctx-A/Ctx-B) and the five key behavioral sessions (Ctx-A pre, Acquisition, Ctx-A post, Ctx-B day1, and Recall) used in the longitudinal analyses. For the latter, cohort 1 was registered across all five sessions, whereas cohort 2 was registered across the three available imaging sessions (Ctx-A pre, Ctx-A post, and Ctx-B day1), as these animals were not imaged during Acquisition or Recall. No evidence for group-dependent differences in longitudinal registration quality was observed in either cohort (Figure S2c–e).

### Analyzing mouse behavior in open field arenas

Videos of mouse behavior in the open field arenas were analyzed using a custom-trained neural network in DeepLabCut (version 2.2.3)^78^ that detects the head-neck position in each frame. A custom Python script was then applied to remove frames with low-confidence estimates (< 50% likelihood) and spurious tracking jumps that exceeded a maximum plausible travel speed (> 90 cm/s). These frames were replaced with the median of surrounding coordinates within a ±5-frame window. To further reduce tracking artifacts from rearing or jumping along the wall of the arena, 2D coordinates were filtered using a percentile-based boundary (1^st^-99^th^ percentiles) calculated across all frames. Coordinates outside this range were considered outliers and replaced with the nearest boundary value. To filter residual high-frequency tracking jitter not captured by the global speed-based exclusion criteria, an additional adaptive jump-cleaning step was applied to remove small frame-to-frame coordinate discontinuities indicative of tracking noise. In this step, x– and y-coordinates were evaluated independently to detect outlier displacements using a rolling median and median absolute deviation (MAD) computed over a 15-frame window. Frames with displacements exceeding the local median by more than 1x MAD (z-threshold = 1) were flagged as outliers. Flagged frames were then replaced with the median of the surrounding frames with a ±5-frame window, and the corrected x and y coordinates were recombined into continuous trajectories. Corrected trajectories were used for all subsequent analyses. Velocity (in cm/s) was calculated as the Euclidean distance between consecutive positions and smoothed using a Savitzky-Golay filter (1 s window, second-order polynomial fit). Grooming bouts were identified using a rule-based classifier combining moderate velocity (2-10 cm/s), low displacement (<1.5 cm), low spatial variance (<2.0 cm^2^), and elevated acceleration (> 0.2 cm/s^2^), with parameters empirically chosen and computed over a 10 s moving window. Grooming bouts were post-processed using morphological closing (0.5 s) and retained only if lasting at least 5 seconds. Mobility states (moving vs. stationary) were defined using a velocity threshold of 2 cm/s, with grooming periods assigned to the stationary state. Brief transitions (< 2 s) between mobility states were corrected to reduce fragmentation. Only mobile periods were used in subsequent analyses unless specifically stated otherwise.

### Place cells identification and analysis

To identify place cells, spatial tuning was computed using the Opexebo open-source Python toolbox together with custom scripts. The arena was divided into 5 x 5 cm spatial bins, and neural activity was binned according to the animal’s position to construct a signal map. Neural activity was defined as the filtered ΔF/σ signal gated by up-phase events, such that only fluorescence amplitudes during the rising phase of calcium transients contributed to spatial tuning calculations. Signal maps were smoothed using a Gaussian kernel (σ = 1.75 bins), while occupancy maps were smoothed separately (σ = 2 bins). Smoothed signal maps were then normalized by smoothed occupancy maps to obtain activity rate maps. Spatial bins visited for less than 200ms were masked and excluded from further analyses. Activity rate maps were subsequently subject to light additional smoothing (σ = 0.3 bins), with masked bins preserved to avoid reintroducing data from under-sampled locations. Spatial information was computed from the activity rate maps using the Skaggs information measure^79,80^ and compared to a null distribution obtained by circularly shuffling the activity trace relative to position (1,000 iterations), thereby preserving temporal structure while disrupting spatial alignment. Spatial information values were z-scored relative to the shuffle distribution, and cells were classified as place cells if their z-score exceeded a threshold corresponding to the 99^th^ percentile. Coherence and selectivity of each cell’s rate map were calculated to further characterize spatial tuning. Coherence was defined as the Pearson correlation between each bin and the average of its eight neighboring bins in the raw rate map. Selectivity was calculated as the ratio of the peak activity rate to the mean activity rate from the smoothed rate map. Both measures were computed after masking bins with insufficient occupancy and prior to any further analyses. Place fields were identified as spatially contiguous regions exceeding a seed threshold (≥ 92% of the peak activity rate), with a minimum mean activity of ≥ 12% of the peak activity rate, a minimum peak activity of ≥ 2% of the maximum, and covering at least 12 adjacent spatial bins. Cells with multiple place fields were included. At the population level, field distributions across the arena were quantified for each session by calculating the normalized fraction of fields in each spatial bin, along with field entropy, and the coefficient of variation to capture variability in field locations. To characterize place cell stability and representational drift across sessions, two complementary approaches were used. First, place cell stability was quantified using four metrics computed for each consecutive day pair within early (days 1−3) and late (days 7−9) familiarization periods, and then averaged within each period: (1) retained place cells, defined as the percentage of place cells from day N that remain place cells on day N+1; (2) recruited place cells, defined as the percentage of non-place cells on day N that became place cells on day N+1; (3) prior place cells, defined as the percentage of place cells on day N+1 that were already place cells on day N; and (4) Jaccard similarity, defined as the size of the intersection of place cell populations across consecutive days divided by the size of their union, providing a symmetric measure of population overlap. Second, for each cell classified as a place cell in both sessions, three metrics of representational similarity were computed: (1) centroid distance, defined as the Euclidean distance between the centers of mass of the primary place field across sessions (in cm); (2) rate map correlation, defined as the Pearson correlation between session N and session N+1 activity rate maps restricted to spatial bins visited in both sessions; and (3) population vector correlation, defined as the Pearson correlation between population activity vectors computed for each spatial bin, averaged across all visited bins. Cue card locations in Ctx-A and Ctx-B were kept consistent to ensure a stable spatial reference frame for within– and across-environment comparisons.

### Ensemble neurons identification

For all neurons and sessions, a tuning score (TS) was computed to quantify the extent to which the activity rate of individual neurons was modulated by exposure to the Training Context^8,76^. This was calculated as the difference in activity rate (in Hz) between the Training Context (TC) and baseline activity in the Home Cage (HC), normalized by their sum:

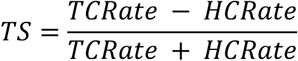

TS values across the recorded population ranged from –1 (neurons exclusively active in HC) to +1 (neurons exclusively active in TC). Neurons with TS ≥ 0.33 were classified as “Activated”, while neurons with TS ≤ –0.33 were identified as “Inhibited”. Neurons with intermediate TS values were classified as “Neutral”. Neurons classified as “Activated” during the acquisition session were defined as acquisition ensemble neurons, those classified as “Activated” during the retrieval session as retrieval ensemble neurons, and neurons classified as “Activated” during both acquisition and retrieval sessions as memory ensemble neurons.

### Significance testing of co-occurrence between neuronal populations

Within-animal permutation approaches were used to test whether observed associations between functionally defined neuronal populations exceeded chance levels. Populations were defined and labeled independently based on task-specific criteria (e.g., place cells and ensemble neurons; see above). For each analysis, labels relevant to the comparison were shuffled independently within each animal, preserving per-animal label frequencies and group sizes while disrupting associations between neuronal classifications (e.g., memory status and spatial map classifications). Unless otherwise specified, analyses were restricted to neurons detected in at least one session relevant to the comparison.

*Monte Carlo enrichment.* For each combination of functionally defined populations, co-occurrence was quantified as the percentage of neurons belonging to that combination across neurons pooled from all animals. Null distributions were generated using either within-animal label shuffling or within-animal random sampling. For label-shuffling analyses, labels relevant to the hypothesis being tested were shuffled independently within each animal (10,000 permutations). For subpopulation-enrichment analyses, cells were randomly sampled without replacement from a reference pool within each animal while preserving subpopulation size. Co-occurrence percentages were recalculated for each iteration to generate the null distribution. An enrichment ratio was calculated as the observed percentage divided by the mean percentage of the null distribution. Z-scores indicate the number of standard deviations by which the observed percentage deviated from the null mean.

*Profile co-occurrence enrichment.* Each neuron was assigned a multi-dimensional profile defined by its labels across multiple sessions. Expected profile frequencies were computed separately for each animal from the marginal probabilities of each neuronal classification, yielding the expected frequency under statistical independence. Observed and expected profile frequencies were compared using a log-enrichment ratio: log((observed + 0.5) / (expected + 0.5)), where 0.5 is a Laplace smoothing constant used to avoid undefined values for profiles with zero counts. Log-enrichment ratios were averaged across animals to obtain a mean log-enrichment ratio for each profile. To generate a null distribution, profile-defining labels were shuffled independently within each animal (10,000 permutations), preserving marginal label frequencies while disrupting associations among neuronal classifications. Following each shuffle, observed profile frequencies were recomputed and compared with the corresponding expected frequencies, and the resulting log-enrichment ratios were averaged across animals to obtain a single null value for each profile. Repeating this procedure across all permutations generated the null distribution.

*Multiple Correspondence Analysis (MCA) and cluster significance.* MCA was applied to binary indicator matrices derived from multi-dimensional neuron profiles to identify low-dimensional structure in label co-occurrence patterns. Each neuron was represented by a binary indicator vector containing one column for each label within each neuronal classification. This resulted in 12 binary columns in total (3 memory labels and 3 spatial labels across 3 sessions). Within each neuronal classification block, exactly one label was assigned a value of 1 and all others a value of 0. MCA identifies dimensions that maximize variance in co-occurrence patterns across neurons. Dimensions with inertia exceeding the average inertia across all dimensions were retained. Ward hierarchical clustering was then applied to the coordinates of each label within each neuronal classification in the retained MCA space. The number of clusters was selected using the Ward linkage elbow criterion, defined as the largest increase in linkage distance between successive merges, which corresponded to four clusters in both groups and was consistent with visual inspection of the dendrogram. Cluster compactness was quantified as the mean pairwise Euclidean distance among points within each cluster in MCA space. To assess significance, the relevant labels were shuffled independently within each animal (10,000 permutations), and MCA was repeated on each shuffled dataset. Within-cluster distances were then recomputed using the original cluster assignments. Cluster boundaries were therefore fixed and were not re-estimated after each permutation. This procedure tests whether the observed clusters are more compact than expected by chance while controlling for marginal label frequencies, thereby testing the strength of the observed associations rather than the existence of clustering per se.

### Bayesian decoder

For the open field arena sessions (Ctx-A and Ctx-B), a custom Naïve Bayes regression model was used to estimate the animal’s position from binarized calcium event data. Mobile frames were split into training (80%) and testing (20%) sets using two strategies: (1) a sequential split that preserves temporal structure, and (2) a random split with shuffled frame order. Frames without any active neurons were excluded from both training and test sets. Using the training data, each neuron’s activity was modeled as a Poisson-distributed variable with activity rates dependent on spatial location. The environment was partitioned into a 16 x 16 spatial grid (5 cm bins). For each neuron, a generalized linear model (GLM) with Poisson family link was fitted to estimate activity rates across all bins. Only neurons with mean activity rates of at least 0.01 Hz in the training set were retained to ensure the decoder was trained on informative population activity. For each test frame, we computed the posterior probability of the animal’s position by combining Poisson log-likelihoods across all neurons and selected the center of the bin with the maximum likelihood position as the decoded location. Decoding error was quantified as the Euclidean distance (in cm) between the decoded and true position at each time point. Test frames where the animal visited locations more than 5 cm outside the training spatial coverage were excluded from analysis to avoid extrapolation errors. To ensure that the decoding error was not influenced by the size of the recorded population, separate analyses performed 50 iterations using a randomly selected subset of 100 neurons, then computing the mean decoding error across iterations. This subsampling approach was used to obtain results unless specified otherwise. As a control, position labels in the training set were circularly shifted by half of the recording length to destroy mapping between neural activity and animal location while preserving the temporal structure and spatial distribution of both signals. Decoding error under this shuffle provides a chance-level baseline.

### Analysis of neural population geometry

A custom pipeline combining principal component analysis (PCA) and canonical correlation analysis (CCA)^68,72^ was developed to characterize the geometry of neural population activity during open field arena sessions in Ctx-A and Ctx-B. Position data were rotated to align the spatial cue across contexts to a common reference direction and spatially binned into a 16 x 16 grid (5 cm bins). Calcium imaging data were smoothed with a Gaussian filter (σ = 3.33 frames, corresponding to a smoothing window of approximately 1 s at 20 Hz). In the main analysis, neuronal activity was analyzed in its original scale, without normalization. To assess whether the results were driven by a small number of highly active neurons, an additional analysis applied z-scoring to each neuron’s activity across time within session as a control. For each session, a random subset of 100 neurons was selected. The full analysis pipeline was repeated over 50 iterations with different subsets, and results were averaged across iterations to reduce sensitivity to neuron subsampling. The average activity of each neuron was computed across all frames in which mice occupied the same spatial bin, producing a population activity matrix (spatial bins x neurons) for each session. The same procedure was applied separately to the first and second halves of each session to enable within-session comparisons, resulting in three population activity matrices per session (full session, first half, second half).

*PCA.* PCA was applied to each spatial population matrix after excluding unvisited bins, projecting the bin-wise population activity onto a lower-dimensional subspace (spatial bins × principal components). To quantify how accurately position could be decoded as a function of dimensionality, a linear regression decoder was trained on mean-centered neural activity projected onto the first *d* principal components (*d* = 1 to 100). Decoder performance was evaluated using a held-out 20% test split within each session and quantified as the median Euclidean distance between predicted and true animal position (cm).

*CCA.* CCA was applied to the first 10 principal components of pairs of population activity matrices. Two comparison types were analyzed: (1) within-session comparisons between the first and second temporal halves of the same recording, and (2) across-session comparisons between all pairs of sessions from the same animal. For each pairwise comparison, only spatial bins visited in both datasets were retained, ensuring that the two matrices were aligned on the same set of shared spatial locations before CCA was performed. CCA identifies linear transformations of the two 10-dimensional PCA spaces that maximize their mutual correlation, generating canonical variates (spatial bins x canonical components) and canonical correlation coefficients (r_1_, r_2_, …) as a measure of representational similarity. A linear regression decoder was then trained on mean-centered neural activity projected onto the first 10 canonical components, following the same two comparison types (within-session and across-session). For each dataset pair, the decoder was trained on one dataset and evaluated on both a held-out 20% test split (within-dataset performance) and the second dataset (cross-dataset performance). For across-session comparisons, decoding was performed in both directions (from dataset 1 to dataset 2 and vice versa), whereas within-session comparisons were evaluated in a single direction (from first half to second half). Two conditions were compared: (1) CCA-aligned decoding using canonical variates after CCA transformation, and (2) unaligned decoding using raw principal components without CCA transformation. Decoding accuracy was quantified as the median Euclidean distance between predicted and true animal position (cm). To establish a lower bound for CCA-based alignment in the absence of spatially organized activity, shuffled control versions of the population activity matrices were generated by randomly permuting spatial bin labels while preserving the population vectors assigned to each bin (each session was permuted individually and differently). Bin averaging already removes temporal structure by collapsing neural activity into spatial-bin-wise population responses prior to PCA and CCA, and the additional shuffle disrupts spatial tuning while preserving neuron-to-neuron covariance. All PCA, CCA and decoding analyses were repeated on the shuffled matrices.

### Tissue processing and histology

At the end of the experiment, mice were transcardially perfused with 4% PFA (Sigma Aldrich) in PBS. Deep anesthesia was induced with isoflurane (5% in 0.5 l/min flow) and intraperitoneal injection of Ketamine/Xylazine (Ketanarkon, 250 mg/kg, Streuli, 2.5 m/kg, Streuli) before the perfusion. Brains were extracted and kept overnight at +4°C in PFA, which was then replaced for a 30% sucrose (Sigma Aldrich) in PBS solution. Brains were frozen and cut into 50 μm sections on a cryostat and kept in PBS. They were then mounted on glass slides and protected by a coverslip with DAPI mounting medium (Sigma Aldrich). Coverslip edges were sealed with nail polish 24 hours after mounting and stored at +4°C in darkness until imaging.

### Plotting and statistics

All statistical analysis were conducted using GraphPad Prism (9.4.1) or Python (version 3.8.12, Python). Results are expressed as the mean ± standard deviation unless otherwise indicated. Box plots show the median (central line), interquartile range (box), and minimum and maximum values (whiskers), and individual data points are shown in each figure unless otherwise indicated. *P* values (*p<0.05, **p<0.01, ***p<0.001) and sample sizes are indicated in the legends (population statistics) and throughout the figures. For quality-control analysis, per session contrasts are reported irrespective of the omnibus test, as the informative result is the absence of differences, otherwise they are only reported when the omnibus identifies a significant group difference. Plots were generated in GraphPad, Matlab, or Python, and subsequently refined in Illustrator (Illustrator 2022 and 2025, Adobe). No statistical tests were used to predetermine sample sizes, but they are similar to the ones reported in previous publications, and post hoc effect sizes are provided when applicable.

**Table S1:**
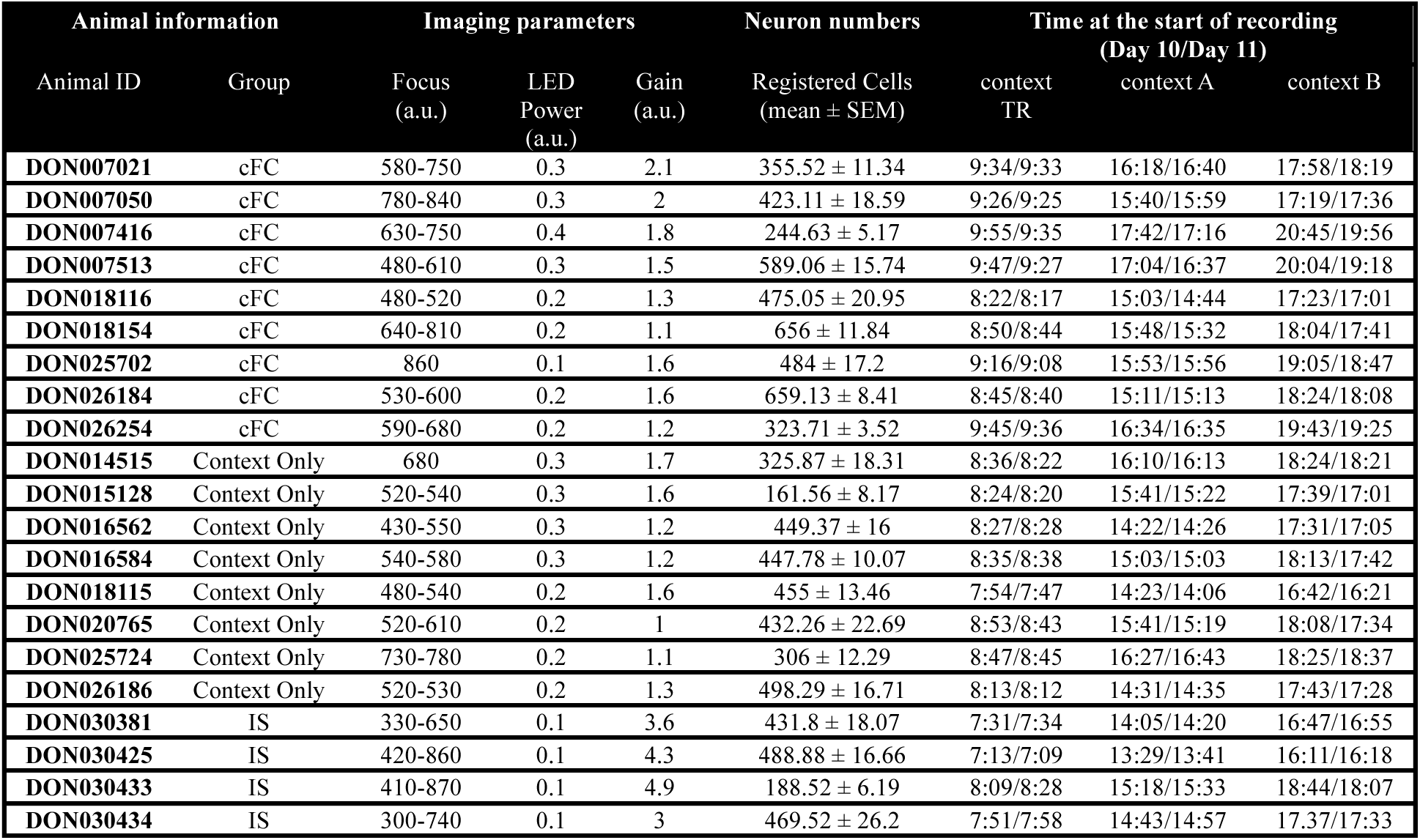
Imaging parameters, number of recorded neurons and recording times for time-lapse calcium imaging of CA3 neurons. Imaging parameters for Fear memory (cFC. n = 9), Context-only (Context Only, n = 8) and Immediate shock (IS, n = 4) mice were set individually during the first imaging session and remained largely constant throughout the experiment. The number of recorded neurons varied from animal to animal. Recording times (hour:min) in each context correspond to Days 10 and 11 and are reported as Day 10/Day 11.

| Animal information |  | Imaging parameters |  |  | Neuron numbers | Time at the start of recording<br>(Day 10/Day 11) |  |  |
| --- | --- | --- | --- | --- | --- | --- | --- | --- |
| Animal ID | Group | Focus<br>(a.u.) | LED<br>Power<br>(a.u.) | Gain<br>(a.u.) | Registered Cells<br>(mean ± SEM) | context<br>TR | context A | context B |
| DON007021 | cFC | 580-750 | 0.3 | 2.1 | 355.52 ± 11.34 | 9:34/9:33 | 16:18/16:40 | 17:58/18:19 |
| DON007050 | cFC | 780-840 | 0.3 | 2 | 423.11 ± 18.59 | 9:26/9:25 | 15:40/15:59 | 17:19/17:36 |
| DON007416 | cFC | 630-750 | 0.4 | 1.8 | 244.63 ± 5.17 | 9:55/9:35 | 17:42/17:16 | 20:45/19:56 |
| DON007513 | cFC | 480-610 | 0.3 | 1.5 | 589.06 ± 15.74 | 9:47/9:27 | 17:04/16:37 | 20:04/19:18 |
| DON018116 | cFC | 480-520 | 0.2 | 1.3 | 475.05 ± 20.95 | 8:22/8:17 | 15:03/14:44 | 17:23/17:01 |
| DON018154 | cFC | 640-810 | 0.2 | 1.1 | 656 ± 11.84 | 8:50/8:44 | 15:48/15:32 | 18:04/17:41 |
| DON025702 | cFC | 860 | 0.1 | 1.6 | 484 ± 17.2 | 9:16/9:08 | 15:53/15:56 | 19:05/18:47 |
| DON026184 | cFC | 530-600 | 0.2 | 1.6 | 659.13 ± 8.41 | 8:45/8:40 | 15:11/15:13 | 18:24/18:08 |
| DON026254 | cFC | 590-680 | 0.2 | 1.2 | 323.71 ± 3.52 | 9:45/9:36 | 16:34/16:35 | 19:43/19:25 |
| DON014515 | Context Only | 680 | 0.3 | 1.7 | 325.87 ± 18.31 | 8:36/8:22 | 16:10/16:13 | 18:24/18:21 |
| DON015128 | Context Only | 520-540 | 0.3 | 1.6 | 161.56 ± 8.17 | 8:24/8:20 | 15:41/15:22 | 17:39/17:01 |
| DON016562 | Context Only | 430-550 | 0.3 | 1.2 | 449.37 ± 16 | 8:27/8:28 | 14:22/14:26 | 17:31/17:05 |
| DON016584 | Context Only | 540-580 | 0.3 | 1.2 | 447.78 ± 10.07 | 8:35/8:38 | 15:03/15:03 | 18:13/17:42 |
| DON018115 | Context Only | 480-540 | 0.2 | 1.6 | 455 ± 13.46 | 7:54/7:47 | 14:23/14:06 | 16:42/16:21 |
| DON020765 | Context Only | 520-610 | 0.2 | 1 | 432.26 ± 22.69 | 8:53/8:43 | 15:41/15:19 | 18:08/17:34 |
| DON025724 | Context Only | 730-780 | 0.2 | 1.1 | 306 ± 12.29 | 8:47/8:45 | 16:27/16:43 | 18:25/18:37 |
| DON026186 | Context Only | 520-530 | 0.2 | 1.3 | 498.29 ± 16.71 | 8:13/8:12 | 14:31/14:35 | 17:43/17:28 |
| DON030381 | IS | 330-650 | 0.1 | 3.6 | 431.8 ± 18.07 | 7:31/7:34 | 14:05/14:20 | 16:47/16:55 |
| DON030425 | IS | 420-860 | 0.1 | 4.3 | 488.88 ± 16.66 | 7:13/7:09 | 13:29/13:41 | 16:11/16:18 |
| DON030433 | IS | 410-870 | 0.1 | 4.9 | 188.52 ± 6.19 | 8:09/8:28 | 15:18/15:33 | 18:44/18:07 |
| DON030434 | IS | 300-740 | 0.1 | 3 | 469.52 ± 26.2 | 7:51/7:58 | 14:43/14:57 | 17:37/17:33 |

